# Inhibition of the DENV2 and ZIKV RNA polymerases by Galidesivir triphosphate measured using a continuous fluorescence assay

**DOI:** 10.1101/2022.12.20.521302

**Authors:** Sandesh Deshpande, Wenjuan Huo, Rinu Shrestha, Kevin Sparrow, Gary B. Evans, Lawrence D. Harris, Richard L. Kingston, Esther M. M. Bulloch

## Abstract

Millions of people are infected by the Dengue and Zika viruses each year, which can result in serious illness, permanent disability or death. There are currently no FDA-approved antivirals for treating infection by these viruses. Galidesivir is an adenosine nucleoside analog which can attenuate flavivirus replication in cell-based and animal models of infection. Galidesivir is converted to the triphosphorylated form by host kinases, and subsequently incorporated into viral RNA by viral RNA-dependent RNA polymerases, leading to the termination of RNA synthesis via an unknown mechanism. Here we report the direct *in vitro* testing of the effects of Galidesivir triphosphate on RNA synthesis by the polymerases of Dengue-2 and Zika virus. Galidesivir triphosphate was chemically synthesized and inhibition of RNA synthesis followed using a continuous fluorescence-based assay. Galidesivir triphosphate was equipotent against the polymerase activity of Dengue-2 and Zika, with IC_50_ values of 42 ± 12 μM and 47 ± 5 μM, respectively. This modest potency *in vitro* is consistent with results previously obtained in cell-based antiviral assays and suggests that the binding affinity for Galidesivir triphosphate is similar to the natural ATP substrate that it closely mimics. The inhibition assay we have developed will allow the rapid screening of Galidesivir and related compounds against other flavivirus polymerases, and the availability of Galidesivir triphosphate will allow detailed analysis of its mechanism of action.

**Highlights:**

- Galidesivir triphosphate was chemically synthesized.
- A continuous assay detecting double-stranded RNA formation was optimized for polymerase inhibition studies.
- Galidesivir triphosphate has moderate potency against DENV2 and ZIKA polymerase activity.
- The availability of Galidesivir triphosphate will facilitate study of its mechanism of action.

## 1. Introduction

Flaviviruses have affected human health for centuries, and their impacts are expanding due to both anthropogenic climate change and the increased mobility of human populations (Beltz, 2021; Pierson & Diamond, 2020). Flaviviruses (Family *Flaviridae*, Genus *Flavivirus*) are positive-sense, enveloped RNA viruses, which are usually spread between vertebrate hosts by arthropod vectors, such as mosquitos and ticks, and are most prevalent in tropical and sub-tropical regions. Important endemic and emerging flaviviruses include Dengue virus (DENV, serotypes 1 to 4), Japanese encephalitis virus (JEV), Tick-borne encephalitis virus (TBEV), West Nile virus (WNV) and Zika virus (ZIKV). When humans are infected with a flavivirus the resulting disease is sometimes mild, but potentially fatal hemorrhagic fever and neurological disorders can also result. In addition, unique clinical manifestations have been reported, such as fetal microcephaly caused by the Zika Virus (Mlakar et al., 2016).

Vaccines are currently available for JEV, TBEV, YFV and DENV, and clinical trials are being undertaken for ZIKV and WNV vaccines (Zhou et al., 2021; Ulbert, 2019). However, existing flavivirus vaccines are of variable efficacy, and their development has not always been straightforward, because of the complex immunological response to infection by different flaviviruses, and the presence of multiple endemic flaviviruses in tropical regions (Fischer et al., 2020; Sridhar et al., 2018). Despite more than 60 years of research, there are currently no FDA-approved antivirals to treat flavivirus infection, and there is an urgent need for these to be developed (Komarasamy et al., 2022).

One approach that has been successful in the treatment of other viral infections is the use of nucleoside analogs to inhibit virally-directed nucleic acid synthesis (Seley-Radtke & Yates, 2018; Yates & Seley-Radtke, 2019; Kataev & Garifullin, 2021). In this study we investigate the inhibitory effects of Galidesivir (Figure 1), an adenosine analog with broad spectrum activity against RNA viruses, including flaviviruses (Warren et al., 2014). Galidesivir is an iminoribitol C-nucleoside, which is converted to the triphosphorylated form by host kinases, and then incorporated into viral RNA by viral RNA-dependent RNA polymerases (RdRps). Primer extension assays with purified Hepatitis C virus (HCV) RdRp indicate that Galidesivir acts as a non-obligate chain terminator, though the mechanism remains undefined. In the case of HCV, two further nucleotides are added subsequent to Galidesivir monophosphate incorporation, and then RNA synthesis ceases (Warren et al., 2014).

**Figure 1:**
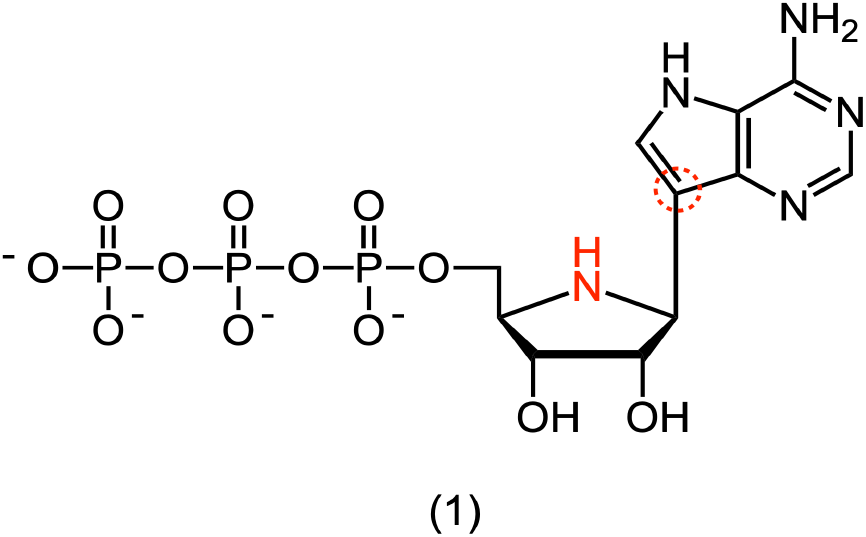
Galidesivir triphosphate (Gal-TP) is produced by the phosphorylation of Galidesivir by kinases *in vivo*. Red highlighting indicates sites of modification relative to adenosine triphosphate.

In cell based assays, Galidesivir inhibits the replication of the flaviviruses YFV, JEV, DENV2, WNV and TBEV with half maximal effective concentrations (EC_50_) in the low micromolar range (Warren et al., 2014; Julander et al., 2017; Eyer et al., 2017). Galidesivir is also effective against YFV, TBEV and ZIKV in rodent models of infection (Julander et al., 2014; Julander et al., 2017; Eyer et al., 2019). In a non-human primate model, Galidesivir provided post-exposure protection to ZIKV infection (Lim et al., 2020). This study also found that Galidesivir was not associated with reproductive toxicity at dosages of ≤75 mg/kg/day in pregnant rats and ≤25 mg/kg/day in pregnant rabbits, and that the antiviral crossed the placenta (Lim et al., 2020). In two phase I clinical trials of Galidesivir no serious adverse effects were associated with administration of the antiviral and it was reported to be well-tolerated (Mathis et al., 2022). Galidesivir is therefore a promising lead compound for further development of therapeutics targeting flaviviruses, by inhibiting their RdRp activity.

The flavivirus RdRp is part of the multifunctional NS5 protein, fused to the C-terminus of a methyltransferase domain that is involved in RNA capping (Egloff et al., 2002). Although the RdRp is active in the absence of the methyltransferase domain, full length NS5 is required for efficient initiation and elongation (Wang et al., 2012; Wu et al., 2015; Potisopon et al., 2014). Flaviviral RdRps are capable of both primed and *de novo* RNA synthesis, with the *de novo* process involving the relatively slow formation of a dinucleotide primer from two incoming mononucleotide triphosphates (Ackermann & Padmanabhan, 2001; Nomaguchi et al., 2003; Surana et al., 2014; Selisko et al., 2012; Potisopon et al., 2014).

In all prior work, the effects of Galidesivir on flaviviral replication have been assessed in cell-based antiviral assays. These effects reflect both the efficiency with which Galidesivir is converted into the active triphosphate form by host kinases, as well as inhibition of the viral RdRp (Warren et al., 2014; Julander et al., 2017; Eyer et al., 2017). In this study, we chemically synthesized Galidesivir triphosphate (Gal-TP) and measured its potency in inhibiting DENV2 and ZIKV NS5 RdRps *in vitro*. RdRp activity was followed using an adaptation of a continuous, SYTO 9 fluorescence-based assay for double stranded RNA formation (Sáez-Álvarez et al., 2019). Dose response curves for Gal-TP against the RNA elongation activity of the flavivirus RdRps were determined using dinucleotide-primed reactions. The results give insight into the relative potency of Galidesivir against flaviviral RdRps, and the methods developed provide a platform for testing of other nucleoside analogs.

## 2. Methods

### 2.1 Synthesis of Galidesivir triphosphate

Full details of the synthesis and characterisation of Galidesivir triphosphate are given in the Supplementary Information.

### 2.2 Protein production and purification

Genes encoding the full length NS5 proteins from DENV2 (GenBank accession no. <u>**NC001474**</u>, UniProt accession no. <u>**P29990**</u>), and ZIKV (GenBank accession no. <u>**NC012532**</u>, UniProt accession no. <u>**Q32ZE1**</u>) were commercially synthesized and cloned into plasmid pUC57 by GenScript Biotech. NS5 coding sequences were subsequently transferred into expression plasmid pET-15b (Novagen) using standard molecular biology procedures (Supplementary Figure S1). These expression constructs appended a N-terminal His_6_-tag to the NS5 protein, with an interleaving Tobacco Etch Virus (TEV) protease cleavage site to facilitate tag removal.

Production of NS5 proteins was carried out using an autoinduction protocol (Studier, 2005), in *E. coli* Rosetta™ 2 (DE3) cells (Novagen). Chemically competent Rosetta™ 2 (DE3) were transformed with the relevant NS5 expression plasmid. Precultures (10 mL) inoculated from a single colony of the transformed bacteria were grown overnight at 30 °C in non-inducing minimal medium, MDG, supplemented with appropriate antibiotics. Overexpression was carried out by inoculating 5 mL of the preculture into 500 mL of auto-inducing medium, ZYM-5052 in baffled flasks. Cultures were maintained at 37 °C for 2.5 h and then 18 °C for 20 h, with shaking at 160 rpm throughout. Bacteria were pelleted by centrifugation and stored at -20 °C until further processed.

To purify NS5 proteins, cell pellets were thawed and the bacteria re-suspended in lysis buffer (10 mL per gram of cell pellet) comprising 50 mM potassium phosphate, pH 7.5, 300 mM NaCl, 10 % glycerol and 5 mM β-mercaptoethanol. HEWL Lysozyme (50 µg/mL, Sigma Aldrich), Bovine pancreatic DNase I (10 µg/mL, Sigma) and an EDTA-free protease inhibitor cocktail tablet (cOmplete™, Roche Diagnostics) were added to the cell suspension before incubating on ice for 15 min. Cells were lysed using a continuous cell disruptor (M-110P Microfluidizer, Constant Systems Ltd). The lysate was centrifuged at 20,000 *xg* for 30 min at 4 °C to pellet cellular debris, and the supernatant was loaded onto TALON® IMAC resin (Takara Bio) pre-equilibrated with the lysis buffer, under gravity flow. NS5 proteins were eluted from the IMAC resin using lysis buffer supplemented with 300 mM imidazole. The eluted protein was dialysed overnight at 4 °C against the lysis buffer, adding recombinant Tobacco Etch Virus (TEV) protease to a final concentration of 0.1 mg/mL during the dialysis step.

The protein was subsequently passed over a fresh bed of TALON® IMAC resin, pre-equilibrated with lysis buffer, to remove uncleaved NS5 protein that retained the His_6_-tag. The column flow through was concentrated to a final volume of ~500 µL and subject to size exclusion chromatography (SEC) using a HiLoad® 16/600 column packed with Superdex® 200 media (Cytiva). The SEC column was pre-equilibrated with 20 mM MOPS/KOH pH 7.5, 300 mM NaCl, 10 %(v/v) glycerol and 1 mM TCEP-HCl, and the protein was eluted isocratically in this buffer. Fractions containing the purified NS5 protein were pooled and concentrated to 0.2-1 mg/mL. The protein was aliquoted; flash frozen in liquid nitrogen; and stored at -80 °C until subsequent use.

### 2.3 Size exclusion chromatography coupled to multi-angle laser light scattering (SEC-MALLS)

SEC-MALLS analysis of the purified NS5 proteins was performed using a Superdex™ 200 10/300 column (Cytiva) connected to an Ultimate 3000 HPLC system (Thermo Fisher Scientific) equipped with in-line MALLS (Polymer Standards Service SLD7000) and differential refractive index (Shodex RI-101) detectors. The SEC column was equilibrated with 20 mM MOPS/KOH pH 7.5, 300 mM NaCl, 10 %(v/v) glycerol, 3 mM sodium azide and 1 mM TCEP-HCl. NS5 proteins were loaded at concentrations of 0.45 and 0.9 mg/mL, and eluted isocratically. The mass averaged molar mass of the eluted proteins was determined from the refractive index and light scattering measurements using the PSS WinGPC UniChrom 8.1 software, under assumption of Rayleigh scattering. A constant refractive index increment of 0.186 was used to estimate all protein concentrations, with the MALLS detector calibrated using a bovine serum albumin solution.

### 2.4 SYTO 9 fluorescence-based RdRp assays

A fluorescence-based assay adapted from a study by Sáez-Álvarez and coworkers was used to measure RdRp activity (Sáez-Álvarez et al., 2019). All reagents and buffers were prepared in nuclease free water. The assays were carried out in 96-well black, flat-bottom microplates (Greiner). Assay mixtures (100 µL) were prepared as detailed below, and the enzymatically catalyzed reactions typically initiated by addition of ATP. Fluorescence was recorded for 1 h on a Biotek Synergy HTX plate reader (Agilent) with excitation and emission filters of wavelength 485/20 and 528/20 nm, respectively.

The background fluorescence signal for each reaction was estimated from a matched negative control, from which only ATP was omitted. The initial rates of fluorescence increase were measured by linear fitting of background-corrected progress curves, excluding the initial region of the curve if there was any apparent lag phase (duration 500 – 1000 s). Technical duplicates or triplicates were collected for each assay condition. Data analysis, including background subtraction, linear fitting to determine initial rates, and non-linear model fitting to estimate apparent kinetic constants and IC_50_ values, was carried out using GraphPad Prism version 9.0.

### 2.5 Optimization of de novo reaction conditions for DENV2 NS5

Unless otherwise stated, the assay mixtures for initial optimization of the *de novo* reaction (section 3.2) contained 50 mM MOPS/KOH pH 7.5, 2.5 mM MnCl_2_, 2.5 µM SYTO 9 (Thermo Fisher), 40 µg/mL polyuridylic acid (poly(U), Sigma), 200 nM DENV2 NS5, and were initiated by the addition of 500 µM ATP (New England Biolabs).

### 2.6 Optimization of primed reaction conditions for DENV2 NS5

Unless otherwise stated, the assay mixtures for optimization of the primed reactions (section 3.3) contained 50 mM MOPS/KOH pH 7.5, 1.5 mM MnCl_2_, 3 µM SYTO 9, 40 µg/mL poly(U), 200 nM DENV2 NS5 and RNA primer. Reactions were initiated with ATP, as detailed below.

For establishing the most efficiently incorporated primer, non-5’-phosphorylated poly(A) primers varying in length from A2 to A10 were obtained commercially. A2 (ApA) RNA dinucleotide primer was purchased from Jena Bioscience. A3 to A10 primers were synthesized by GenScript Biotech. Comparative assays were run as described above with 10 μM RNA primer (A2 to A10) present, or no primer for the *de novo* reaction. Reactions were initiated by addition of 10 μM ATP.

Experiments to measure the effect of A2 primer concentration on the apparent *K*_M_ and *V*_max_ (section 3.4) were performed by measuring the initial rate at ATP concentrations ranging from 0 - 1500 μM and at A2 concentrations from 0 to 40 μM. ATP concentrations did not exceed 1.5 mM to prevent depletion of free Mn^2+^, which would result in large rate variations (see Section 3.3.4).

The ATP-dependence data were fitted to the Michaelis-Menten equation:

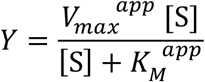

where Y is the reaction rate, [S] is the substrate concentration, *V*_max_^app^ is the maximal reaction rate and *K*_M_^app^ is the substrate concentration at half the maximal reaction rate.

The apparent turnover number, *k*_cat_^app^, is defined as:

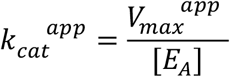

where is [E_A_] is the concentration of NS5 enzyme active sites.

For ATP-dependence experiments the rate of fluorescence change was converted to the rate of nucleotide incorporation using a standard curve generated with the double-stranded RNA poly(A)-poly(U) (Sigma Aldrich), which is the reaction product detected by SYTO 9. The poly(A)-poly(U) was added at 0 to 8 µg/mL concentrations to assay mixtures containing the same components as the reaction mixtures, with the exception of ATP. The fluorescence of the standard solutions was read after a 2000 s incubation in the plate reader to allow for signal stabilisation. Fluorescence was plotted against the effective concentration of A:U base pairs in each standard sample, and a linear fit used to determine the fluorescence change per nucleotide incorporated (example standard curve presented in Supplementary Figure S5). After converting raw reaction rates to nucleotides incorporated per minute, rates were normalized by the enzyme concentration, giving final reaction rates in terms of nucleotides incorporated per NS5 molecule per minute (min^-1^).

### 2.7 Inhibition assays with nucleoside triphosphate analogs

Assay mixtures for measuring inhibitory effects of Gal-TP on DENV2 and ZIKV NS5 contained 50 mM MOPS/KOH, pH 7.5, 2.0 mM MnCl_2_, 3 µM SYTO 9, 40 µg/mL poly(U), 20 µM A2 primer, 20 µM ATP, 0 to 2 mM Gal-TP, and were initiated by addition of 200 nM DENV2 NS5. Gal-TP did not exceed 2 mM to prevent formation of an insoluble complex with Mn^2+^ (data not shown). Inhibition assays with 3’-deoxyATP were carried in the same fashion but with 0 to 200 µM 3’-deoxyATP in place of Gal-TP.

Reaction rates were measured and expressed as a percentage of the uninhibited rate for each NS5. The data were fit to an empirical four-parameter logistic model (Gottschalk & Dunn, 2005):

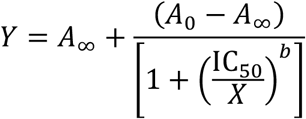

where *Y* is the reaction rate, *X* is the inhibitor concentration, *A*_*0*_ and *A*_*∞*_ are the asymptotic values for the reaction rate at very low and very high inhibitor concentrations, respectively, IC_50_ is the half maximal inhibitory concentration, and *b* is a slope parameter.

## 3. Results and discussion

### 3.1 Selection of flaviviral NS5 proteins for study

NS5 proteins from Dengue-2 (DENV2) and Zika virus (ZIKV) were selected for testing of the inhibitory effects of Gal-TP due to the current global impacts of these viruses. NS5 is the most highly conserved flaviviral protein, with near complete conservation of residues involved in substrate-binding and catalysis within the RdRp domain (da Fonseca et al., 2017). DENV2 and ZIKV NS5 have 66 % sequence identity and 85% similarity when aligned using Clustal Omega (Supplementary Figure S2) (McWilliam et al., 2013).

### 3.2 Production of full length DENV2 and ZIKV NS5 proteins

DENV2 and ZIKV NS5 proteins were heterologously produced in *E. coli* with an N-terminal His_6_-tag and purified using a standardized protocol as described in the methods (Section 2.2). This protocol included removal of the His_6_-tag, the presence of which is reported to have a deleterious effect on DENV2 RdRp activity (Kamkaew & Chimnaronk, 2015).

While DENV2 and ZIKV NS5 were purified free of significant contaminants, as assessed by SDS PAGE (Supplementary Figure S3), some proteolytic degradation was apparent, particularly for Zika NS5. As confirmed by mass spectrometry (data not shown), the major site of cleavage was the flexible linker between the methyltransferase and RdRp domains.

Analysis of purified DENV2 and ZIKV NS5 by size exclusion chromatography coupled to multi-angle laser light scattering (SEC-MALLS) indicated that both proteins are predominantly monomeric in solution at concentrations less than 1 mg/mL (data not shown).

### 3.3 Development of a NS5 RdRp activity assay for inhibition studies

#### 3.3.1 Adaption of a SYTO 9-based assay for flavivirus NS5 RdRp activity

In this study a fluorescence-based continuous assay developed by Sáez-Álvarez *et al*. (Sáez-Álvarez et al., 2019), was adapted to enable testing of Gal-TP against flavivirus NS5 RdRp activity. The underlying principle of the assay is enhancement in fluorescence of SYTO 9 upon binding to double-stranded RNA (dsRNA) in preference to single-stranded RNA (ssRNA). SYTO 9 is a cyanine dye (Deligeorgiev et al., 2009), however the exact structure is proprietary and its nucleic acid binding mechanism is not fully characterized. Importantly, unlike many other dsRNA-binding dyes, SYTO 9 is reported to not significantly inhibit RdRp activity, and hence is suitable for use in a continuous assay (Sáez-Álvarez et al., 2019).

DENV2 NS5 was used for the assay optimisation experiments as it was obtained at higher yield than ZIKV NS5 (data not shown). The standardized assay was carried out at pH 7.5, consistent with previously reported activity optima (Yu et al., 2007; Wu et al., 2015; Grun & Brinton, 1986; Smith et al., 2015; Shimizu et al., 2019), and at a temperature of 25 µC, in order to minimize temperature perturbations when 96 well plates were transiently moved into ambient temperature to initiate reactions.

An example of a *de novo* reaction progress curve for DENV2 NS5 following initiation of the reaction by addition of ATP is shown in Figure 2A, together with a matched negative control (ATP omitted) used to estimate the background fluorescence signal. Initial rates of fluorescence change were determined by linear fit of background-corrected progress curves, excluding an apparent lag phase evident in the first 500-1000s of the reaction.

**Figure 2:**
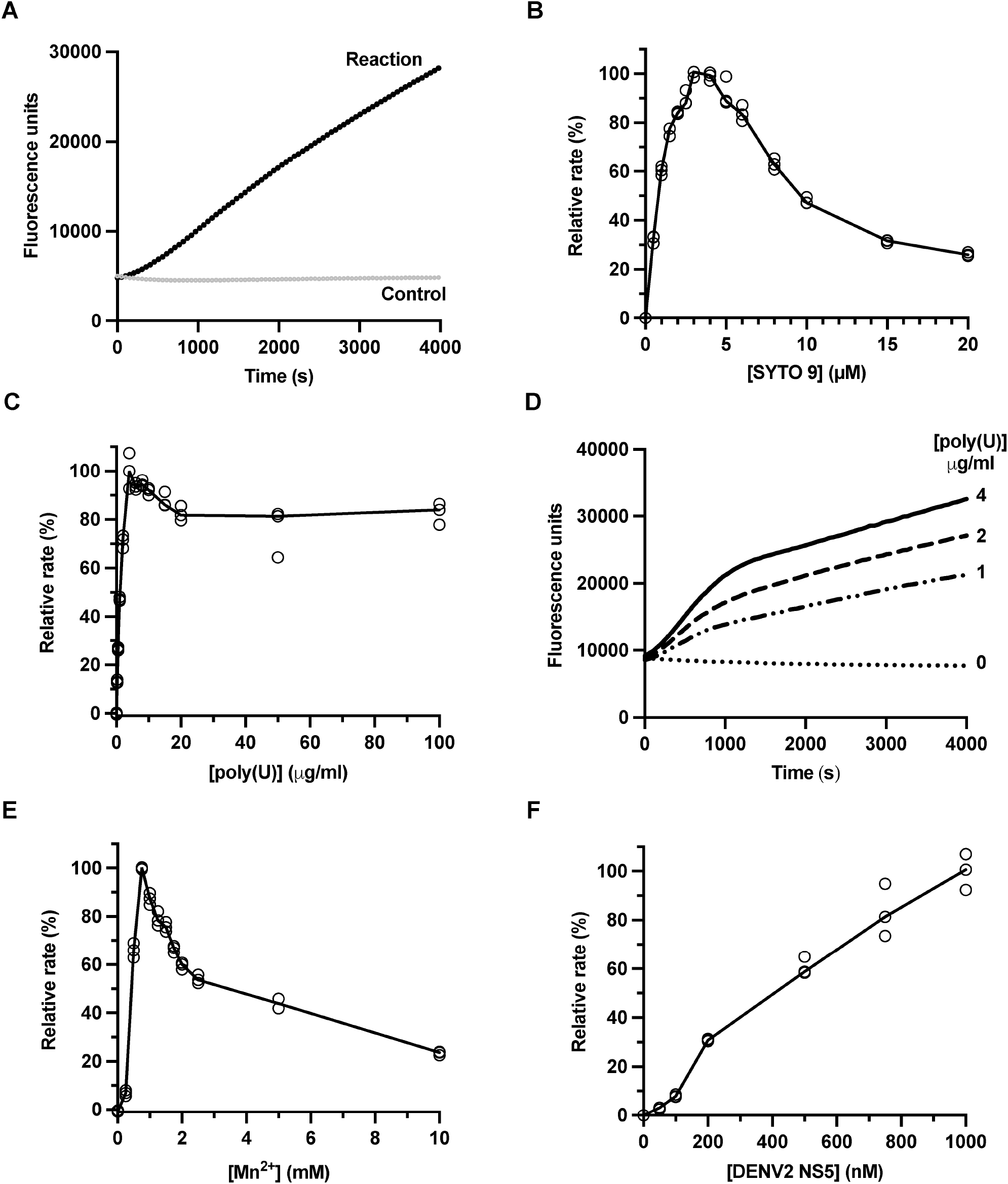
Optimisation of the SYTO 9 fluorescence-based RdRp assay using DENV2 NS5. Unless otherwise stated reactions contained 200 nM DENV2 NS5, 2.5 mM MnCl_2_, 40 μg/mL poly(U), 2.5 μM SYTO 9 and were initiated by addition of 0.5 mM ATP. (Panel A) Time-dependent fluorescence (excitation 485/emission 528 nm) from a DENV2 NS5 catalyzed reaction, and a matched negative control without ATP. (Panels B to F) Dependence of measured rate on concentrations of: SYTO 9 dye (Panel B), poly(U) (Panels C and D*), Mn^2+^ (Panel E) and DENV2 NS5 (Panel F). Measured rates in fluorescence units per second are given as a percentage relative to the maximum rate for that experiment. In panels B, C, E and F the hollow circles show the results of individual technical replicates, while the solid lines are drawn through the median of each set of technical replicates. * Panel C shows measured rate as a function of poly(U) concentration while panel D shows the biphasic reaction progress curves characteristically observed with poly(U) concentrations under 10 μg/mL.

A series of experiments were carried out to examine the effect of SYTO 9, poly(U) template, divalent metal ion (Mn^2+^ or Mg^2+^) and enzyme concentration on the measured rate for DENV2 NS5.

#### 3.3.2 SYTO 9 dye concentration

Varying the SYTO 9 concentration in the assay from 0.5 to 20 μM resulted in a rapid increase in the measured rate of fluorescence change up to a maximum of 3 μM SYTO 9 (Figure 2B). Above this concentration there was an apparent decrease in the rate of fluorescence change, likely due to an inner-filter effect at higher SYTO 9 concentrations. A concentration of 3 μM SYTO 9 was selected for the subsequent standardised inhibition assays.

#### 3.3.3 RNA template

Since Galidesivir is an adenosine analog, a homopolymeric poly(U) template was selected for development of an Gal-TP inhibition assay. The initial reaction rate increased steeply with increasing poly(U) concentration with a maximum at 4 - 10 μg/mL (Figure 2C), similar to the maximum reported previously for ZIKV RdRp domain (Sáez-Álvarez et al., 2019). Although there was a slight suppression of rate at poly(U) concentrations above 10 μg/mL (Figure 2C), a concentration of 40 μg/mL was selected for the subsequent assays. This is because the progress curves at low poly(U) concentrations (< 10 μg/mL) were clearly biphasic (Figure 2D), with an initial rapid phase followed by an extended slower phase. The use of elevated poly(U) concentrations afforded a longer pseudo-linear phase for measurement of initial reaction rates.

#### 3.3.4 Metal ion dependence

Although, both the divalent metal ions Mn^2+^ and Mg^2+^ support flavivirus RdRp activity *in vitro*, the presence of Mn^2+^ has been shown to result in significant rate enhancement (Selisko et al., 2006; Kim et al., 2007) However, activation of viral RdRps by Mn^2+^ is associated with increased misincorporation rates and copy-back activity. (Arnold et al., 2004; Liu et al., 2018; Chen et al., 2022). For DENV2 NS5, non-templated formation of dinucleotide RNA primers occurs *in vitro* in the presence of Mn^2+^ ions but not Mg^2+^ ions (Selisko et al., 2012). Studies of bacteriophage χΠ6 RdRp suggest that Mn^2+^ binding to this enzyme provides increased structural flexibility, facilitating the dynamic movements required for catalysis (Poranen et al., 2008).

The measured initial rates for DENV2 NS5 in the SYTO 9 based assay were highest in the presence of Mn^2+^, with a sharp optimum at 0.75 mM Mn^2+^ for the *de novo* reaction with 0.5 mM ATP (Figure 2E). At higher Mn^2+^ concentrations RdRp activity was suppressed. The maximum rate in the presence of Mg^2+^ was two orders of magnitude lower than the rate in the presence of 0.75 mM Mn^2+^ (Supplementary Figure S4).

The free divalent metal ion concentration is highly dependent on the concentration of nucleotide triphosphates, due to the tight association of divalent metal ions and NTPs (Sigel, 1977). Therefore, as detailed in the following sections, the concentration of Mn^2+^ in each assay was varied according to the maximum nucleotide triphosphate concentration used.

#### 3.3.5 NS5 concentration dependence

For a simple monomeric enzyme, the initial reaction rate should be strictly proportional to enzyme concentration. In contrast, for DENV2 NS5 (0 to 1 μM) the observed reaction rate is initially a convex function of enzyme concentration (Figure 2F), with the relationship becoming approximately linear at higher concentrations ([NS5] > 200 nM). The non-linear response at low enzyme concentrations could reflect a linkage between enzyme activity and self-association, as previously proposed (Saw et al., 2015; Saw et al., 2017; Klema et al., 2016; Ferrero et al., 2019). While isolated DENV2 NS5 is predominantly monomeric at the concentrations used in the assay (Section 3.2), NS5 oligomerization could be promoted by the presence of RNA template, primer and/or nucleotide substrates.

### 3.4 Development of a primed assay for DENV2 NS5 RdRp activity

*De novo* RNA synthesis by flavivirus RdRps involves a slow, rate-limiting, initiation phase in which a dinucleotide is formed (Ackermann & Padmanabhan, 2001; Nomaguchi et al., 2003; Surana et al., 2014; Selisko et al., 2012; Potisopon et al., 2014). With the addition of a RNA primer, the initiating step is effectively bypassed and the RdRp can transition to faster processive RNA synthesis. Given the proposed mechanism of action of Gal-TP is termination of processive RNA synthesis (Warren et al., 2014), it is most relevant to measure its inhibitory effect in a primed RNA synthesis assay, rather than a *de novo* assay.

A prior study of primed RNA elongation by DENV2 NS5 showed that a tetranucleotide primer was more efficiently incorporated than either shorter or longer primers, in an assay with a subgenomic RNA as template (Nomaguchi et al., 2003). Similarly, for JEV NS5 the presence of a dinucleotide U2 primer led to higher rates of activity than a U10 primer in assays with a poly(A) template (Wang et al., 2012). Finally, in a study of the distantly related hepatitis C virus NS5B, pGG and pGGG primers were incorporated at higher efficiency than longer primers in assays with complementary heteropolymeric templates (Zhong et al., 2000).

Consequently, we tested the effects of poly(A) primers from two to ten nucleotides in length, on DENV2 NS5 RdRp activity with a poly(U) template (Figure 3). The A3 primer was most efficiently incorporated by DENV2 NS5, with a reaction rate 20-fold higher than in the *de novo* reaction at a primer concentration of 10 μM. The reaction rate in the presence of A2 and A4 primers was in each case 6-fold higher than the *de novo* reaction, while the reaction rate with the A5 primer was only 3-fold higher than the *de novo* reaction. With the use of longer primers (A6 to A10) rate enhancements decreased, and there were also substantial increases in background fluorescence due to annealing of the primers to the poly(U) template (data not shown).

**Figure 3:**
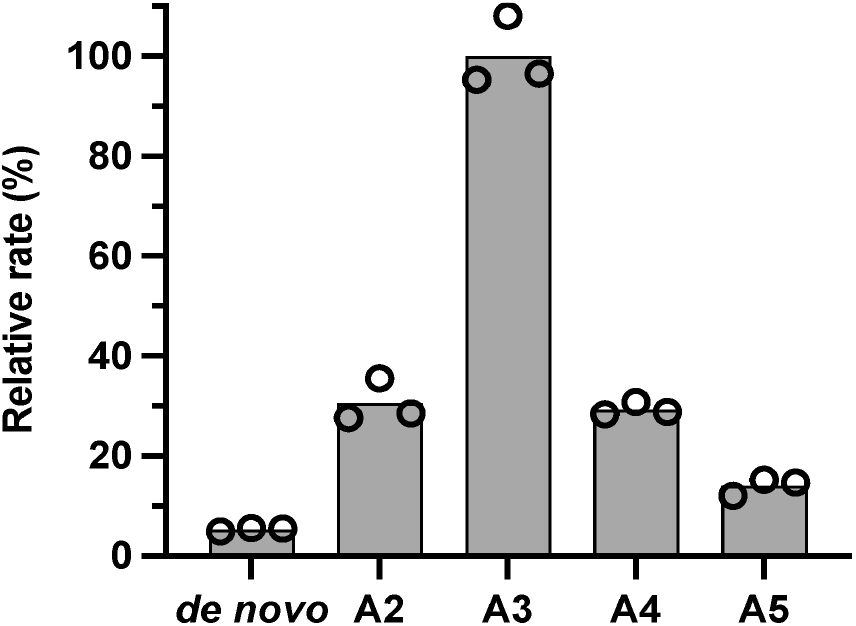
Enhancement of NS5 DENV2 activity by the addition of short poly(A) primers. Reactions contained 200 nM DENV2 NS5, 1.5 mM MnCl_2_, 40 μg/mL poly(U), 3 μM SYTO 9 and were initiated with 10 μM ATP. In addition, reactions contained either no primer (*de novo* reaction) or 10 μM of a non-5’-phosphorylated poly(A) primer. The primers tested were A2 (ApA), A3 (ApApA), A4 (ApApApA) and A5 (ApApApApA). Grey bars represent the mean relative rate of three technical replicates, and hollow circles indicate the individual replicate values.

Although the trinucleotide A3 primer is most efficiently incorporated by DENV2 NS5, this RNA primer must be custom synthesized and the costs are currently prohibitive. Hence, the dinucleotide A2 primer was used for the standardised inhibition assays, as it is readily sourced commercially.

### 3.5 ATP-dependence of DENV2 NS5 RdRp activity in de novo and A2 primed reactions

The ATP-dependence of DENV2 NS5 RdRp activity was measured for both *de novo* and primed reactions with varying concentrations of the A2 RNA primer (Figure 4 and Table 1). For these experiments rates of fluorescence change were converted to a nucleotide incorporation rate though use of poly(A):poly(U) dsRNA standards (Section 2.5). This allowed absolute comparisons of nucleotide incorporation rates in the presence and absence of the primer.

**Figure 4:**
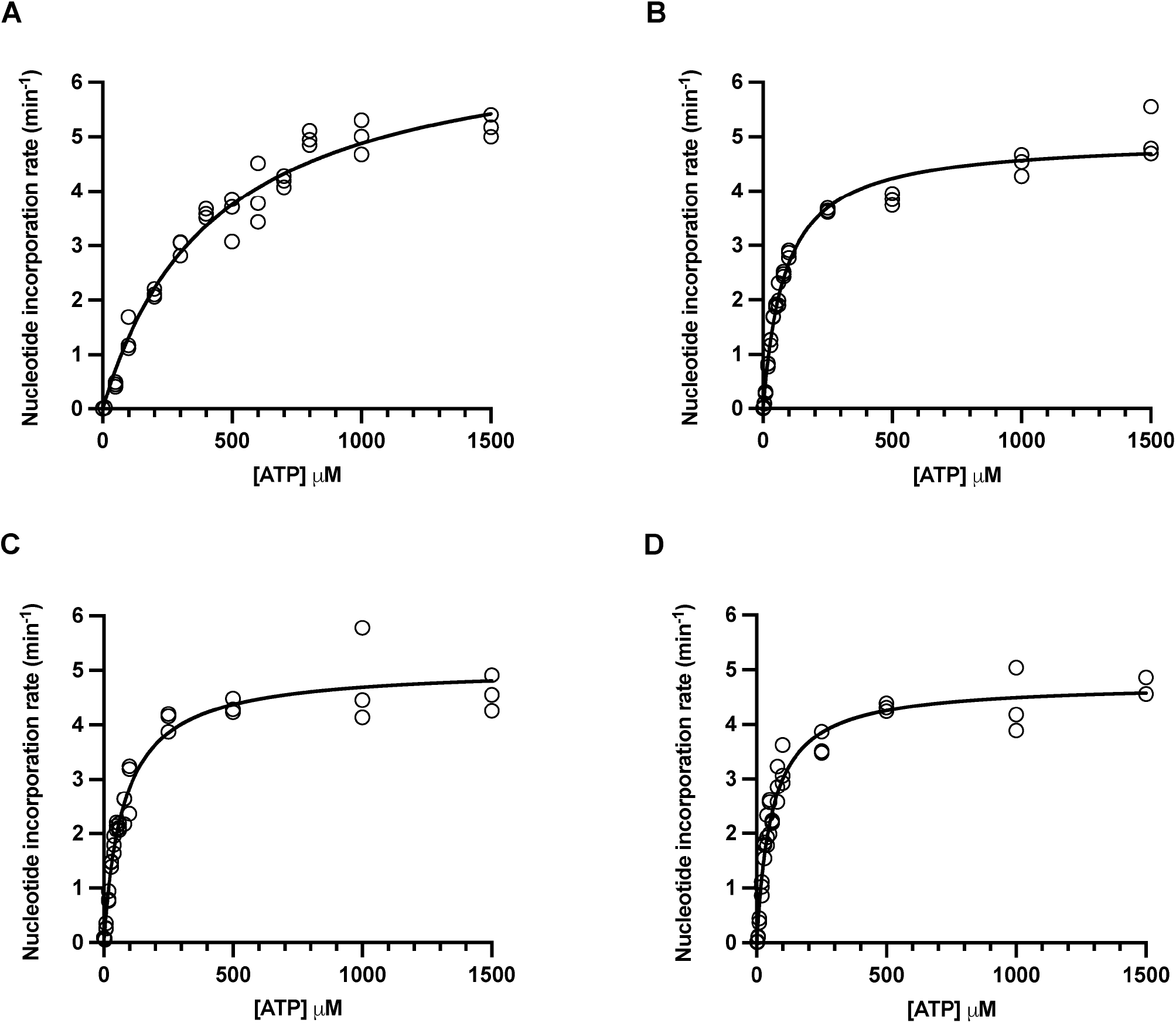
ATP dependence of DENV2 NS5 activity in the absence and presence of an A2 primer. The apparent *K*_M_ for ATP was determined in the absence of primer (*de novo*, Panel A), and in the presence of A2 (ApA-primer at concentrations of 10, 20 and 40 μM (Panels B, C and D respectively). Rates are reported as nucleotides incorporated per molecule of DENV2 NS5 per minute. Reactions contained 200 nM DENV2 NS5, 0 to 40 μM A2 primer, 40 μg/mL poly(U), 2.5 mM MnCl_2_, 3 μM SYTO 9, and were initiated with ATP at concentrations ranging from 0 to 1500 μM. Hollow circles show the experimental measurements, while solid lines show the fit of the data to the Michaelis-Menten equation.

**Table 1:**
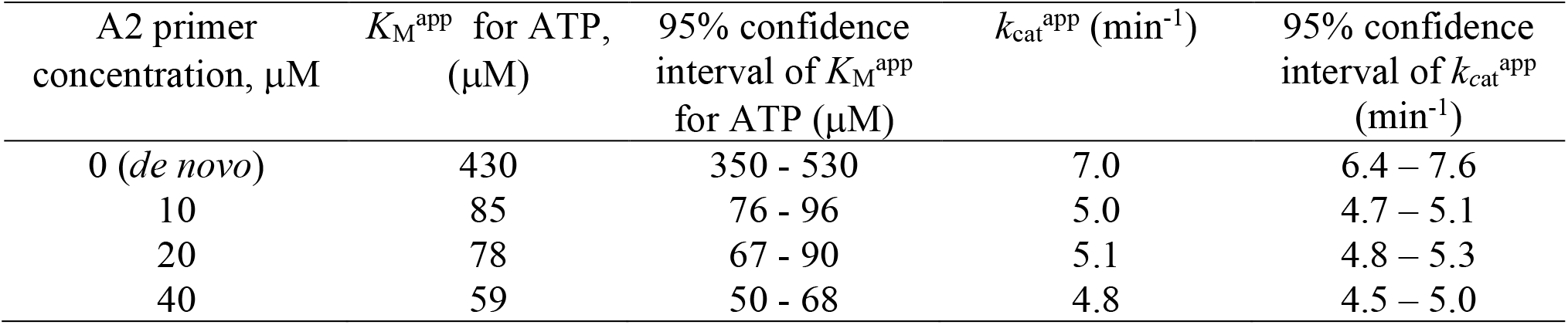
Effect of A2 primer concentration on the apparent kinetic constants for DENV2 NS5.

| A2 primer concentration, $\mu\text{M}$ | $K_M^{\text{app}}$ for ATP, ($\mu\text{M}$) | 95% confidence interval of $K_M^{\text{app}}$ for ATP ($\mu\text{M}$) | $k_{\text{cat}}^{\text{app}}$ ($\text{min}^{-1}$) | 95% confidence interval of $k_{\text{cat}}^{\text{app}}$ ($\text{min}^{-1}$) |
| --- | --- | --- | --- | --- |
| 0 (*de novo*) | 430 | 350 - 530 | 7.0 | 6.4 – 7.6 |
| 10 | 85 | 76 - 96 | 5.0 | 4.7 – 5.1 |
| 20 | 78 | 67 - 90 | 5.1 | 4.8 – 5.3 |
| 40 | 59 | 50 - 68 | 4.8 | 4.5 – 5.0 |

For both *de novo* and primed reactions the ATP-dependence data could be fit to the Michaelis-Menten equation to determine an apparent Michaelis constant *K*_M_ for ATP, and apparent rate constant *k*_cat_ for nucleotide incorporation (Figure 4 and Table 1). It should be noted that the reaction rates measured in the presence of primer result from both primed RNA synthesis and *de novo* initiation. At low ATP concentrations the more efficient A2 primed reaction dominates, and at higher ATP concentrations *de novo* initiation dominates.

In the absence of a primer (*de novo* reaction) the *K*_M_^app^ for ATP was 430 μM (Figure 4A and Table 1). In the presence of 10 μM of A2 primer the *K*_M_^app^(ATP) was 85 μM, a 5-fold decrease relative to the *de novo* reaction (Figure 4B and Table 1). As the A2 primer concentration was further raised, *K*_M_^app^(ATP) incrementally decreased, down to a value of 59 μM with 40 μM of A2 primer (Figure 4C, 4D and Table 1). When the A2 primer was introduced, the apparent rate constant, *k*_cat_^app^, decreased 1.4-fold relative to the *de novo* reaction, however there was no further change with increasing primer concentration (Table 1).

The Michaelis constant for *de novo* RNA synthesis by DENV2 NS5, is consistent with the *K*_M_^app^(ATP) of 560 ± 40 μM previously reported for *de novo* RNA synthesis by the ZIKV RdRp domain using a poly(U) template (Sáez-Álvarez et al., 2019). A lower *K*_M_^app^ (ATP) of 32 ± 1 μM was reported for DENV2 NS5 in *de novo* initiation studies, using the first 20 nucleotides of the DENV2 antigenome as a template and with a Mn^2+^ cofactor (Potisopon et al., 2014). This is likely to reflect more efficient initiation on templates carrying the native CU-3’ sequence, relative to a homopolymeric poly(U) sequence. Sub-micromolar *K*_M_ values have been reported for all NTPs in processive RNA synthesis assays of DENV2 NS5 using long templates corresponding to the native 3’ UTR sequence of the DENV2 genome, and with a Mg^2+^ cofactor (Latour et al., 2010).

For the standardised inhibition assays we set [ATP] = [A2 primer] = 20 μM. At these concentrations the rate of DENV2 NS5-catalysed nucleotide incorporation is approximately ten-fold faster than the *de novo* reaction with 20 μM ATP alone (Figure 4). This indicates that the reaction is dominated by primed RNA synthesis under these conditions. Although the reaction could be made still faster at higher A2 primer concentrations, the selected concentrations balance the ease of measurement with cost.

### 3.6 Inhibition of DENV2 and ZIKV NS5 RdRp activity by Gal-TP

The standardised assay conditions for testing inhibition of RdRp activity by Gal-TP are summarized in Table 2 (see also Section 2.7). In all other experiments enzymatic reactions were initiated by addition of ATP, but for the inhibition assays the reaction was initiated by addition of the NS5 protein. This avoids extended incubation of NS5 proteins with Gal-TP, which could conceivably result in the formation of ppp(Gal)p(Gal) dinucleotides and the initiation of RNA synthesis. DENV2 NS5 catalyses the formation of dinucleotide pppApG primers when incubated with ATP and GTP in the presence of Mn^2+^, even in the absence of template (Selisko et al., 2012).

**Table 2:**
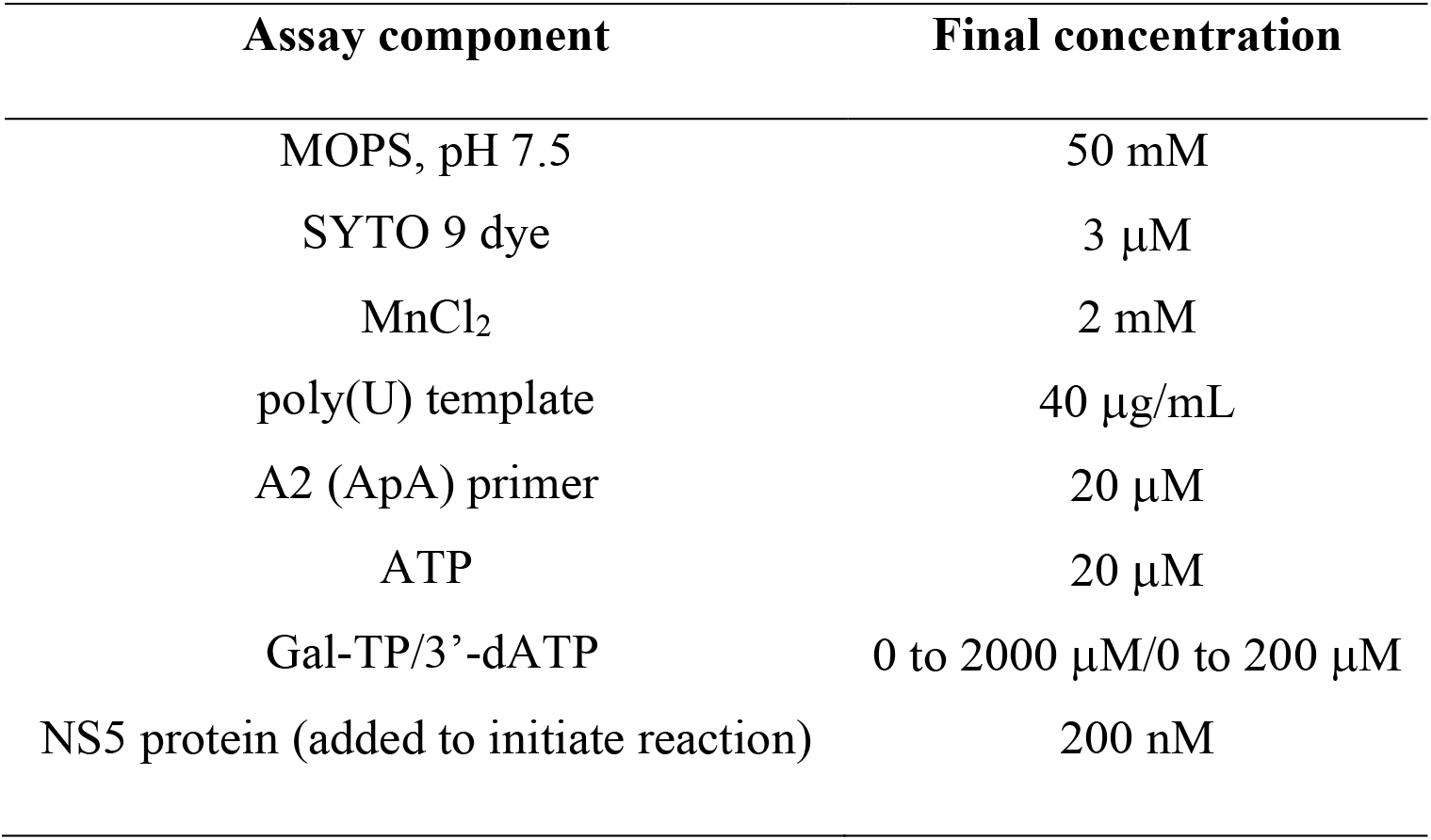
Assay components for NS5 RdRp inhibition assays.

Three independent dose-response experiments were carried out for DENV2 and ZIKV NS5 ([Gal-TP] = 0 – 2 mM) and the data fitted to a four-parameter logistic model (Figure 5). Gal-TP was equipotent against the RdRp activity of DENV2 and ZIKV NS5, with mean IC_50_ values of 42 ± 12 μM and 47 ± 5 μM, respectively (Table 3). The potency of Gal-TP inhibition is of the same order as the *K*_M_^app^(ATP) for DENV2 NS5 under similar assay conditions (Table 1). Hence Gal-TP and ATP may bind the NS5 RdRP with similar affinity, reflecting their highly analogous structures (Figure 1).

**Figure 5:**
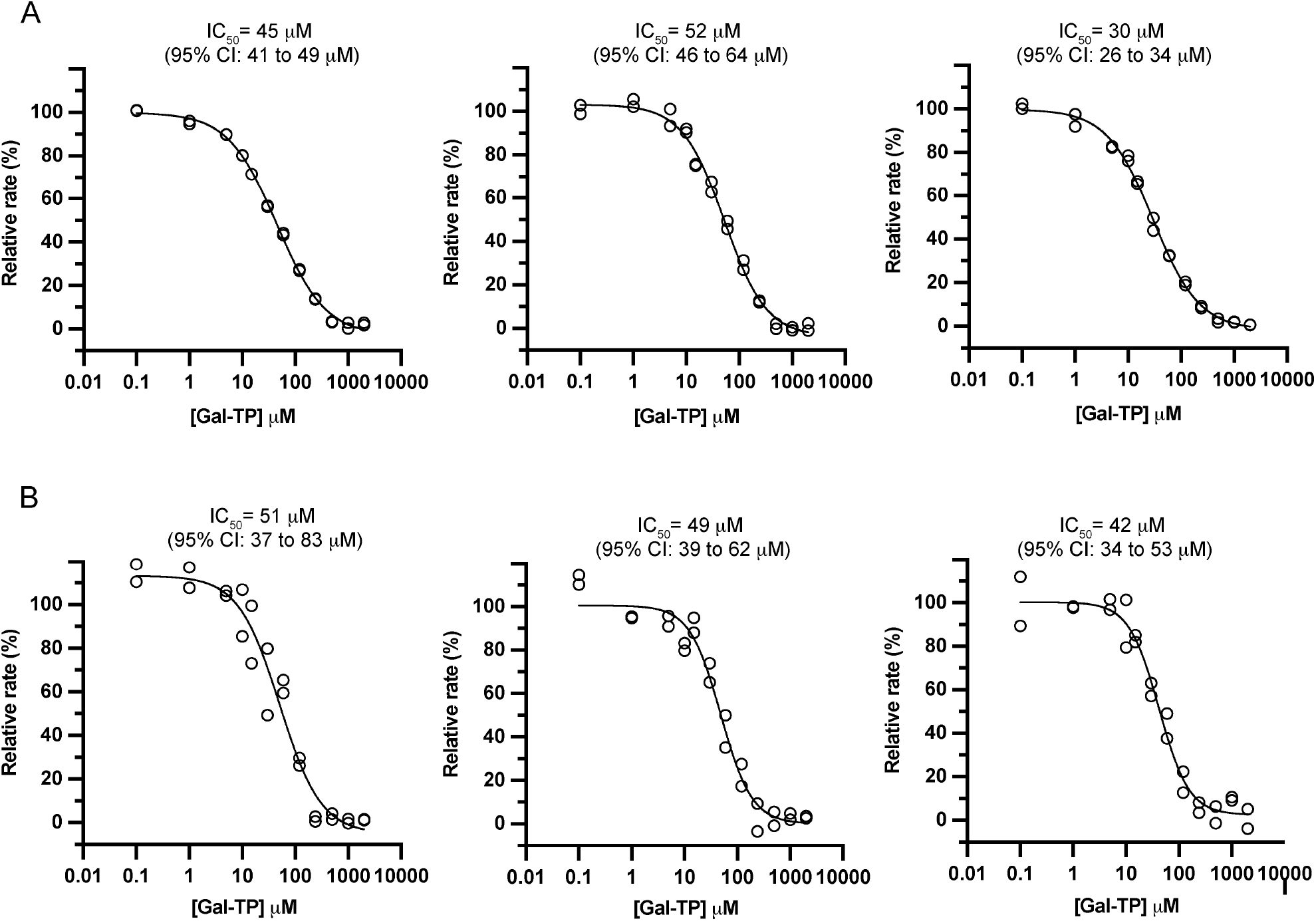
Inhibition of DENV2 (Panel A) and ZIKV (Panel B) NS5 RdRp activity by Gal-TP in A2 primed reactions. Dose-response curves for Gal-TP against NS5 RdRp activity were determined in three independent experiments as shown. Inhibition assay components are given in Table 2. Rates are given as a percentage relative to the uninhibited rate in the absence of Gal-TP. Data (hollow circles) were fit to a four-parameter logistic model (solid lines) to determine IC_50_ values.

**Table 3:** Inhibition constants for Gal-TP and 3’-dATP against DENV2 and ZIKV NS5 RdRp activity.

| Nucleotide analogue | NS5 | $IC_{50}^*$<br>($\mu M$, mean ± std dev.) | Slope parameter*<br>($b$, mean ± std dev.) |
| --- | --- | --- | --- |
| Gal-TP | DENV2 | $42 \pm 12$ | $-1.0 \pm 0.1$ |
| Gal-TP | ZIKV | $47 \pm 5$ | $-1.4 \pm 0.2$ |
| 3'-dATP | DENV2 | $0.48 \pm 0.05$ | $-1.2 \pm 0.1$ |
| 3'-dATP | ZIKV | $0.06 \pm 0.02$ | $-0.9 \pm 0.1$ |

\* Mean and standard deviation from three independent experiments. The four parameter logistic model used for data fitting is detailed in section 2.7.

To validate the inhibition assay, the potency of the obligate chain terminator 3’-deoxyadenosine triphosphate (3’-dATP or cordycepin triphosphate) was also determined against DENV2 and ZIKV NS5 (Figure 6). Relative to Gal-TP, 3’-dATP was two-orders of magnitude more potent against DENV2 NS5 and almost three-order of magnitudes more potent against ZIKV NS5, with IC50 values of 0.48 ± 0.05 μM and 0.06 ± 0.02 μM, respectively (Table 3). Although direct comparisons are difficult due to dependence of IC_50_ on assay conditions, these results are generally consistent with the low to sub-micromolar IC_50_ values reported previously for 3’-dATP against the processive RdRp activity of DENV2 and ZIKV NS5 in various discontinuous assays (Niyomrattanakit et al., 2011; Latour et al., 2010; Lin et al., 2019). The much higher potency of an ATP analog lacking a 3’-hydroxyl, compared with Gal-TP and other ATP analogs, could indicate that there is a binding penalty associated with this substituent.

**Figure 6:**
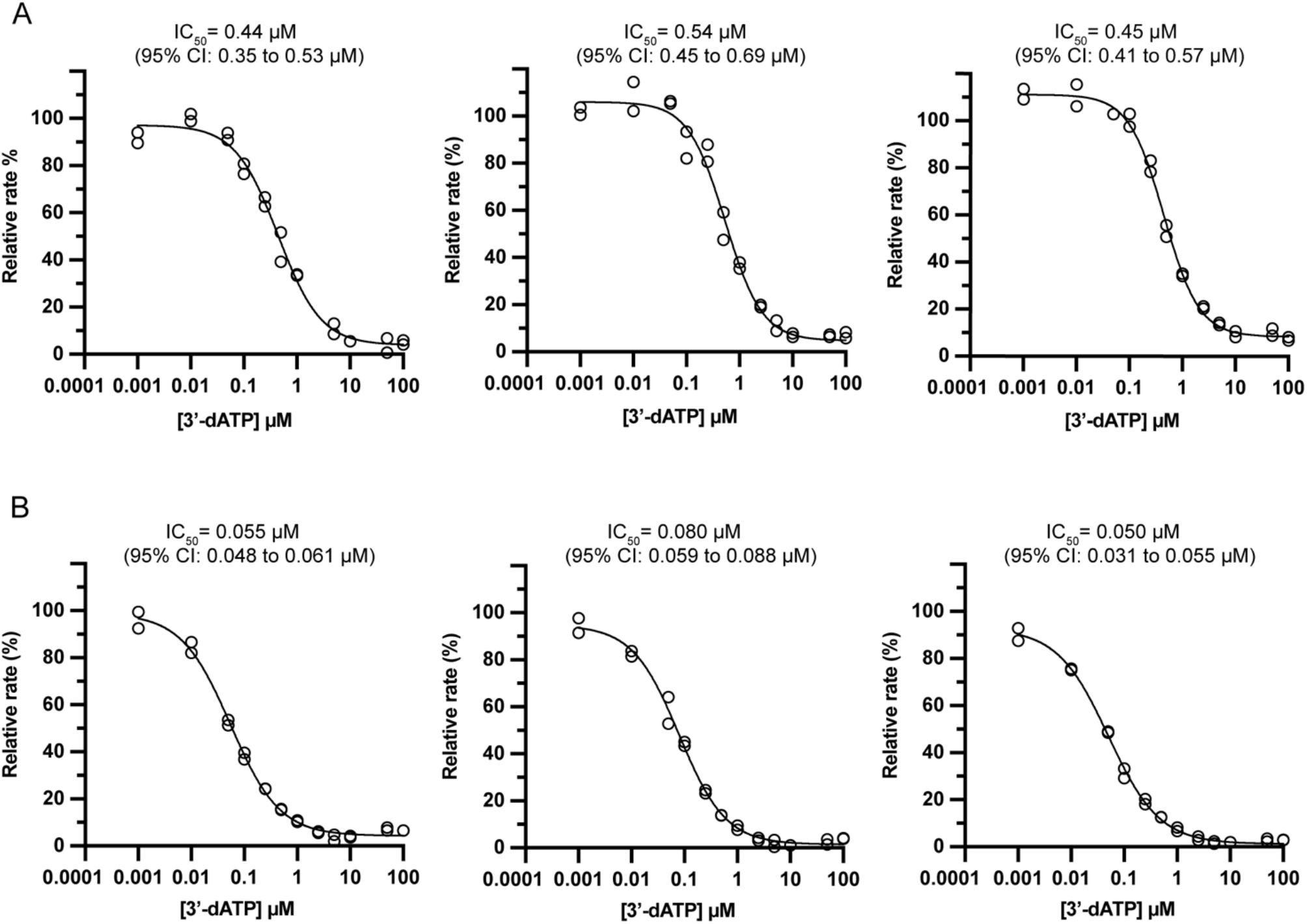
Inhibition of DENV2 (Panel A) and ZIKV (Panel B) NS5 RdRp activity by 3’-dATP in A2 primed reactions. Dose-response curves for 3’-dATP against NS5 RdRp activity were determined in three independent experiments as shown. Inhibition assay components are given in Table 2. Rates are given as a percentage relative to the uninhibited rate in the absence of 3’-dATP. Data (hollow circles) were fit to a four-parameter logistic model (solid lines) to determine IC_50_ values.

The modest *in vitro* potency of Gal-TP against DENV2 and ZIKV NS5 RdRp activity determined in this study, is consistent with previous mammalian cell-based antiviral assays. A 50% effective concentration (EC_50_) of 32.8 μM, was reported for Galidesivir against DENV2 replication in a monkey kidney-derived cell line (Warren et al., 2014). Similarly, EC_50_ values ranging from 3.8 to

11.7 μM were reported for Galidesivir against three different ZIKV strains in various cell lines (Julander et al., 2017). Reported EC_50_ values for Galidesivir in cell-based assays for other flaviviruses vary from 1.5 to 44 μM (Julander et al., 2014; Warren et al., 2014; Eyer et al., 2017; Julander et al., 2017; Julander et al., 2021). Despite this apparently modest potency *in vitro*, Galidesivir abrogates flavivirus infection in animal models, including against Zika virus in Rhesus macaques (Julander et al., 2014; Julander et al., 2017; Lim et al., 2020).

### 3.7 Comparison of Gal-TP with other ATP analog inhibitors of flavivirus NS5 RdRp activity

In the literature, reported IC_50_ and *K*_i_ values for various ATP analogs against flavivirus NS5 RdRp activity *in vitro* are typically in the low-to sub-micromolar range (Eyer et al., 2018). For example, an ATP analog with a 2’-C-ethylnyl substituent on the ribose (7-deaza-2’-C-ethynyladenosine, NITD008) was estimated to have a *K*_i_^app^ of 0.74 ± 0.072 μM against DENV2 NS5 in a radiometric assay and using a template based on the native 3’-UTR sequence (Latour et al., 2010). Analysis of RNA products by gel electrophoresis suggested that incorporation of this nucleoside analog leads to immediate chain termination.

Two ATP analogs carrying 2’-C-methyl substituents on the ribose (2’-C-methyladenosine and 7-deaza-2’-C-methyladenosine) had IC_50_ values of 5.6 ± 0.07 μM and 7.9 ± 1.5 μM, respectively, in a radiometric assay with a poly(U) template (Hercík et al., 2017). NMR studies with poliovirus RdRp suggest that once incorporated the 2-C’-methyl substituent prevents incorporation of the following nucleotide by blocking closure of the RdRp active site (Boehr et al., 2019).

Remdesivir, is a 1’-C-cyano substituted C-nucleoside analog of adenosine that is effective against various flaviviruses in cell-based assays, with EC_50_ values in the range 10 – 50 μM, similar to Galidesivir. Studies with coronavirus polymerases using Remdesivir triphosphate show that three nucleotides are added following incorporation of this nucleoside analog, the polymerase then stalls due to a steric clash between the 1’-C-cyano substituent and a residue sidechain (Kokic et al., 2021; Wang et al., 2020; Gordon et al., 2020b; Gordon et al., 2020a). However, this stalling is overcome in the presence of high concentrations of NTPs (Kokic et al., 2021). This result may explain a report that flavivirus NS5s produced full length RNA products despite incorporation of Remdesivir (Konkolova et al., 2020).

The mechanism of action of Galidesivir has not been fully elucidated. For HCV NS5B RdRp it was shown that following incorporation of Galidesivir up to two further nucleotides are added before apparent termination of RNA synthesis (Warren et al., 2014). However, the generality of this result has not been established. Unlike the adenosine analogs described above, Galidesivir does not carry additional substituents (Figure 1), and hence must function through a different mechanism. The production of Gal-TP as described in this study, will facilitate further work to probe the mechanism of action of Galidesivir.

## 4. Conclusion

Given the impacts of flaviviruses on human health, and the likelihood of increased viral migration in a warming and interconnected world, there is a need to develop effective antiviral compounds. Nucleoside analogs such as Galidesivir are a promising avenue to pursue in antiviral development. In this study we synthesised Gal-TP and tested its inhibition of the RdRp activity of DENV2 and ZIKV NS5 using a dinucleotide-primed, continuous, fluorescence assay. As far as we are aware this is the first direct quantitative testing of the effects of Gal-TP on virally-directed RNA synthesis.

Gal-TP had modest and equivalent micromolar potency against DENV2 and ZIKV NS5 RdRp activity. In contrast, the obligate chain terminator 3’-dATP had sub-micromolar potency against both NS5 proteins in the inhibition assay, consistent with prior reports. Establishment of a straightforward continuous assay for processive viral RdRp activity, will allow the rapid testing of many other classes of nucleoside analogs for inhibitory effects.

## Acknowledgements

This work was supported by grants from the New Zealand Ministry of Business Innovation and Employment (UOOX1904), and the Maurice Wilkins Centre for Molecular Biodiscovery. EMMB was funded by a Faculty Research Development Fund Grant from the University of Auckland. We thank David Goldstone (School of Biological Sciences, University of Auckland) for assistance with SEC-MALLS.

## Abbreviations

DENV: Dengue virus
EDTA: Ethylenediaminetetraacetic acid
Gal-TP: Galidesivir triphosphate
HEWL: Hen egg-white lysozyme
IMAC: Immobilized metal affinity chromatography
JEV: Japanese encephalitis virus
MALLS: Multi-angle laser light scattering
MOPS: 3-(*N*-morpholino)propanesulfonic acid RdRp
RNA: dependent RNA polymerase SEC Size exclusion chromatography
TBEV: Tick-borne encephalitis virus
TCEP: Tris(2-carboxyethyl)phosphine
TEV: Tobacco etch virus
WNV: West Nile virus
YFV: Yellow fever virus
ZKV: Zika virus

