## Supplementary Information for "Inhibition of the DENV2 and ZIKV RNA polymerases by Galidesivir triphosphate measured using a continuous fluorescence assay"

#### Section A: Supplementary Figures

Supplementary Figures 1 to 5.

#### Section B: Chemical synthesis of Galidesivir triphosphate

Synthesis scheme for Galidesivir triphosphate, experimental details and compound characterization data.

#### Section A: Supplementary Figures

```
                                                                                                 DENV2 NS5
    NcoI                     His6-tag              SpeI         spacer          TEV cleavage site     1   2   3        898 899 900        XhoI
     Met Gly Ser Ser His His His His His His Thr Ser Gly Ser Gly Ser Glu Asn Leu Tyr Phe Gln Gly Thr Gly  …  Val Leu Trp ---
ACC ATG GGC AGC AGC CAT CAT CAT CAT CAT CAC ACT AGT GGG AGT GGC AGC GAA AAC CTG TAT TTT CAG GGA ACT GGC  …  GTT CTG TGG TGA CTC GAG

                                                                                                  ZIKV NS5
    NcoI                     His6-tag              SpeI         spacer          TEV cleavage site     1   2   3        901 902 903        XhoI
     Met Gly Ser Ser His His His His His His Thr Ser Gly Ser Gly Ser Glu Asn Leu Tyr Phe Gln Gly Gly Gly  …  Gly Val Leu ---
ACC ATG GGC AGC AGC CAT CAT CAT CAT CAT CAC ACT AGT GGG AGT GGC AGC GAA AAC CTG TAT TTT CAG GGA GGT GGG  …  GGA GTG TTG TGA CTC GAG
```

Supplementary Figure S1. Schematic diagram showing the detail of NS5 gene insertion into the multiple cloning site of vector pET15b(+) (Novagen)

```
DENV2_NS5           GTGNIGETLGEKWKSRLNALGKSEFQIYKKSGIQEVDRTLAKEGIKRGE-TDHHAVSRGS  59
ZKV_NS5             -GGGTGETLGEKWKARLNQMSALEFYSYKKSGITEVCREEARRALKDGVATGGHAVSRGS  59
                      *. *********:*** :.  **  ****** ** *  *:..:* *  *. *******

DENV2_NS5           AKLRWFVERNMVTPEGKVVDLGCGRGGWSYYCGGLKNVREVKGLTKGGPGHEEPIPMSTY  119
ZKV_NS5             AKIRWLEERGYLQPYGKVVDLGCGRGGWSYYAATIRKVQEVRGYTKGGPGHEEPMLVQSY  119
                    **:**: **. : * ****************.. :::*:**:* **********: :.:*

DENV2_NS5           GWNLVRLQSGVDVFFIPPEKCDTLLCDIGESSPNPTVEAGRTLRVLNLVENWLNNN-TQF  178
ZKV_NS5             GWNIVRLKSGVDVFHMAAEPCDTLLCDIGESSSSPEVEETRTLRVLSMVGDWLEKRPGAF  179
                    ***:***:******.:  * ************ .* **  ******.:* :**::.   *

DENV2_NS5           CIKVLNPYMPSVIEKMEALQRKYGGALVRNPLSRNSTHEMYWVSNASGNIVSSVNMISRM  238
ZKV_NS5             CIKVLCPYTSTMMETMERLQRRHGGGLVRVPLCRNSTHEMYWVSGAKSNIIKSVSTTSQL  239
                    ***** **  :::*.** ***::**.*** **.***********.*..**:.**.  *::

DENV2_NS5           LINRFTMRYKKATYEPDVDLGSGTRNIGIESEIPNLDIIGKRIEKIKQEHETSWHYDQDH  298
ZKV_NS5             LLGRMDGPRRPVKYEEDVNLGSGTRAVASCAEAPNMKIIGRRIERIRNEHAETWFLDENH  299
                    *:.*:    : ..** **:****** :.  :* **:.***:***:*::**  :*. *::*

DENV2_NS5           PYKTWAYHGSYETKQTGSASSMVNGVVRLLTKPWDVVPMVTQMAMTDTTPFGQQRVFKEK  358
ZKV_NS5             PYRTWAYHGSYEAPTQGSASSLVNGVVRLLSKPWDVVTGVTGIAMTDTTPYGQQRVFKEK  359
                    **:*********:   *****:********:******  ** :*******:*********

DENV2_NS5           VDTRTQEPKEGTKKLMKITAEWLWKELGKKKTPRMCTREEFTRKVRSNAALGAIFTDENK  418
ZKV_NS5             VDTRVPDPQEGTRQVMNIVSSWLWKELGKRKRPRVCTKEEFINKVRSNAALGAIFEEEKE  419
                    ****. :*:***:::*:*.:.********:* **:**:*** .************ :*::

DENV2_NS5           WKSAREAVEDSRFWELVDKERNLHLEGKCETCVYNMMGKREKKLGEFGKAKGSRAIWYMW  478
ZKV_NS5             WKTAVEAVNDPRFWALVDREREHHLRGECHSCVYNMMGKREKKQGEFGKAKGSRAIWYMW  479
                    **:* ***:* *** ***:**: **.*:*.:************ ****************

DENV2_NS5           LGARFLEFEALGFLNEDHWFSRENSLSGVEGEGLHKLGYILRDVSKKEGGAMYADDTAGW  538
ZKV_NS5             LGARFLEFEALGFLNEDHWMGRENSGGGVEGLGLQRLGYILEEMNRAPGGKMYADDTAGW  539
                    *******************:.**** .**** **::*****.:::.:  ** *********

DENV2_NS5           DTRITLEDLKNEEMVTNHMEGEHKKLAEAIFKLTYQNKVVRVQRPTPRG-TVMDIISRRD  597
ZKV_NS5             DTRISKFDLENEALITNQMEEGHRTLALAVIKYTYQNKVVKVLRPAEGGKTVMDIISRQD  599
                    ****:  **:** ::**:**  *:.** *::* *******:* **:  * ********:*

DENV2_NS5           QRGSGQVGTYGLNTFTNMEAQLIRQMEGEGVFKSIQHLTITEEIAVQNWLARVGRERLSR  657
ZKV_NS5             QRGSGQVVTYALNTFTNLVVQLIRNMEAEEVLEMQDLWLLRKPEKVTRWLQSNGWDRLKR  659
                    ******* **.******: .****:**.* *::  :   : :   * .**   * :**.*

DENV2_NS5           MAISGDDCVVKPLDDRFASALTALNDMGKIRKDIQQWEPSRGWNDWTQVPFCSHHFHELI  717
ZKV_NS5             MAVSGDDCVVKPIDDRFAHALRFLNDMGKVRKDTQEWKPSTGWSNWEEVPFCSHHFNKLY  719
                    **:*********:***** **  ******:*** *:*:** **.:* :********::*

DENV2_NS5           MKDGRVLVVPCRNQDELIGRARISQGAGWSLRETACLGKSYAQMWSLMYFHRRDLRLAAN  777
ZKV_NS5             LKDGRSIVVPCRHQDELIGRARVSPGAGWSIRETACLAKSYAQMWQLLYFHRRDLRLMAN  779
                    :**** :*****:*********:* *****:******.*******.*:********* **

DENV2_NS5           AICSAVPSHWVPTSRTTWSIHAKHEWMTTEDMLTVWNRVWIQENPWMEDKTPVESWEEIP  837
ZKV_NS5             AICSAVPVDWVPTGRTTWSIHGKGEWMTTEDMLMVWNRVWIEENDHMEDKTPVTKWTDIP  839
                    ******* .****.*******.* ********* *******:**  ******* .* :**

DENV2_NS5           YLGKREDQWCGSLIGLTSRATWAKNIQAAINQVRSLIGNE-EYTDYMPSMKRFRREE-EE  895
ZKV_NS5             YLGKREDLWCGSLIGHRPRTTWAENIKDTVNMVRRIIGDEEKYMDYLSTQVRYLGEEGST  899
                    ******* *******   *:***:**: ::* ** :**:* :* **: :  *:  ** .

DENV2_NS5           AGVLW      900
ZKV_NS5             PGVL-      903
                     ***
```

Supplementary Figure S2: Sequence alignment for full length NS5s from DENV2 and ZIKV. Sequences were aligned in Clustal Omega (McWilliam et al., 2013).

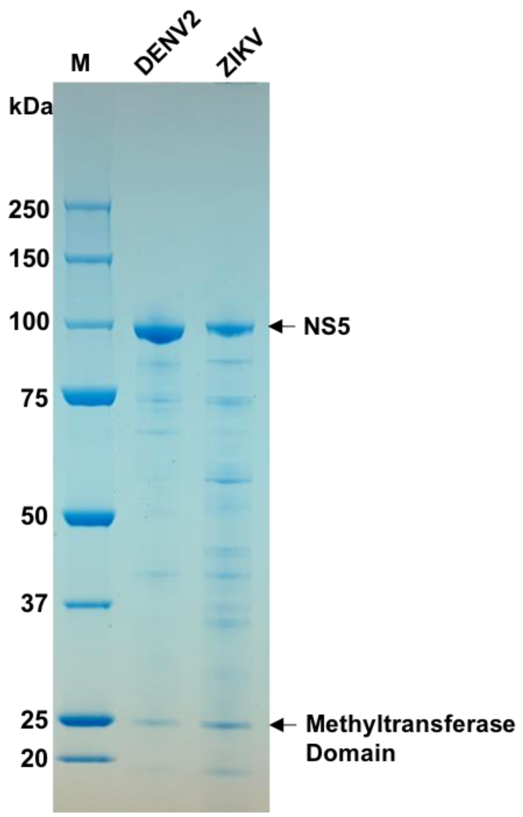

Supplementary Figure S3. Analysis of purified flavivirus NS5 proteins by SDS-PAGE using a NuPAGE™ 4 to 12% Bis-Tris gel and 1x NuPAGE™ MOPS SDS running buffer. Proteins were visualized using colloidal Coomassie staining.

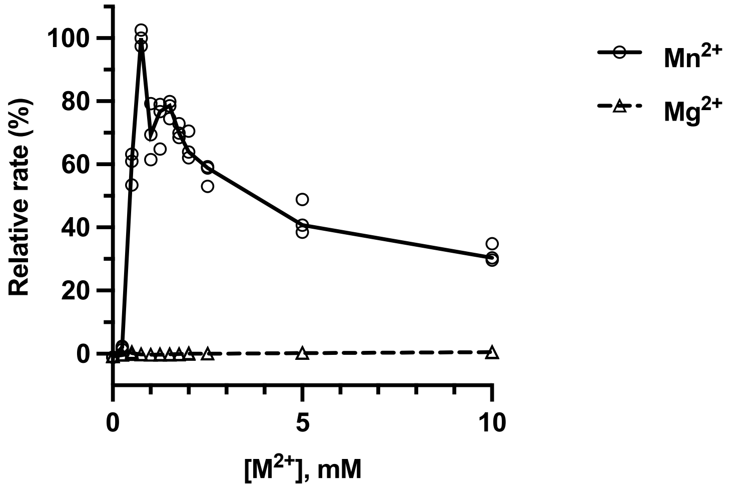

Supplementary Figure S4: Comparison of the RdRp activity of DENV2 NS5 in the presence of $Mn^{2+}$ and $Mg^{2+}$ ions. Reactions contained 200 nM DENV2 NS5, 2.5 μM SYTO 9, 40 μg/ml poly(U), 0 to 10 mM $MnCl_2$ or $MgCl_2$ and were initiated with 0.5 mM ATP. Measured rates in fluorescence units per second are given as a percentage relative to the maximum rate for that experiment. Technical triplicates are shown, connected by a line passing through the median of each set of triplicates.

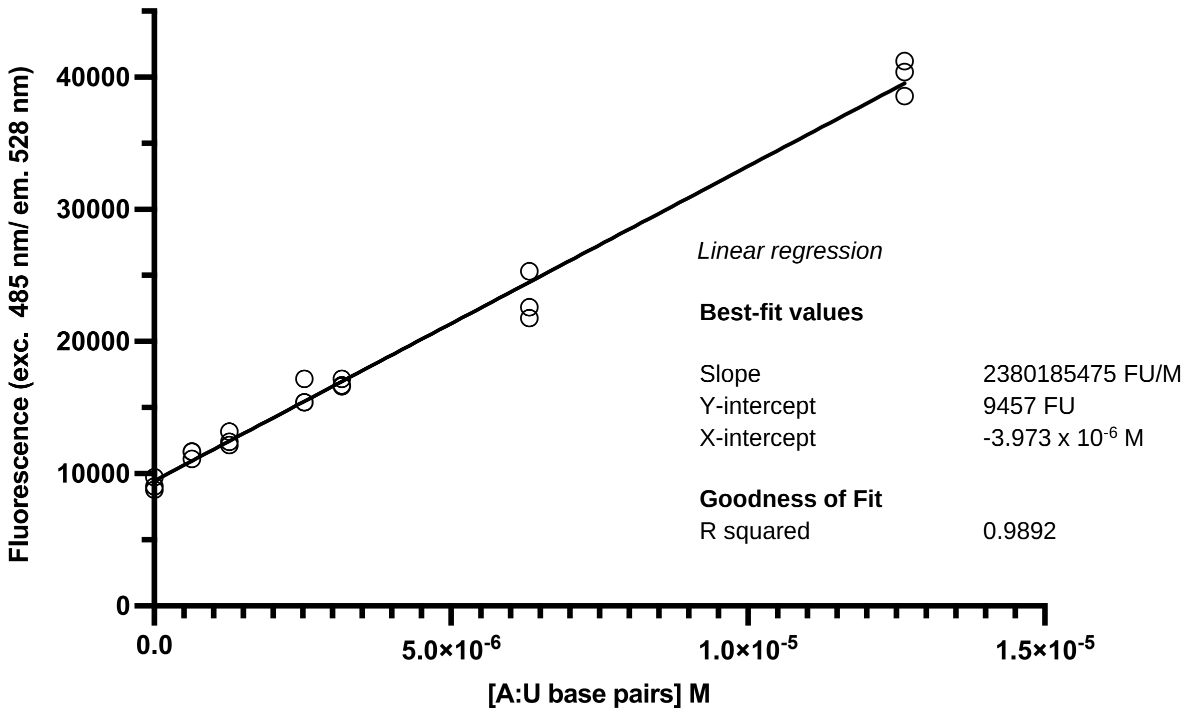

Supplementary Figure S5: Example of a poly(A):poly(U) standard curve generated using the NS5 RdRp assay conditions (200 nM DENV2 NS5, 20 μM A2 (ApA) primer, 40 μg/ml poly(U), 2.5 mM $MnCl_2$, 3 μM SYTO 9). The poly(A):poly(U) concentration was varied from 0 to 8 μg/ml. This corresponds to an effective A:U base pair concentration of 0 to 12.6 x $10^{-6}$ M, given a molar mass of 633.07 g/mol for each A:U base pair.

#### Section B: Chemical Synthesis of Galidesivir Triphosphate

**Synthetic scheme for Galidesivir Triphosphate**

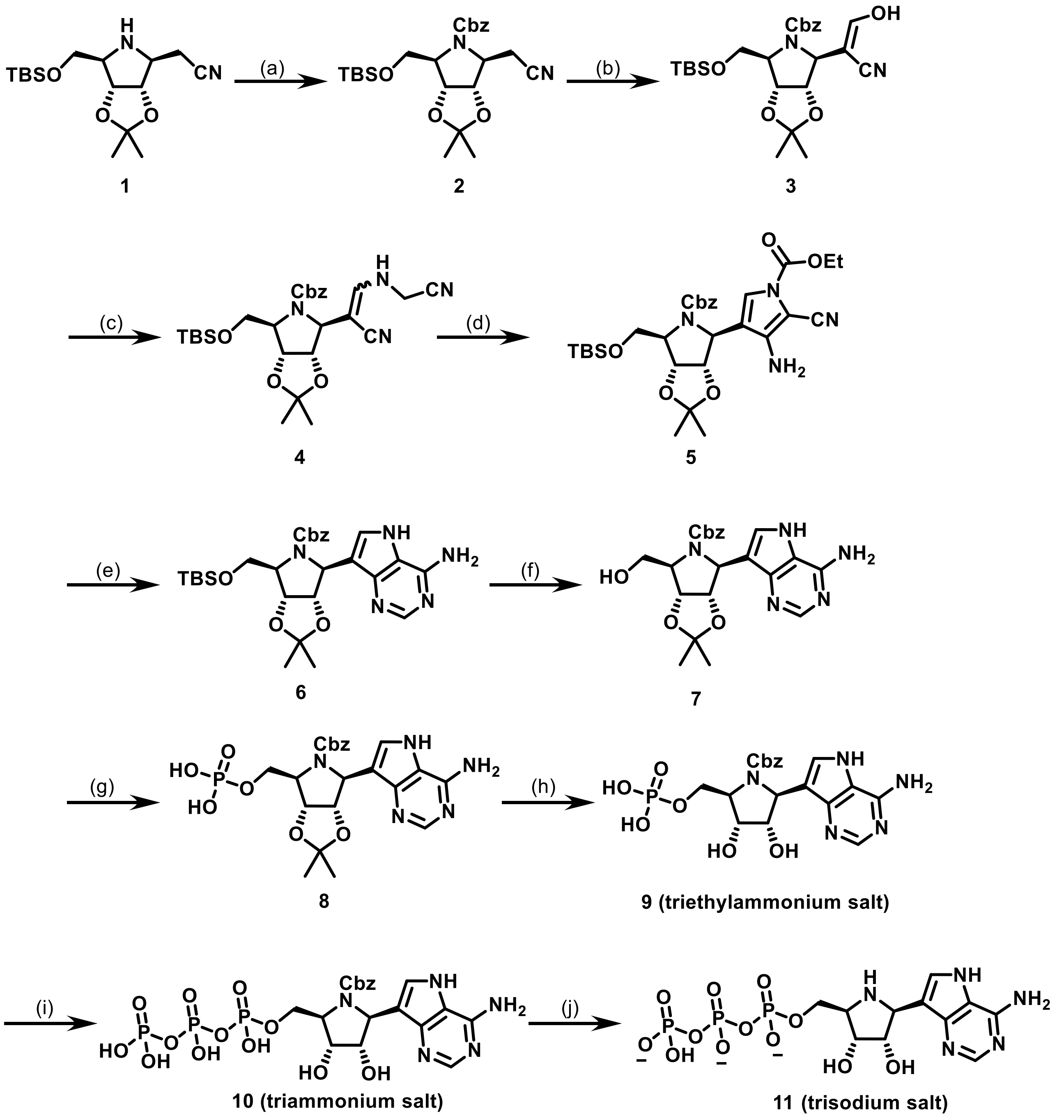

**Scheme (I)**: (a) Cbz-Cl, $K_2CO_3$, PhMe, $H_2O$, 1 h, rt, 96% yield; (b) (i) *t*-BuOCH($NMe_2$)$_2$, DMF, 70 °C, 1.5 h; (ii) THF, AcOH, $H_2O$ (1:1:1 v/v/v), 2 h; 71% yield over 2 steps; (c) aminoacetonitrile bisulfate, NaOAc, MeOH, reflux, 3 h; (d) $ClCO_2Et$, DBU, DCM, reflux, 2 h; 67% yield over 2 steps; (e) (i) $K_2CO_3$, EtOH, rt, 1 h; (ii) formamidine acetate, EtOH, reflux, 20 h; 86% yield over 2 steps; (f) TBAF, THF, rt, 20 h, 91% yield; (g) $POCl_3$, (MeO)$_3$PO, 0 °C for 1 h, rt for 2 h, then TEAB buffer (2 M aq.), 36% yield; (h) (i) TFA:DCM (1:1 v/v), rt, 20 h, 67% yield; (i) (i) CDI, DMF, rt, 3 h; (ii) 5% $Et_3N$ in MeOH:$H_2O$ (1:1 v/v), rt, 4 h; (iii) 2(*n*-$Bu_3N$)×$H_4P_2O_7$, *n*-$Bu_3N$, DMF, rt, 20 h, then TEAB buffer (2 M aq.); (iv) ion-pair chromatography, then ion exchange with DOWEX® 50W X8 $NH_4^+$ form; 56% yield; (j) (i) $H_2$, Pd/C, water, rt, 1 h; (ii) ion-pair chromatography, then ion exchange with DOWEX® 50W X8 $Na^+$ form; 66% yield.

**Experimental Section**

Bis(tributylammonium) pyrophosphate[1] and compound **1**[2] were prepared according to literature procedures. Reactions requiring anhydrous conditions were carried out in flame-dried glassware under a positive pressure of argon in anhydrous solvents, using standard Schlenk techniques. Reaction temperatures above room temperature (22–23 ˚C) were carried out in heating mantles with an internal temperature probe. Reaction progress was monitored by thin layer chromatography (TLC) on Merck aluminum-backed silica gel coated TLC plates (60 Å, F254 indicator). TLC plates were visualized by exposure to ultraviolet light (254 nm), and/or ceric ammonium molybdate stain (Hanessian's Stain). Flash column chromatography was performed with a Büchi Pure C-815 Flash automated flash chromatography system using prepacked FlashPure cartridges containing either silica gel (50 μm irregular) or C18 silica gel (50 μm spherical), and ACS grade solvents. NMR spectra were recorded using a Bruker 500 MHz spectrometer and analyzed using MestReNova software. Data are represented as follows: chemical shift (δ) in parts per million (ppm), multiplicity (s = singlet, d = doublet, dd (doublet of doublets), t = triplet, q = quartet, m = multiplet), coupling constants (*J*) in Hertz (Hz), and integration. High resolution electrospray ionization (ESI) mass spectrometric analysis and liquid chromatography–mass spectrometric analysis (LC-MS) were performed on Waters Q-TOF Premier™ Tandem Mass spectrometer fitted with a Waters 2795 HPLC and analyzed using MassLynx software. LCMS was performed using a ACQUITY UPLC™ BEH C18 column (1.7 µm, 100 × 2.1 mm, 130 Å) and method that used: mobile phase (A: 10 mM ammonium formate in water; B: methanol), flow rate (0.2 mL/min), temperature (30 °C), and detection method (diode array).

*N*-Benzyloxycarbonyl-7-*O*-*tert*-butyldimethylsilyl-2,3,6-trideoxy-3,6-imino-4,5-*O*-isopropylidene-D-*allo*-heptononitrile (**2**)

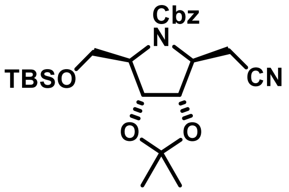

To a mixture of compound **1**[2] (2.7 g, 8.2 mmol, 1.0 equiv.) in toluene (30 mL) and water (30 mL), potassium carbonate (2.3 g, 16 mmol, 2.0 equiv.) was added followed by benzyl chloroformate (1.5 mL, 10 mmol, 1.2 equiv.). This reaction mixture was stirred at room temperature for 1 h. The organic layer was separated, washed with saturated aqueous $NaHCO_3$ solution and brine, dried over $MgSO_4$, filtered, and concentrated *in vacuo*. The crude oil was purified by flash column chromatography (normal phase silica gel, 0-10% *n*-hexane/ethyl acetate) to afford the *title compound* as a colorless oil (3.6 g, 7.8 mmol, 96% yield). $^1$H NMR (500 MHz, $CDCl_3$) δ 7.39 – 7.28 (m, 5H), 5.23 – 5.06 (m, 2H), 4.71 – 4.65 (m, 1H), 4.58 (dd, *J* = 18.5, 5.1 Hz, 1H), 4.28 – 4.09 (m, 2H), 3.92 – 3.64 (m, 2H), 2.99 – 2.67 (m, 2H), 1.46, 1.44, 1.33 (3s, 6H), 0.90, 0.88 (2s, 9H), 0.08, 0.06, 0.03, 0.02 (4s, 6H); $^{13}$C NMR (126 MHz, $CDCl_3$) δ 154.5, 154.2, 136.1, 128.71, 128.68, 128.39, 128.36, 128.1, 128.0, 117.2, 117.1, 112.4, 84.2, 83.1, 82.1, 81.3, 67.62, 67.58, 66.9, 65.9, 63.6, 63.0, 62.5, 61.6, 27.3, 25.3, 26.1, 21.5, 20.6, 18.5, -5.36, -5.43. HRMS (ESI-MS) m/z $[M+H]^+$ calculated for $C_{24}H_{37}N_2O_5Si$ 461.2466, found 461.2470.

*N*-Benzyloxycarbonyl-7-*O*-*tert*-butyldimethylsilyl-2,3,6-trideoxy-3,6-imino-4,5-*O*-isopropylidene-D-*allo*-hept-1-enitol (**3**)

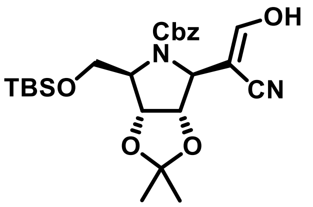

To a solution of Cbz protected compound **2** (3.65 g, 7.93 mmol, 1.00 equiv.) in anhydrous DMF (55 mL), *tert*-butoxy bis(*N*,*N*-dimethylamino)methane (7.3 mL, 32 mmol, 4.0 equiv.) was added. The resulting mixture was stirred at 70 °C for 1.5 h. The reaction mixture was cooled down to room temperature, quenched with brine and extracted with ethyl acetate (3 × 50 mL). The combined organic extracts were washed with brine, dried over $MgSO_4$, filtered, and concentrated *in vacuo*. The crude oil was stirred with a mixture of THF (40 mL), acetic acid (40 mL) and water (40 mL) for 2 h. This mixture was diluted with dichloromethane (100 mL) and water (50 mL) and neutralized with solid $NaHCO_3$. The organic layer was separated, washed with saturated aqueous $NaHCO_3$ solution and brine, dried over $MgSO_4$, filtered, and concentrated *in vacuo*. The crude oil was purified by flash column chromatography (normal phase silica gel, 0-40% *n*-hexane/ethyl acetate) to afford the *title compound* as an orange oil (2.73 g, 5.59 mmol, 70.5% yield). $^1$H NMR (500 MHz, $CDCl_3$) δ 7.41 – 7.29 (m, 5H), 7.14 (s, 1H), 5.20 (s, 2H), 4.95 – 4.90 (m, 1H), 4.86 – 4.80 (m, 2H), 4.16 – 4.07 (m, 1H), 3.65 (dd, *J* = 10.2, 5.5 Hz, 1H), 3.60 – 3.53 (m, 1H), 1.44, 1.33 (2s, 6H), 0.87 (s, 9H), 0.03 (s, 6H); $^{13}$C NMR (126 MHz, $CDCl_3$) δ 162.8, 157.9, 135.3, 128.8, 128.7, 128.1, 118.7, 112.4, 90.3, 82.7, 82.2, 69.0, 66.3, 62.6, 59.7, 27.1, 25.1, 26.0, 18.5, -5.27, -5.31. HRMS (ESI-MS) m/z $[M+H]^+$ calculated for $C_{25}H_{37}N_2O_6Si$ 489.2415, found 489.2416.

(1*S*)-1-(3-Amino-*N*-ethoxycarbonyl-2-cyano-pyrrol-4-yl)-*N*-benzyloxycarbonyl-5-*O*-*tert*-butyldimethylsilyl-1,4-dideoxy-1,4-imino-2,3-*O*-isopropylidene-D-ribitol (**5**)

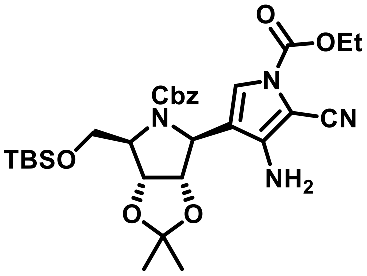

A mixture of compound **3** (2.63 g, 5.39 mmol, 1.00 equiv.), aminoacetonitrile hydrogen sulfate (4.4 g, 28 mmol, 5.2 equiv.) and sodium acetate (5.0 g, 60 mmol, 11 equiv.) in anhydrous methanol (55 mL) was refluxed for 3 h, then concentrated *in vacuo*. The remaining residue was taken up in ethyl acetate (50 mL), washed with brine (3 × 50 mL), dried over $MgSO_4$, filtered, and concentrated *in vacuo* to afford compound **4** as a mixture of isomers (2.7 g, 5.1 mmol, 0.94 equiv.). To a solution of this compound in anhydrous DCM (80 mL), DBU (15 mL, 99 mmol, 20 equiv.) was added followed by ethyl chloroformate (3.5 mL, 36 mmol, 7.1 equiv.) and the mixture heated under reflux for 2 h. The reaction mixture was cooled down to room temperature, diluted with DCM (50 mL), washed with 1 M cold aqueous HCl solution (2 × 75 mL), saturated aqueous $NaHCO_3$ solution

(2 × 75 mL) and brine (1 × 100 mL), dried over $MgSO_4$, filtered, and concentrated *in vacuo*. The crude oil was purified by flash column chromatography (normal phase silica gel, 0-30% *n*-hexane/ethyl acetate) to afford the *title compound* as a white foam (2.03 g, 3.39 mmol, 67% yield). $^1$H NMR (500 MHz, $CDCl_3$) δ 7.44 – 7.13 (m, 5H), 6.99 (s, 1H), 5.20 – 5.02 (m, 2H), 4.87 – 4.61 (m, 3H), 4.46 – 4.36 (m, 2H), 4.32 – 4.14 (m, 1H), 3.88 – 3.46 (m, 2H), 1.50, 1.34 (2s, 6H), 1.40 (t, *J* = 7.1 Hz, 3H), 0.81 (s, 9H), -0.02, -0.04 (2s, 6H); $^{13}$C NMR (126 MHz, $CDCl_3$) δ 155.8, 148.9, 136.2, 128.6, 128.3, 127.9, 122.3, 117.6, 113.5, 112.5, 86.2, 84.1, 82.0, 67.8, 66.2, 64.4, 63.4, 60.7, 27.7, 25.6, 25.9, 18.4, 14.3, -5.4, -5.5. HRMS (ESI-MS) m/z $[M+H]^+$ calculated for $C_{30}H_{43}N_4O_7Si$ 599.2896, found 599.2899.

(1*S*)-1-(9-Deazaadenin-9-yl)-*N*-benzyloxycarbonyl-5-*O*-*tert*-butyldimethylsilyl-1,4-dideoxy-1,4-imino-2,3-*O*-isopropylidene-D-ribitol (**6**)

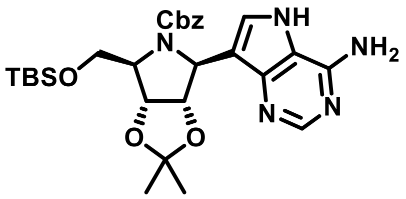

To a solution of compound **5** (1.88 g, 3.14 mmol, 1.00 equiv.) in ethanol (60 mL), $K_2CO_3$ (870 mg, 6.29 mmol, 2.00 equiv.) was added, then stirred at room temperature for 1 h. The reaction mixture was diluted with ethyl acetate (75 mL), washed with water (1 × 75 mL), and brine (1 × 75 mL), dried over $MgSO_4$, filtered, and concentrated *in vacuo* to a white foam. To a solution of this compound (1.68 g, 3.19 mmol) in ethanol (50 mL), formamidine acetate (2.35 g, 22.3 mmol, 7.01 equiv.) was added and the mixture heated under reflux for 20 h. The reaction mixture was concentrated *in vacuo* and purified by flash column chromatography (normal phase silica gel, 0-7% methanol/DCM) to afford the *title compound* as a white foam (1.68 g, 2.74 mmol, 86% yield). $^1$H NMR (500 MHz, $CDCl_3$) δ 8.20 (s, 1H), 7.44 – 7.04 (m, 5H), 6.92 (s, 1H), 6.10 (s, 2H), 5.42 – 5.30 (m, 1H), 5.30 – 4.99 (m, 3H), 4.99 – 4.80 (m, 1H), 4.29 – 4.19 (m, 1H), 3.88 – 3.66 (m, 2H), 1.45, 1.30 (2s, 6H), 0.82 (s, 9H), -0.03 (s, 6H); $^{13}$C NMR (126 MHz, $CDCl_3$) δ 156.1, 150.3, 149.8, 143.8, 136.2, 128.8, 128.4, 127.7, 126.6, 114.8, 114.0, 112.1, 84.3, 82.4, 67.9, 66.6, 62.8, 62.3, 27.4, 25.3, 26.0, 18.4, -5.2. HRMS (ESI-MS) m/z $[M+H]^+$ calculated for $C_{28}H_{40}N_5O_5Si$ 554.2793, found 554.2792.

(1*S*)-1-(9-Deazaadenin-9-yl)-*N*-benzyloxycarbonyl-1,4-dideoxy-1,4-imino-2,3-*O*-isopropylidene-D-ribitol (**7**)

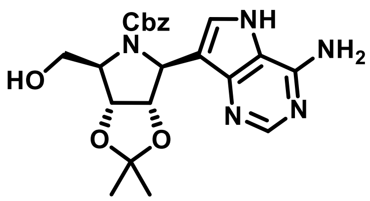

To a solution of compound **6** (700 mg, 1.26 mmol, 1.00 equiv.) in anhydrous THF (20 mL), TBAF (1.0 mol/L in THF) (4.0 mL, 4.0 mmol, 3.2 equiv.) was added, then stirred at room temperature for 4 h. The reaction mixture was concentrated *in vacuo* and purified by flash column chromatography (normal phase silica gel, 0-10% ethanol/DCM) to afford the *title compound* as a white foam (499 mg, 1.14 mmol, 90% yield). $^1$H NMR (500 MHz, $CDCl_3$) δ 9.64 (s, 1H), 8.05 (s, 1H), 7.38 – 7.18 (m, 5H), 7.08 (s, 1H), 5.21 (d, *J* = 13.0 Hz, 1H), 5.08 (d, *J* = 13.0 Hz, 1H), 4.93 (d, *J* = 4.8 Hz, 1H), 4.87 – 4.78 (m, 2H), 4.45 – 4.41 (m, 1H), 4.39 (d, *J* = 11.7 Hz, 1H),

3.79 (dd, $J$ = 12.4, 2.1 Hz, 1H), 1.65, 1.36 (2s, 6H); $^{13}$C NMR (126 MHz, $CDCl_3$) δ 155.8, 150.1, 149.8, 142.5, 136.3, 128.9, 128.4, 127.1, 127.6, 114.9, 114.0, 112.0, 85.5, 82.3, 67.2, 66.4, 65.0, 62.9, 28.3, 25.8. HRMS (ESI-MS) m/z $[M+H]^+$ calculated for $C_{22}H_{26}N_5O_5$ 440.1928, found 440.1931.

Compound **8**

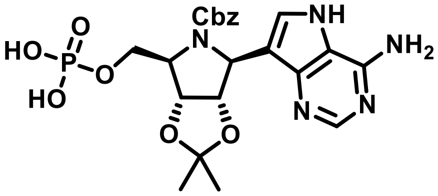

A solution of compound **7** (115 mg, 0.262 mmol, 1.00 equiv.) in trimethyl phosphate (1.3 mL) was stirred at room temperature under argon for 10 min. To this solution cooled down to 0 °C, a solution of phosphoryl chloride (0.15 mL, 1.6 mmol, 6.1 equiv.) in trimethyl phosphate (1.3 mL) was added dropwise, then stirred at 0 °C for 1 h and at 25 °C for 2 h. Consumption of the starting material was confirmed by TLC (normal phase silica gel using a mixture of IPA, $H_2O$ and aqueous $NH_3$ (4:2:4)). The reaction was quenched with 2 M aqueous solution of triethylammonium bicarbonate buffer (pH 7) (4.0 mL, 8.0 mmol, 31 equiv.), concentrated to a white solid, and purified by by flash column chromatography (C18 reversed-phase column, 0-50% water with 0.1% formic acid /methanol) to afford the *title compound* as a white powder (49.0 mg, 0.094 mmol, 36 % yield). $^1$H NMR (500 MHz, MeOD) δ 8.42 – 7.71 (m, 2H), 7.46 – 6.53 (m, 5H), 5.17 – 4.93 (m, 3H), 4.85 – 4.52 (m, 2H), 4.47 – 4.04 (m, 3H), 1.51 (s, 3H), 1.30 (s, 3H); $^{13}$C NMR (126 MHz, MeOD) δ 156.4, 152.6, 145.3, 136.9, 132.1, 129.2, 128.8, 115.0, 113.2, 88.5, 86.7, 83.6, 82.9, 68.5, 66.0, 65.8, 62.0, 61.4, 28.0, 25.8; $^{31}$P NMR (202 MHz, MeOD) δ 0.6. HRMS (ESI-MS) m/z $[M-H]^-$ calculated for $C_{22}H_{25}N_5O_8P$ 518.1446, found 518.1445.

Compound **9**

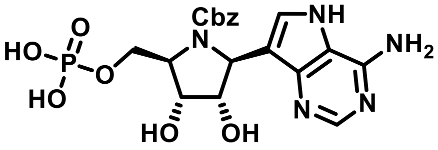

**(triethylammonium salt)**

A solution of compound **8** (36 mg, 69 µmol) was stirred with a mixture of TFA (1 mL) and DCM (1 mL) at room temperature for 20 h. Diol deprotection was monitored by TLC (normal phase silica gel using a mixture of IPA, $H_2O$ and aqueous $NH_3$ (5:2:3)). The reaction mixture was neutralized with triethylamine (0.2 mL, 1 mmol) and concentrated *in vacuo*. The crude product was purified by flash column chromatography (C18 reversed-phase column, 0-50% water/methanol) to afford the *title compound* as a white powder (27 mg, 47 µmol, 67% yield as triethylammonium salt). $^1$H NMR (500 MHz, $D_2O$) δ 8.22 – 7.74 (m, 2H), 7.42 – 6.89 (m, 3H), 6.64 – 6.39 (m, 2H), 4.78 – 4.61 (m, 3H), 4.54 – 4.39 (m, 2H), 4.39 – 4.25 (m, 1H), 4.25 – 4.15 (m, 1H), 4.15 – 4.10 (m, 1H); $^{13}$C NMR (126 MHz, $D_2O$) δ 156.5, 149.4, 143.5, 137.7, 134.6, 130.8, 128.3, 127.8, 113.8, 111.8, 77.7, 72.9, 67.6, 65.9, 63.4, 55.8; $^{31}$P NMR (202 MHz, $D_2O$) δ 2.3. HRMS (ESI-MS) m/z $[M-H]^-$ calculated for $C_{19}H_{21}N_5O_8P$ 478.1133, found 478.1136.

Compound **10**

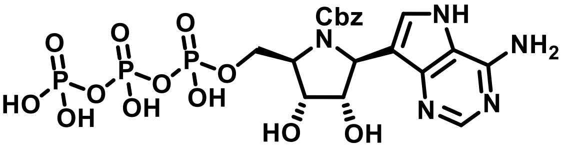

**(triammonium salt)**

1,1′-carbonyldiimidazole (46 mg, 0.28 mmol, 5.9 equiv.) was added to a solution of compound **9** (27 mg, 47 µmol, 1.0 equiv.) in anhydrous DMF (0.3 mL) and stirred under argon at room temperature for 3 h. Consumption of the starting material was monitored by TLC (normal phase silica gel using a mixture of IPA, $H_2O$ and aqueous $NH_3$ (6:1:3)). A 5% triethylamine solution in a mixture of methanol and water (1:1 v/v) (1 mL) was added to the reaction mixture and stirred at room temperature for 4 h. Hydrolysis of cyclic carbonate was monitored by LCMS and upon completion, the reaction mixture was concentrated *in vacuo*. The remaining residue was dissolved in anhydrous DMF (0.3 mL), then stirred with a solution of tributylammonium pyrophosphate (78 mg, 0.14 mmol, 3.1 equiv.) in anhydrous DMF (0.3 mL) at room temperature for 20 h. Conversion of the monophosphate to triphosphate was confirmed by TLC (normal phase silica gel using a mixture of IPA, $H_2O$ and aqueous $NH_3$ (4:2:4)). The reaction mixture was diluted with 2 M aqueous solution of triethylammonium bicarbonate buffer (pH 7) (2.0 mL) and concentrated *in vacuo*. The resulting residue was purified by ion pair chromatography (C18 reversed-phase column, 0-50% mobile phases A/B; mobile phase A composed of deionized water with 10 mM tributylamine and 30 mM acetic acid; mobile phase B composed of methanol with 15 mM tributylamine), then treated with DOWEX® 50W X8 $NH_4^+$ form to afford the *title compound* as a white powder (18 mg, 26 µmol, 56% yield as triammonium salt). $^1H$ NMR (500 MHz, $D_2O$) δ 8.10 – 7.84 (m, 2H), 7.56 – 6.97 (m, 4H), 6.97 – 6.69 (m, 2H), 5.16 (d, *J* = 11.4 Hz, 1H), 4.77 – 4.60 (m, 3H), 4.57 – 4.52 (m, 1H), 4.45 – 4.31 (m, 2H), 4.18 – 4.14 (m, 1H); $^{13}C$ NMR (126 MHz, $D_2O$) δ 156.7, 150.9, 144.0, 134.9, 134.8, 131.4, 131.3, 128.5, 128.2, 127.9, 111.9, 111.8, 77.2, 72.4, 67.9, 65.4, 64.8, 64.7, 55.4. $^{31}P$ NMR (202 MHz, $D_2O$) δ -10.4 (d, *J* = 19.1 Hz), -11.4 (d, *J* = 18.9 Hz), -22.9 (t, *J* = 18.7 Hz). HRMS (ESI-MS) m/z $[M+H]^+$ calculated for $C_{19}H_{25}N_5O_{14}P_3$ 640.0605, found 640.0601.

Compound **11**

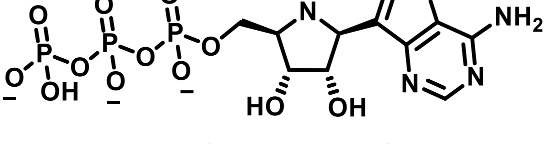

**(trisodium salt)**

A solution of compound **10** (20 mg, 29 µmol) in deionized water (2 mL) was degassed with argon, then stirred with palladium on carbon (10% w/w) (20 mg) under hydrogen atmosphere at room temperature for 1 h. Cbz-deprotection was monitored by TLC (normal phase silica gel using a mixture of IPA, $H_2O$ and aqueous $NH_3$ (5:2:3)). Reaction mixture was filtered through a Celite® pad and lyophilized. Resulting residue was purified by ion pair chromatography (C18 reversed-phase column, 0-30% mobile phases A/B; mobile phase A composed of deionized water with 10 mM tributylamine and 30 mM acetic acid; mobile phase B composed of methanol with 15 mM tributylamine), then treated with DOWEX® 50W X8 $Na^+$ form to afford the *title compound* as a white powder (11.0 mg, 19.3 µmol, 66.5% yield assuming trisodium salt). Full characterization was performed using bis-tributylammonium salt of compound **11**. $^1H$ NMR (500 MHz, $D_2O$) δ 8.31 (s, 1H), 8.05 (s, 1H), 4.99 (d, $J$ = 8.6 Hz, 1H), 4.77 – 4.74 (m, 1H), 4.60 – 4.49 (m, 2H), 4.49 – 4.38 (m, 1H), 4.15 – 4.06 (m, 1H), 3.22 – 3.01 (m, 12H), 1.78 – 1.66 (m, 12H), 1.43 (m, 12H), 0.99 (t, $J$ = 7.4 Hz, 18H); $^{13}C$ NMR (126 MHz, $D_2O$) δ 150.4, 150.3, 146.7, 146.6, 139.2, 139.1, 131.8, 113.0, 112.9, 104.9, 73.4, 70.1, 63.8, 63.7, 63.0, 55.7, 52.7, 47.3, 27.5, 25.2, 19.3, 19.2, 12.8; $^{31}P$ NMR (202 MHz, $D_2O$) δ -8.7 (d, $J$ = 20.2 Hz), -11.6 (d, $J$ = 19.6 Hz), -22.4 (t, $J$ = 17.2 Hz). HRMS (ESI-MS) m/z $[M-H]^-$ calculated for $C_{11}H_{17}N_5O_{12}P_3$ 504.0092, found 504.0093. Purity of the triphosphate was checked by LCMS using diode array as the detection method.

1. Thillier, Y.; Sallamand, C.; Baraguey, C.; Vasseur, J.-J.; Debart, F., Solid-Phase Synthesis of Oligonucleotide 5′-(α-P-Thio)triphosphates and 5′-(α-P-Thio)(β,γ-methylene)triphosphates. *European Journal of Organic Chemistry* **2015,** *2015* (2), 302-308.
2. Evans, G. B.; Furneaux, R. H.; Gainsford, G. J.; Schramm, V. L.; Tyler, P. C., Synthesis of Transition State Analogue Inhibitors for Purine Nucleoside Phosphorylase and N-Riboside Hydrolases. *Tetrahedron* **2000,** *56* (19), 3053-3062.

*N*-Benzyloxycarbonyl-7-*O*-*tert*-butyldimethylsilyl-2,3,6-trideoxy-3,6-imino-4,5-*O*-isopropylidene-D-*allo*-heptononitrile (**2**)

$^1$H NMR (500 MHz, $CDCl_3$)

fe-rsh-av20-046b.10.fid
Owner rsh
Sample RSh-AV20-046b
Platform 3355
Job No fe

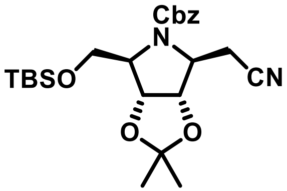

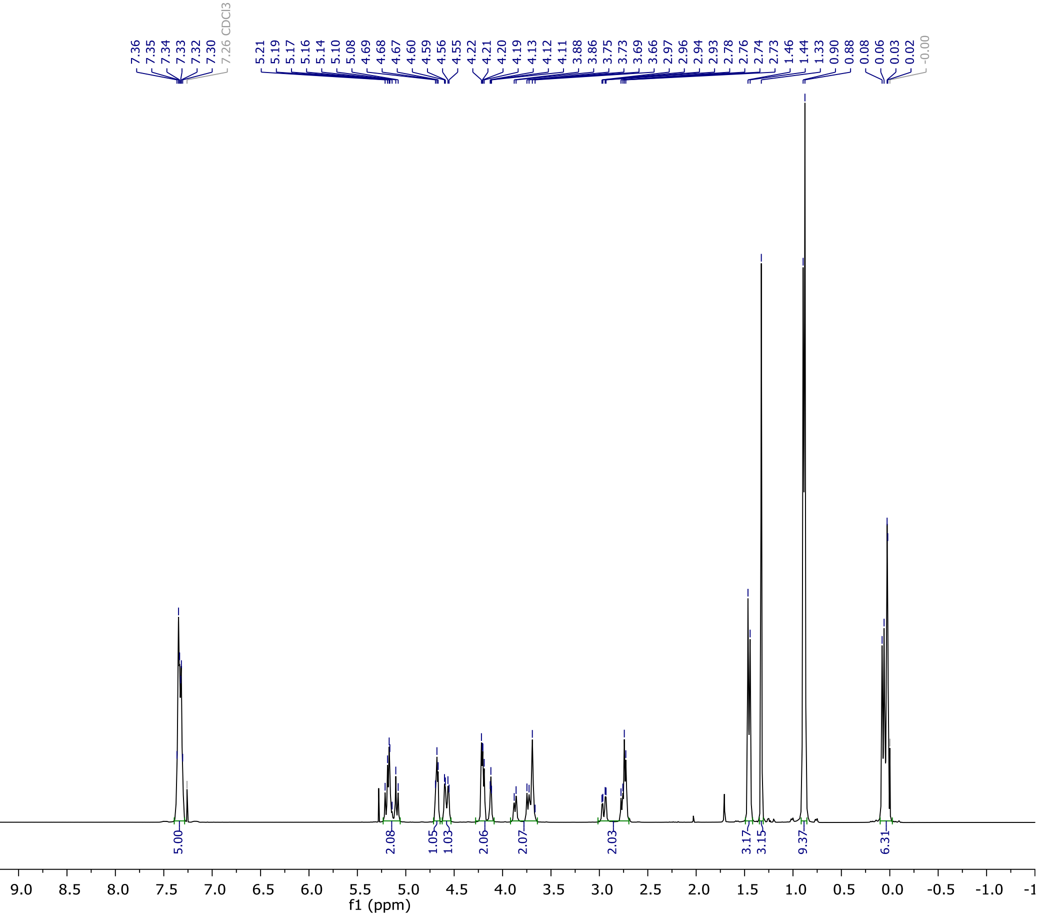

$^{13}C$ NMR (126 MHz, $CDCl_3$)

fe-rsh-av20-046b.11.fid
Owner rsh
Sample RSh-AV20-046b
Platform 3355
Job No fe

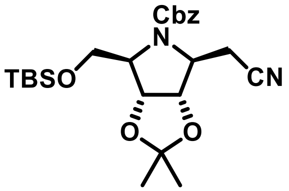

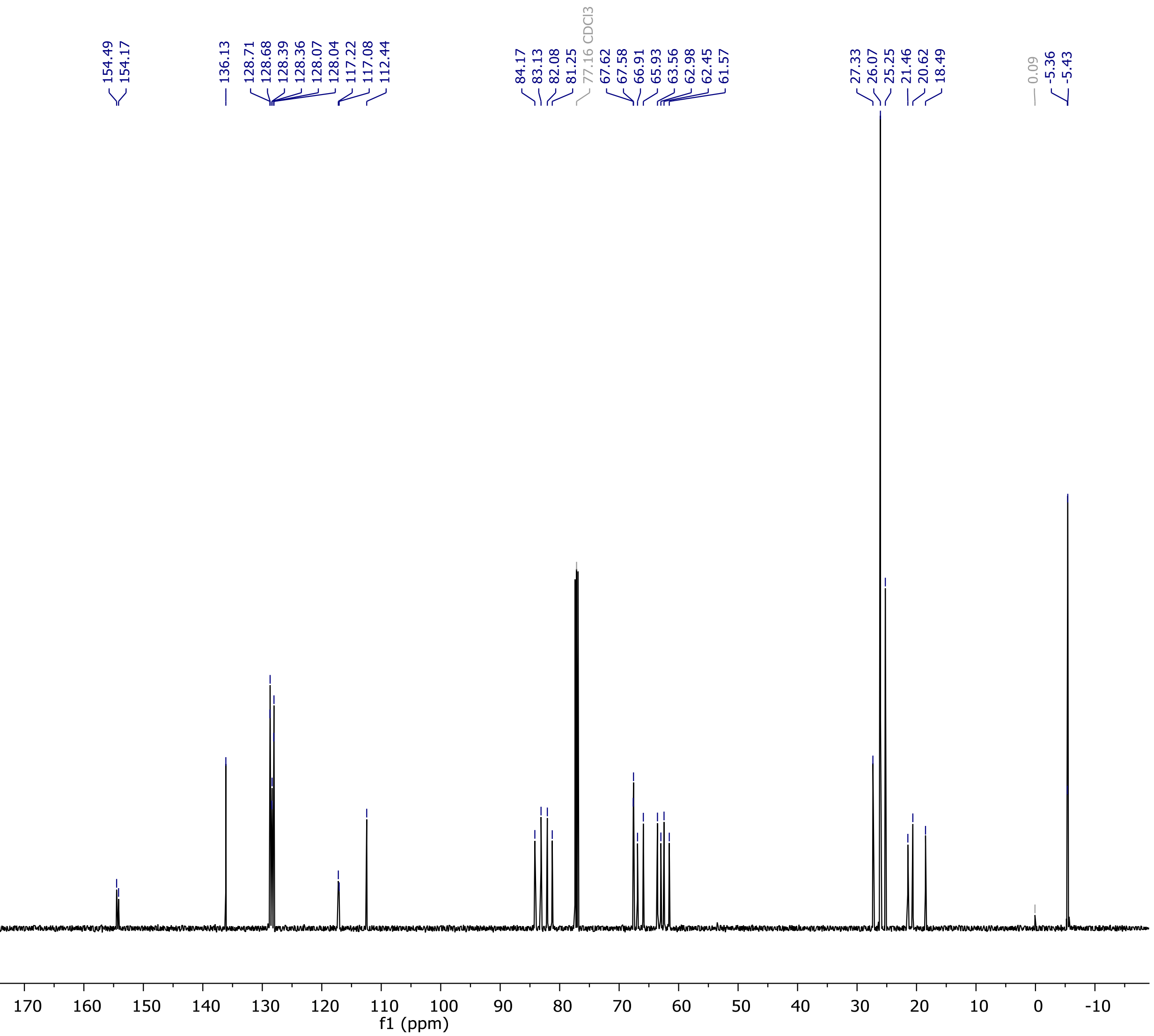

HRMS (ESI-MS)

### Elemental Composition Report

#### Single Mass Analysis

Tolerance = 5.0 mDa / DBE: min = -1.5, max = 150.0
Element prediction: Off
Number of isotope peaks used for i-FIT = 3

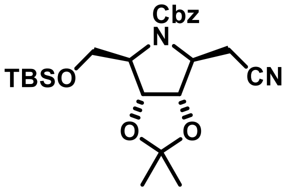

Monoisotopic Mass, Even Electron Ions
165 formula(e) evaluated with 2 results within limits (up to 100 closest results for each mass)
Elements Used:
C: 0-80 H: 0-100 N: 2-2 O: 0-20 Na: 0-1 Si: 1-1

RSh-AV20-046b 17 (0.197) Cm (13:19-25:34)

13-Jul-2020
1: TOF MS ES+
3.33e+007

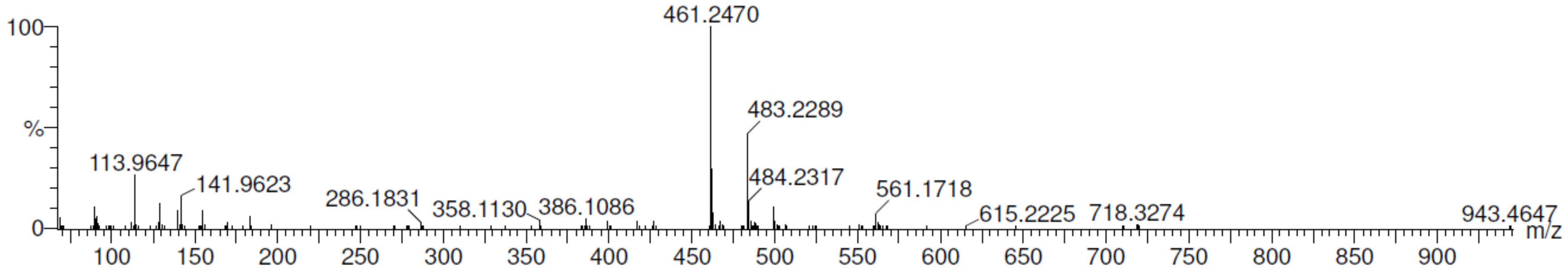

| | | | | | | | | |
|---|---|---|---|---|---|---|---|---|
| Minimum: | | | | -1.5 | | | | |
| Maximum: | | 5.0 | 5.0 | 150.0 | | | | |
| Mass | Calc. Mass | mDa | PPM | DBE | i-FIT | Norm | Conf(%) | Formula |
| 461.2470 | 461.2472 | -0.2 | -0.4 | 8.5 | 87.8 | 0.757 | 46.93 | C24 H37 N2 O5 Si |
| | 461.2448 | 2.2 | 4.8 | 5.5 | 87.7 | 0.634 | 53.07 | C22 H38 N2 O5 Na Si |

*N*-Benzyloxycarbonyl-7-*O*-*tert*-butyldimethylsilyl-2,3,6-trideoxy-3,6-imino-4,5-*O*-isopropylidene-D-*allo*-hept-1-enitol (**3**)

$^1$H NMR (500 MHz, $CDCl_3$)

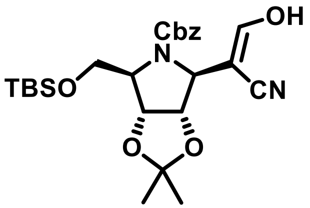

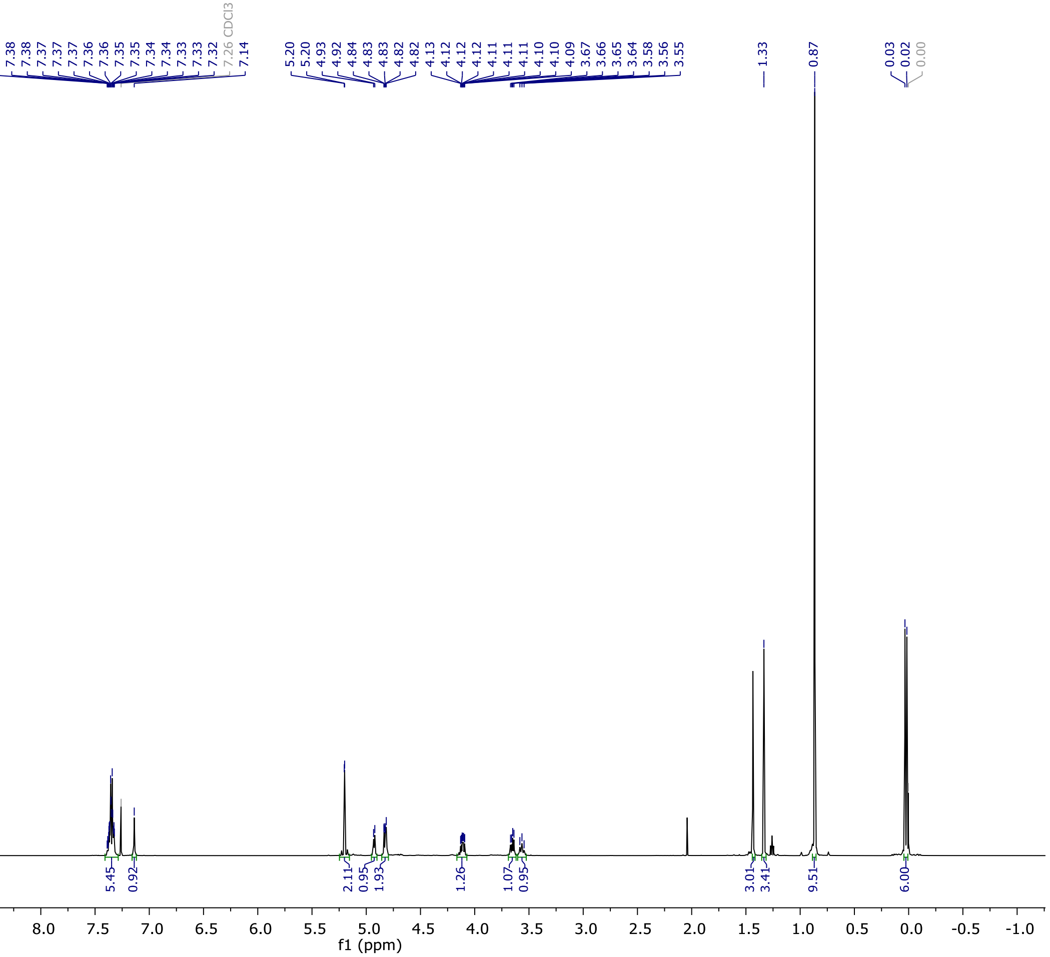

$^{13}C$ NMR (126 MHz, $CDCl_3$)

fe-rsh-av20-047c.11.fid
Owner rsh
Sample RSh-AV20-047c
Platform 3355
Job No fe

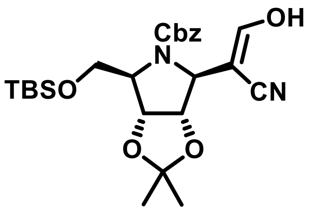

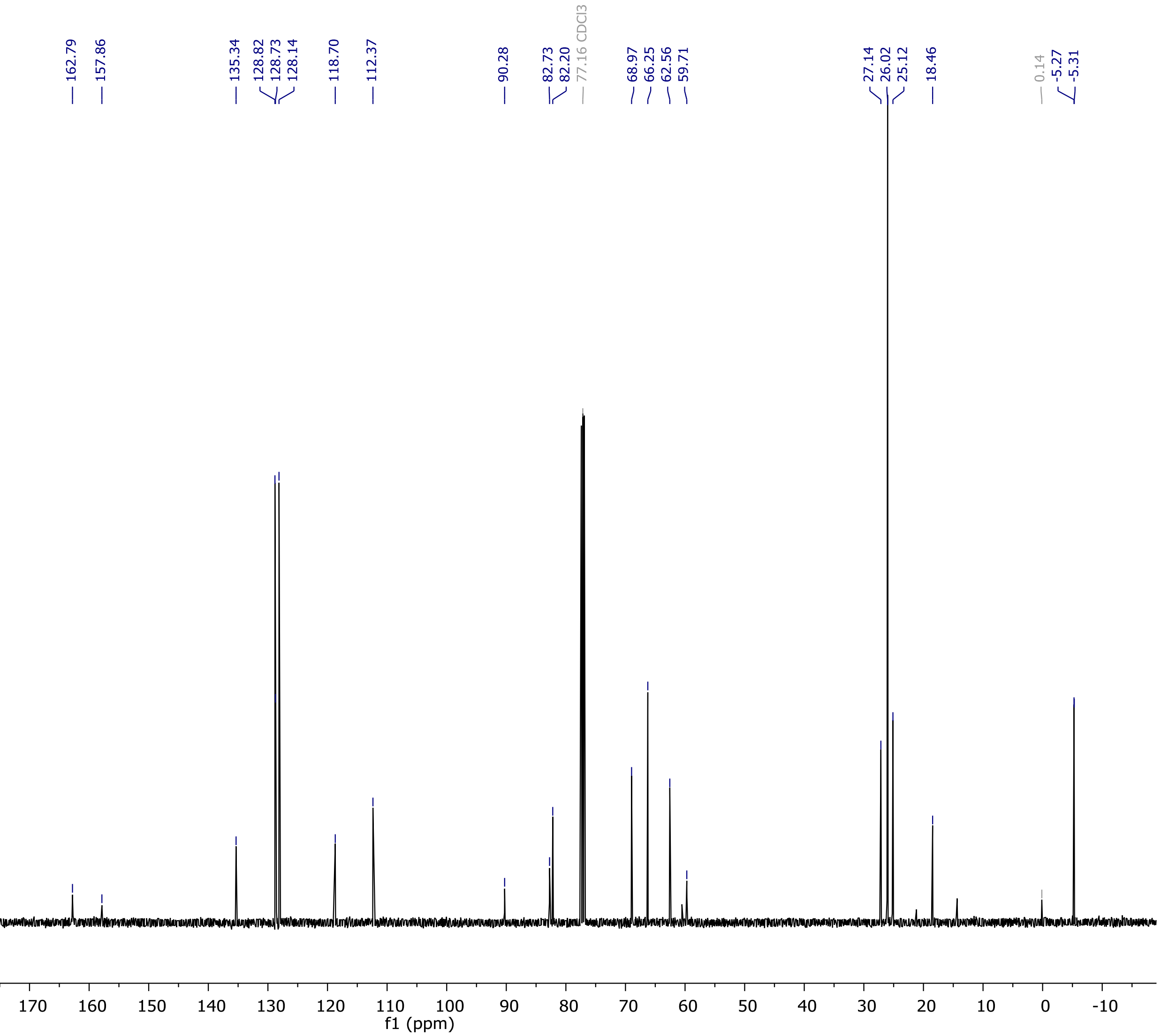

HRMS (ESI-MS)

### Elemental Composition Report

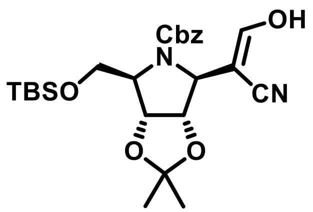

#### Single Mass Analysis

Tolerance = 5.0 mDa / DBE: min = -1.5, max = 150.0
Element prediction: Off
Number of isotope peaks used for i-FIT = 3

Monoisotopic Mass, Even Electron Ions
179 formula(e) evaluated with 2 results within limits (up to 100 closest results for each mass)
Elements Used:
C: 0-80 H: 0-100 N: 2-2 O: 0-20 Na: 0-1 Si: 1-1

RSh-AV20-047c 16 (0.177) Cm (14:17-25:33)

13-Jul-2020
1: TOF MS ES+
6.93e+006

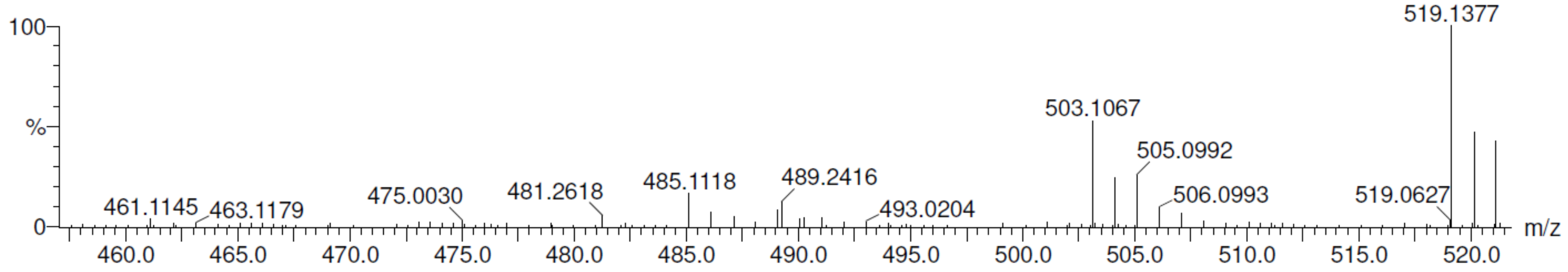

| | | | | | | | | |
|---|---|---|---|---|---|---|---|---|
| Minimum: | | | | -1.5 | | | | |
| Maximum: | | 5.0 | 5.0 | 150.0 | | | | |
| Mass | Calc. Mass | mDa | PPM | DBE | i-FIT | Norm | Conf(%) | Formula |
| 489.2416 | 489.2421 | -0.5 | -1.0 | 9.5 | 87.3 | 0.755 | 46.99 | C25 H37 N2 O6 Si |
| | 489.2397 | 1.9 | 3.9 | 6.5 | 87.2 | 0.635 | 53.01 | C23 H38 N2 O6 Na Si |

(1*S*)-1-(3-Amino-*N*-ethoxycarbonyl-2-cyano-pyrrol-4-yl)-*N*-benzyloxycarbonyl-5-*O*-*tert*-butyldimethylsilyl-1,4-dideoxy-1,4-imino-2,3-*O*-isopropylidene-D-ribitol (**5**)

$^{1}$H NMR (500 MHz, $CDCl_3$)

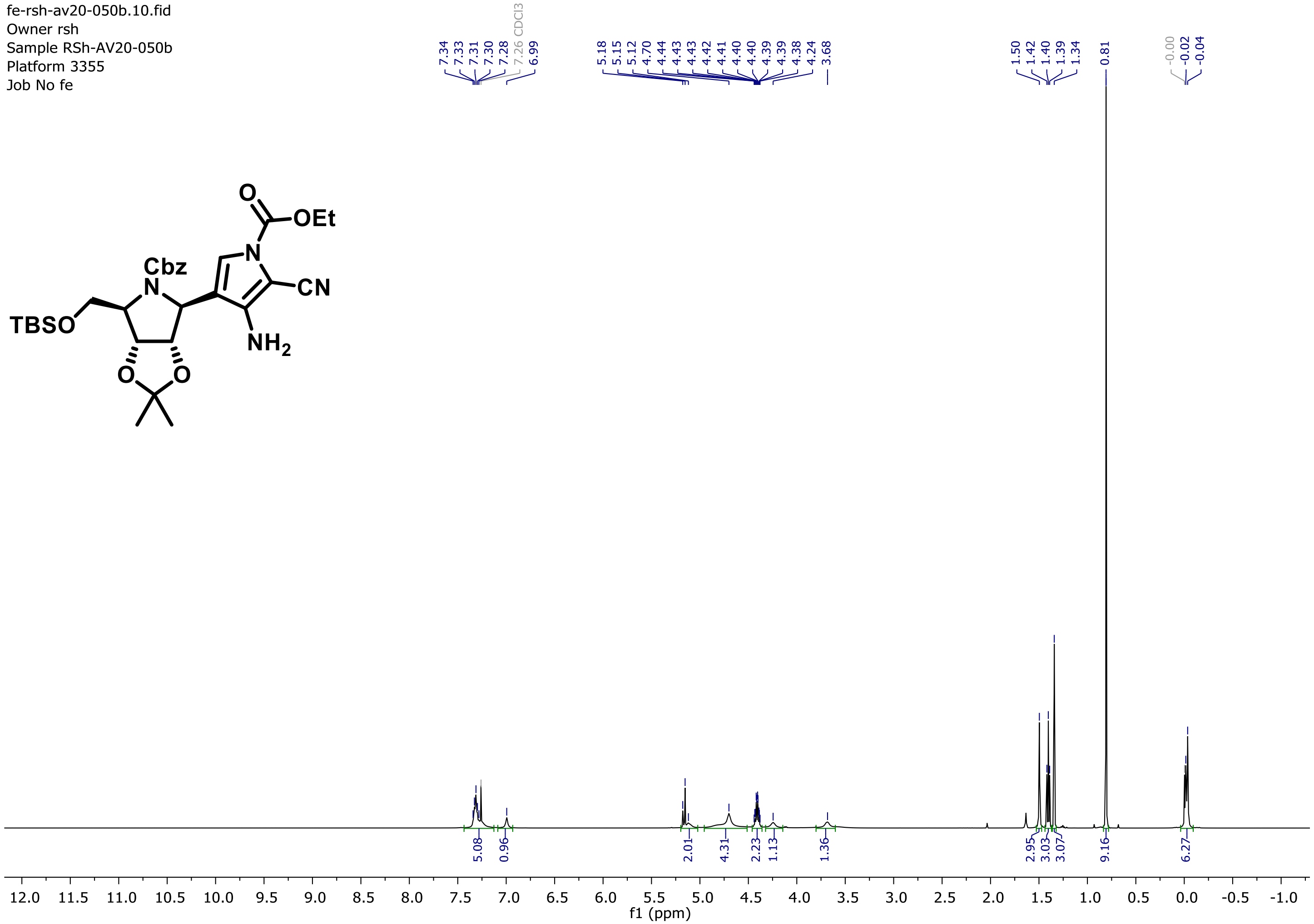

$^{13}C$ NMR (126 MHz, $CDCl_3$)

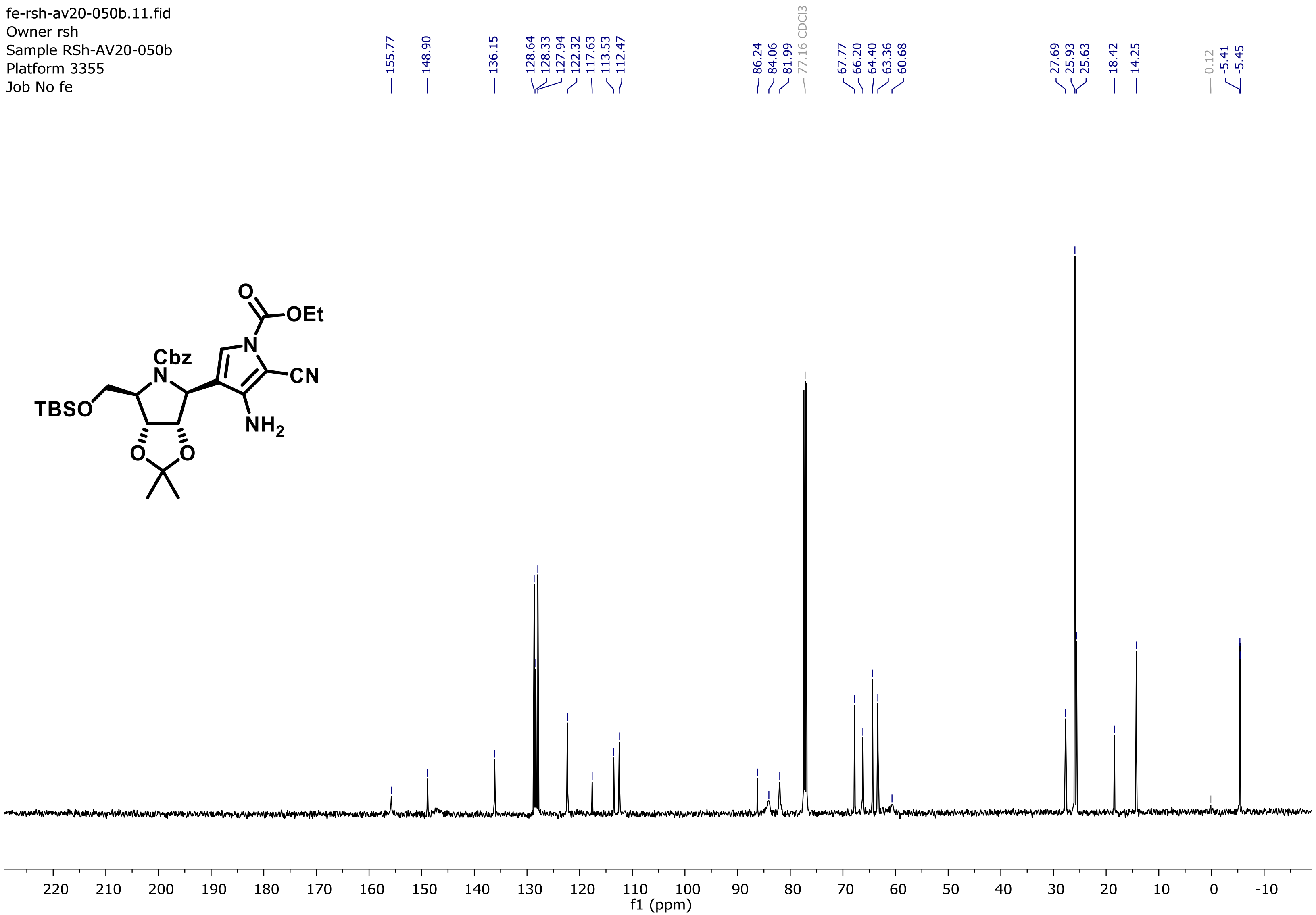

HRMS (ESI-MS)

### Elemental Composition Report

#### Single Mass Analysis

Tolerance = 5.0 mDa / DBE: min = -1.5, max = 150.0
Element prediction: Off
Number of isotope peaks used for i-FIT = 3

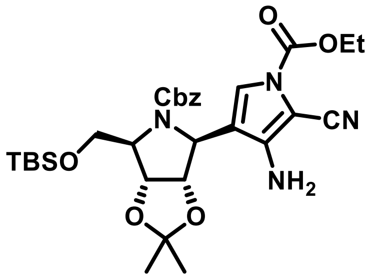

Monoisotopic Mass, Even Electron Ions
456 formula(e) evaluated with 3 results within limits (up to 100 closest results for each mass)
Elements Used:
C: 0-80 H: 0-100 N: 4-5 O: 0-20 Na: 0-1 Si: 1-1

RSh-AV20-050b 24 (0.257) Cm (21:24)

20-Jul-2020
1: TOF MS ES+
7.68e+006

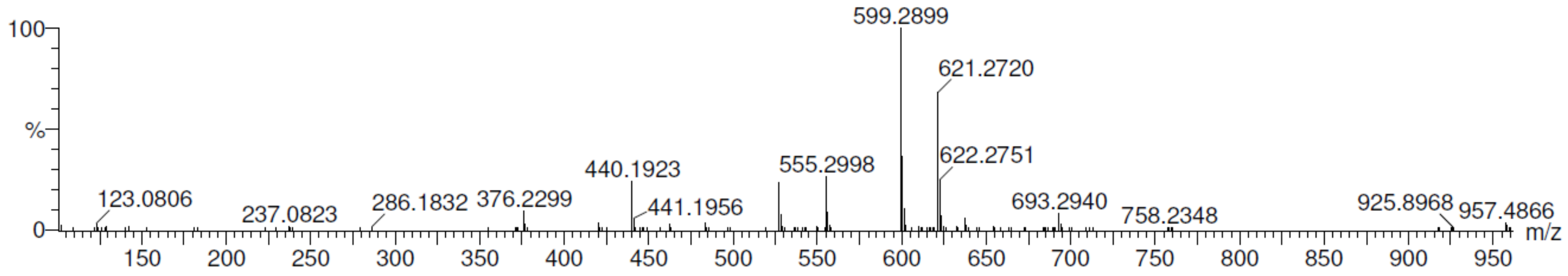

| Mass | Calc. Mass | mDa | PPM | DBE | i-FIT | Norm | Conf(%) | Formula |
|---|---|---|---|---|---|---|---|---|
| Minimum: | | | | -1.5 | | | | |
| Maximum: | | 5.0 | 5.0 | 150.0 | | | | |
| 599.2899 | 599.2901 | -0.2 | -0.3 | 12.5 | 52.0 | 1.044 | 35.19 | C30 H43 N4 O7 Si |
| | 599.2877 | 2.2 | 3.7 | 9.5 | 51.4 | 0.466 | 62.74 | C28 H44 N4 O7 Na Si |
| | 599.2936 | -3.7 | -6.2 | 0.5 | 54.8 | 3.877 | 2.07 | C21 H48 N4 O12 Na Si |

(1*S*)-1-(9-Deazaadenin-9-yl)-*N*-benzyloxycarbonyl-5-*O*-*tert*-butyldimethylsilyl-1,4-dideoxy-1,4-imino-2,3-*O*-isopropylidene-D-ribitol (**6**)

$^1$H NMR (500 MHz, $CDCl_3$)

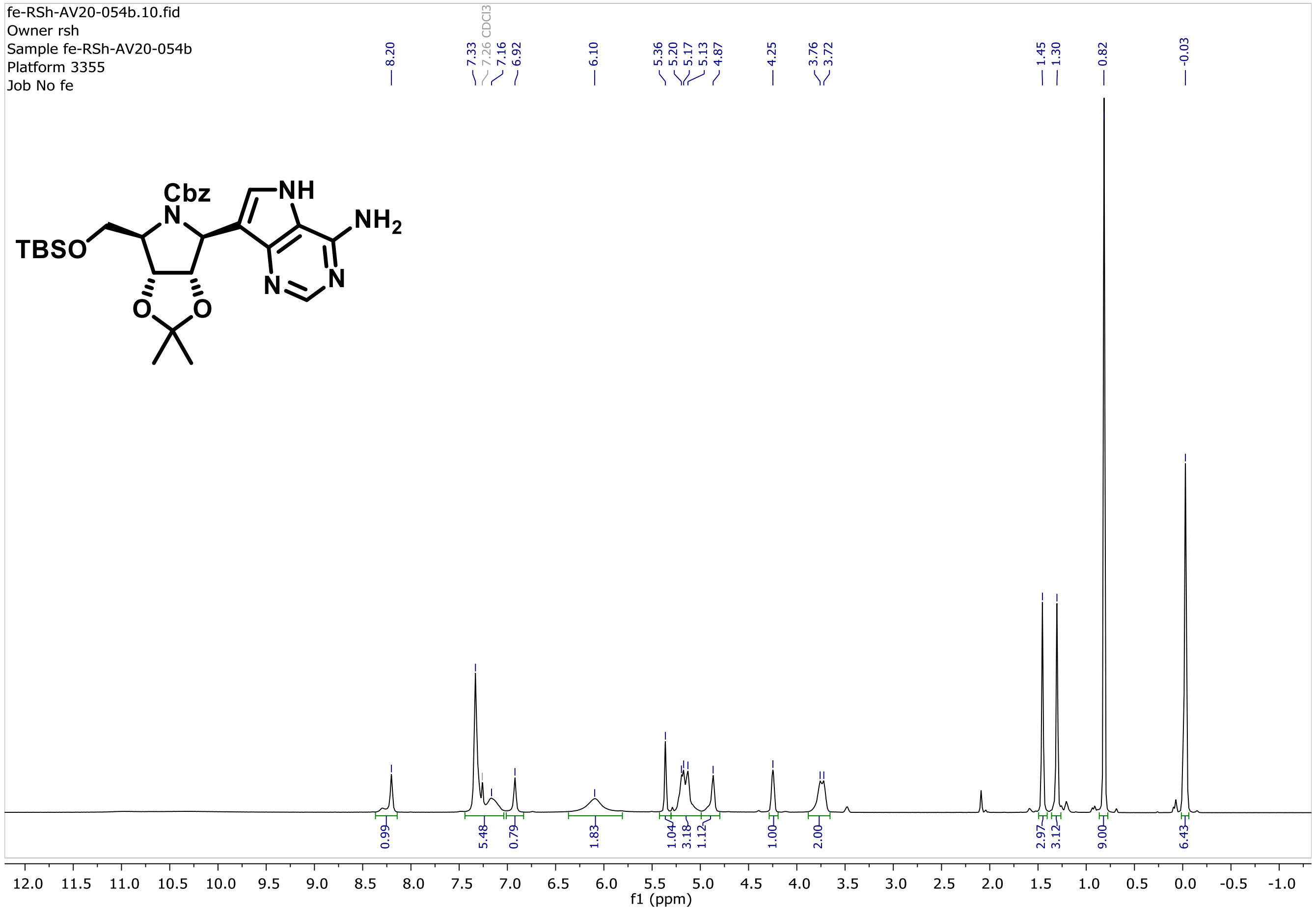

$^{13}C$ NMR (126 MHz, $CDCl_3$)

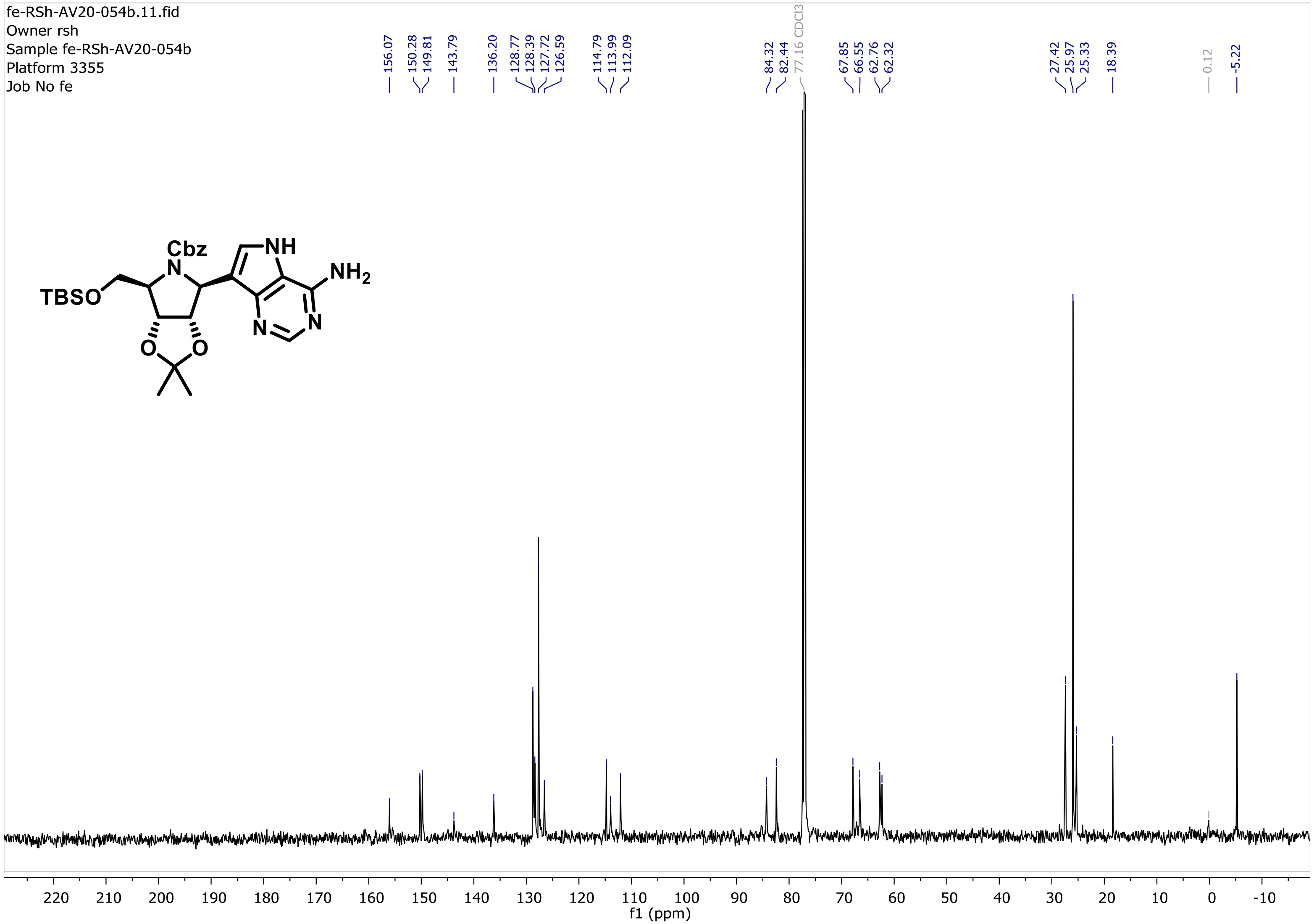

HRMS (ESI-MS)

### Elemental Composition Report

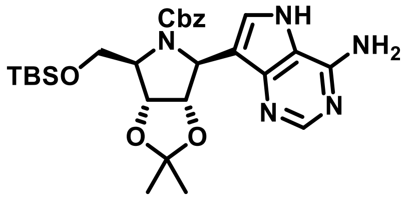

#### Single Mass Analysis

Tolerance = 5.0 mDa / DBE: min = -1.5, max = 150.0
Element prediction: Off
Number of isotope peaks used for i-FIT = 3

Monoisotopic Mass, Even Electron Ions
203 formula(e) evaluated with 3 results within limits (up to 100 closest results for each mass)
Elements Used:
C: 0-80 H: 0-100 N: 5-5 O: 0-20 Na: 0-1 Si: 1-1

RSh-AV20-054b 32 (0.337)

20-Jul-2020
1: TOF MS ES+
9.89e+006

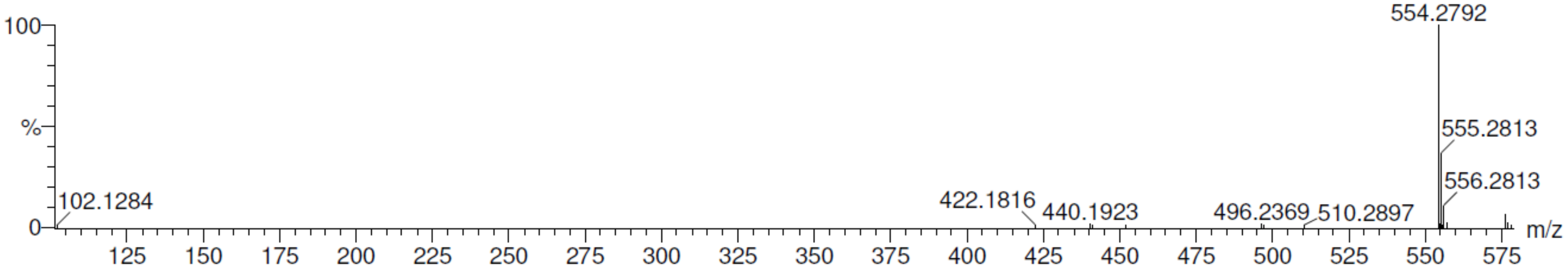

| | | mDa | PPM | DBE | | | | |
|---|---|---|---|---|---|---|---|---|
| Minimum: | | | | -1.5 | | | | |
| Maximum: | | 5.0 | 5.0 | 150.0 | | | | |
| **Mass** | **Calc. Mass** | **mDa** | **PPM** | **DBE** | **i-FIT** | **Norm** | **Conf(%)** | **Formula** |
| 554.2792 | 554.2799 | -0.7 | -1.3 | 12.5 | 98.8 | 3.936 | 1.95 | C28 H40 N5 O5 Si |
| | 554.2775 | 1.7 | 3.1 | 9.5 | 94.9 | 0.020 | 98.05 | C26 H41 N5 O5 Na Si |
| | 554.2833 | -4.1 | -7.4 | 0.5 | 105.9 | 11.047 | 0.00 | C19 H45 N5 O10 Na Si |

(1*S*)-1-(9-Deazaadenin-9-yl)-*N*-benzyloxycarbonyl-1,4-dideoxy-1,4-imino-2,3-*O*-isopropylidene-D-ribitol (**7**)

$^1$H NMR (500 MHz, $CDCl_3$)

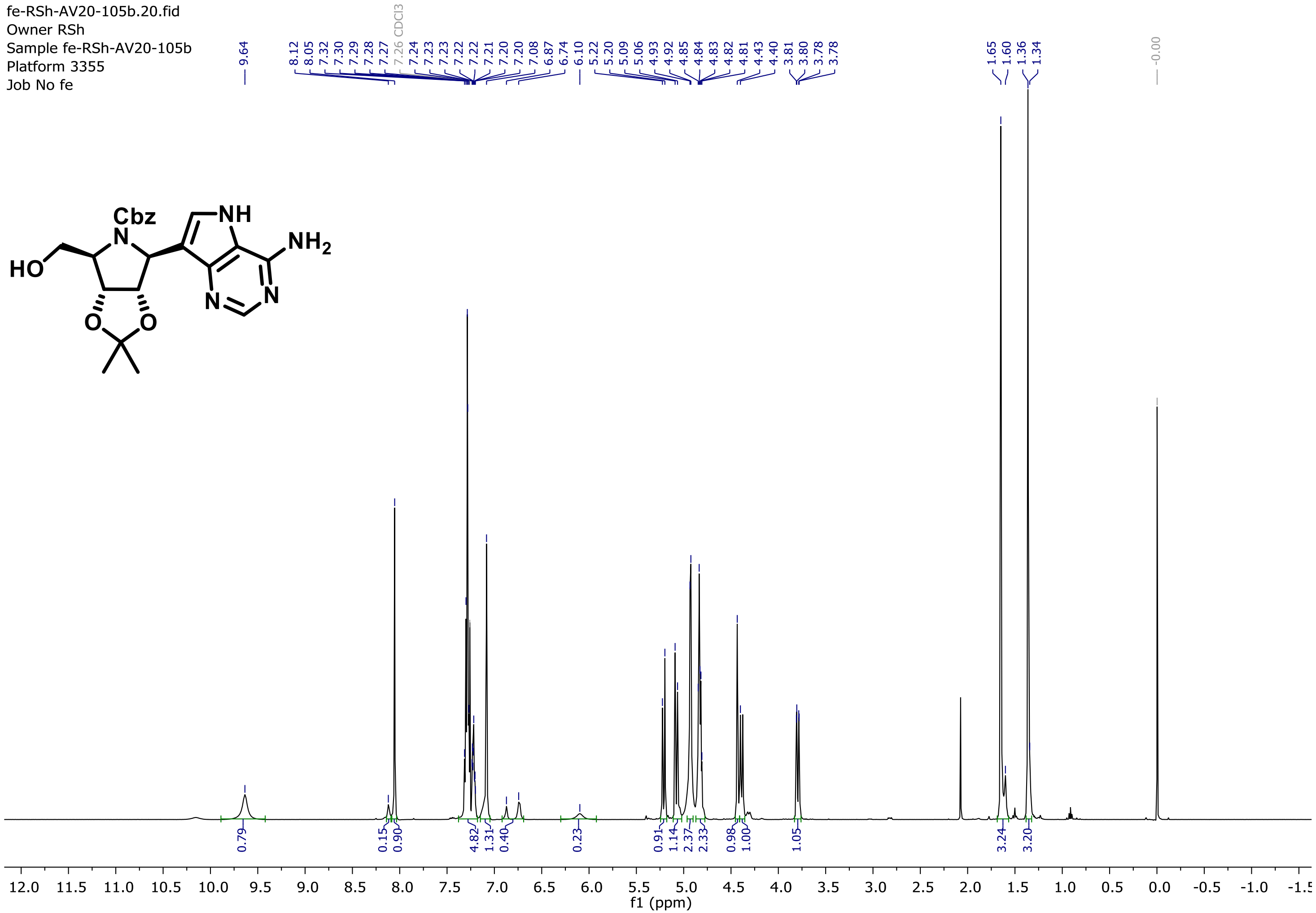

$^{13}C$ NMR (126 MHz, $CDCl_3$)

fe-RSh-AV20-105b.21.fid
Owner RSh
Sample fe-RSh-AV20-105b
Platform 3355
Job No fe

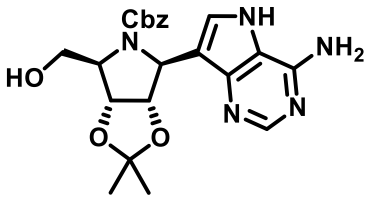

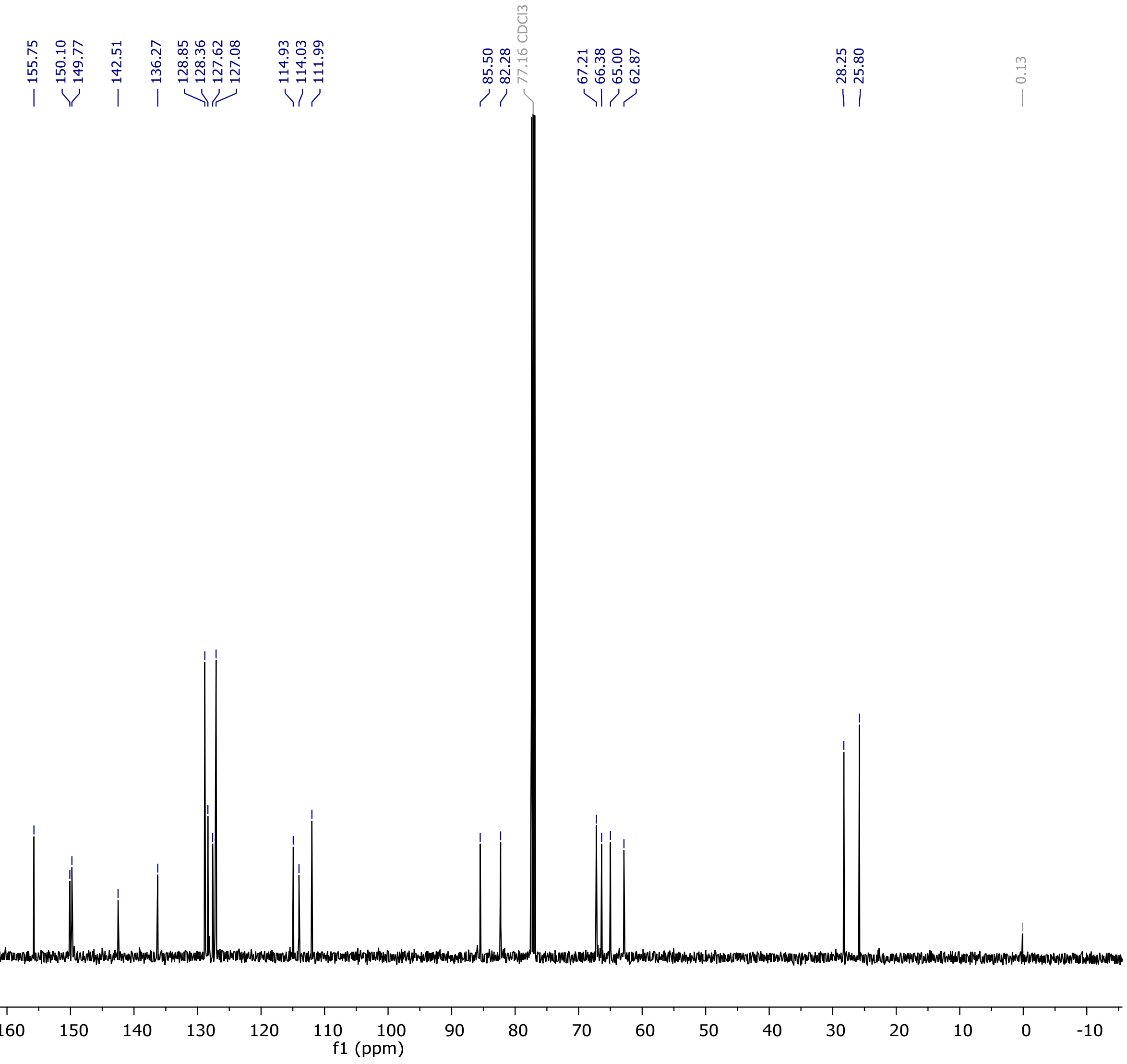

HRMS (ESI-MS)

### Elemental Composition Report

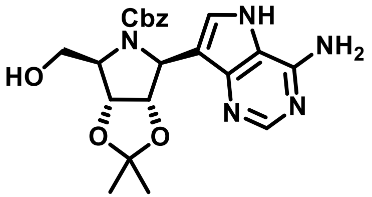

#### Single Mass Analysis

Tolerance = 5.0 mDa / DBE: min = -1.5, max = 150.0
Element prediction: Off
Number of isotope peaks used for i-FIT = 3

Monoisotopic Mass, Even Electron Ions
154 formula(e) evaluated with 3 results within limits (up to 100 closest results for each mass)
Elements Used:
C: 0-80 H: 0-120 N: 5-5 O: 0-20 Na: 0-1

RSh-AV20-105b 21 (0.231) Cm (17:22-1:7)

25-Nov-2020
1: TOF MS ES+
2.73e+007

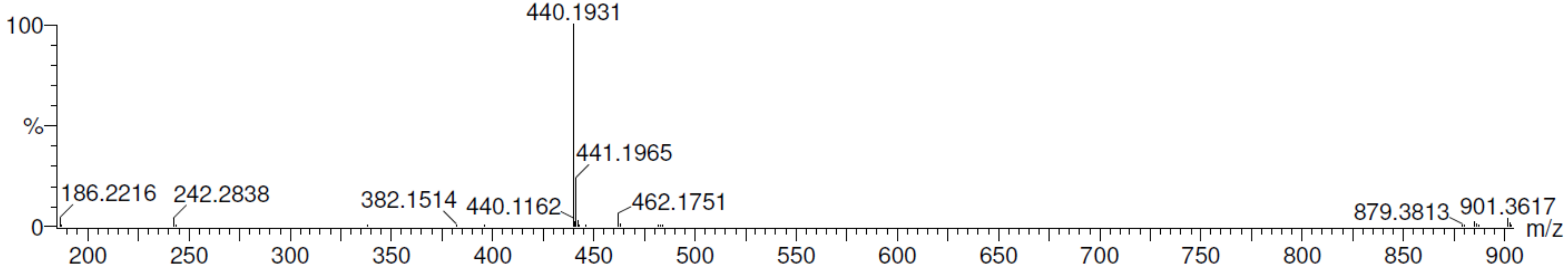

| Mass | Calc. Mass | mDa | PPM | DBE | i-FIT | Norm | Conf(%) | Formula |
|---|---|---|---|---|---|---|---|---|
| Minimum: | | | | -1.5 | | | | |
| Maximum: | | 5.0 | 5.0 | 150.0 | | | | |
| 440.1931 | 440.1934 | -0.3 | -0.7 | 12.5 | 86.5 | 1.018 | 36.14 | C22 H26 N5 O5 |
| | 440.1910 | 2.1 | 4.8 | 9.5 | 86.0 | 0.456 | 63.37 | C20 H27 N5 O5 Na |
| | 440.1969 | -3.8 | -8.6 | 0.5 | 90.8 | 5.309 | 0.49 | C13 H31 N5 O10 Na |

Compound **8**

$^1$H NMR (500 MHz, MeOD)

fe-RSh-AV20-067c.10.fid
Owner RSh
Sample fe-RSh-AV20-067c
Platform 3355
Job No fe

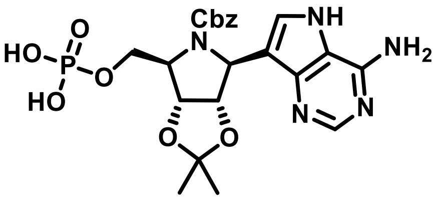

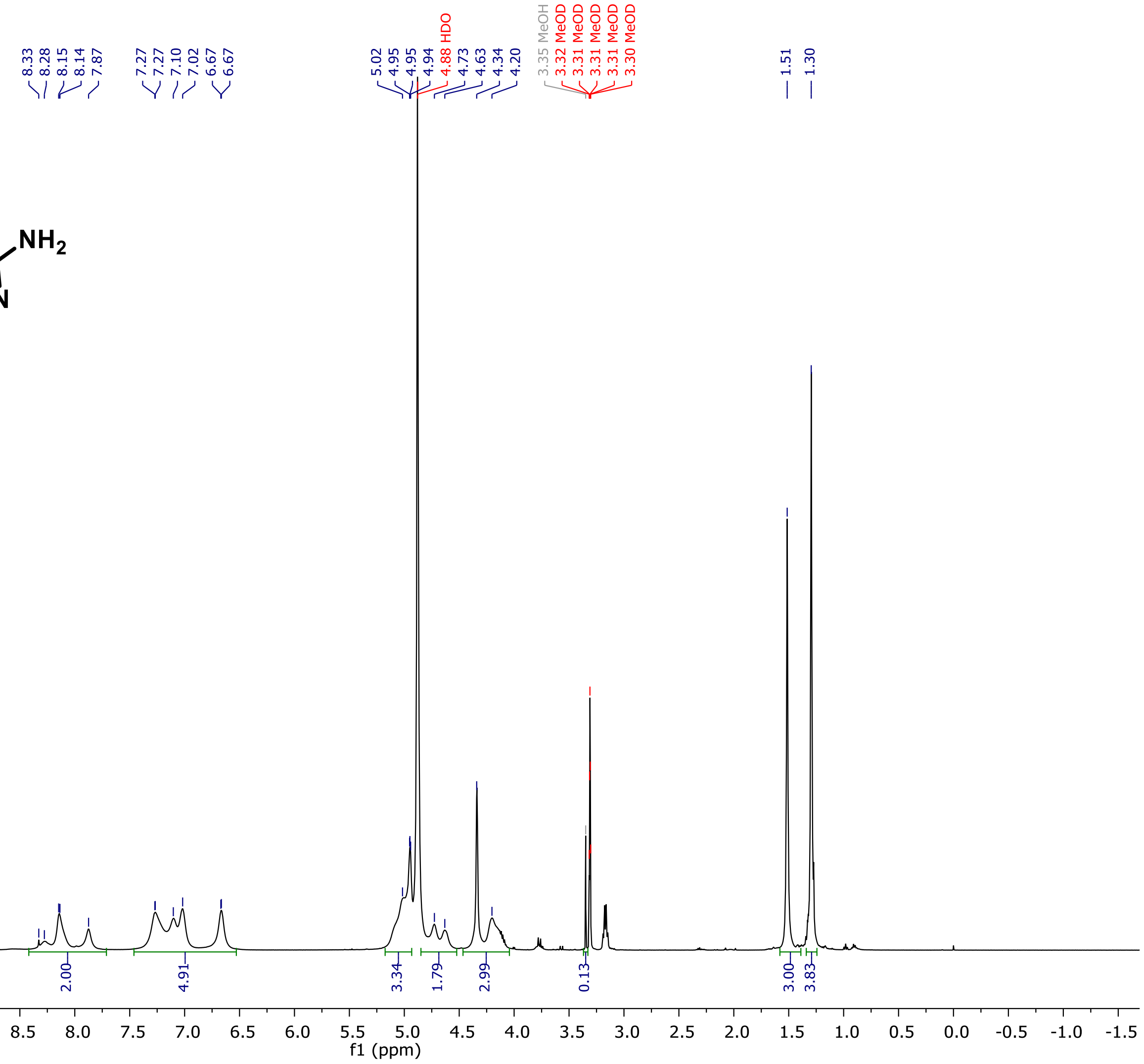

$^{13}$C NMR (126 MHz, MeOD)

fe-RSh-AV20-067c.11.fid
Owner RSh
Sample fe-RSh-AV20-067c
Platform 3355
Job No fe

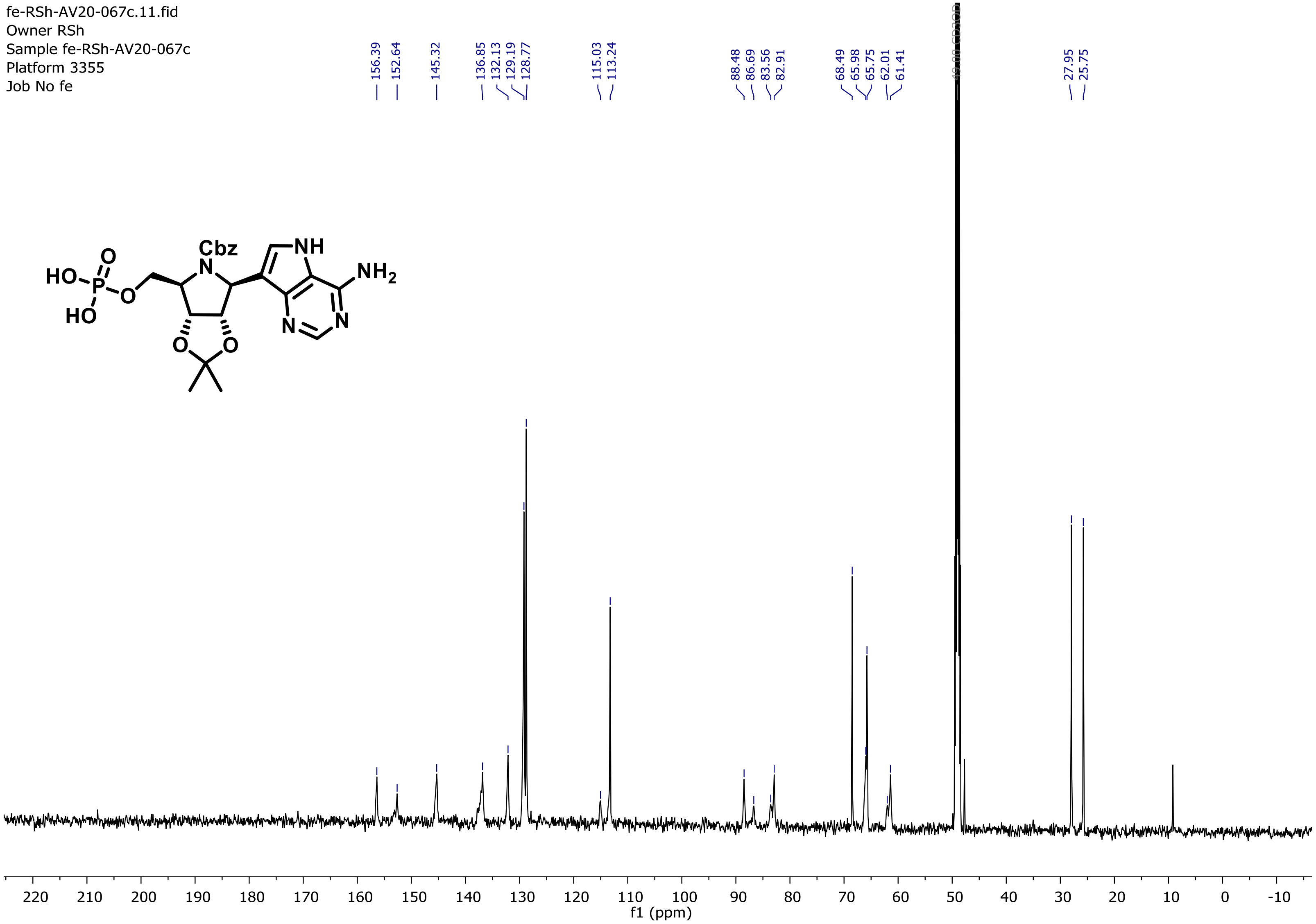

$^{31}$P NMR (202 MHz, MeOD)

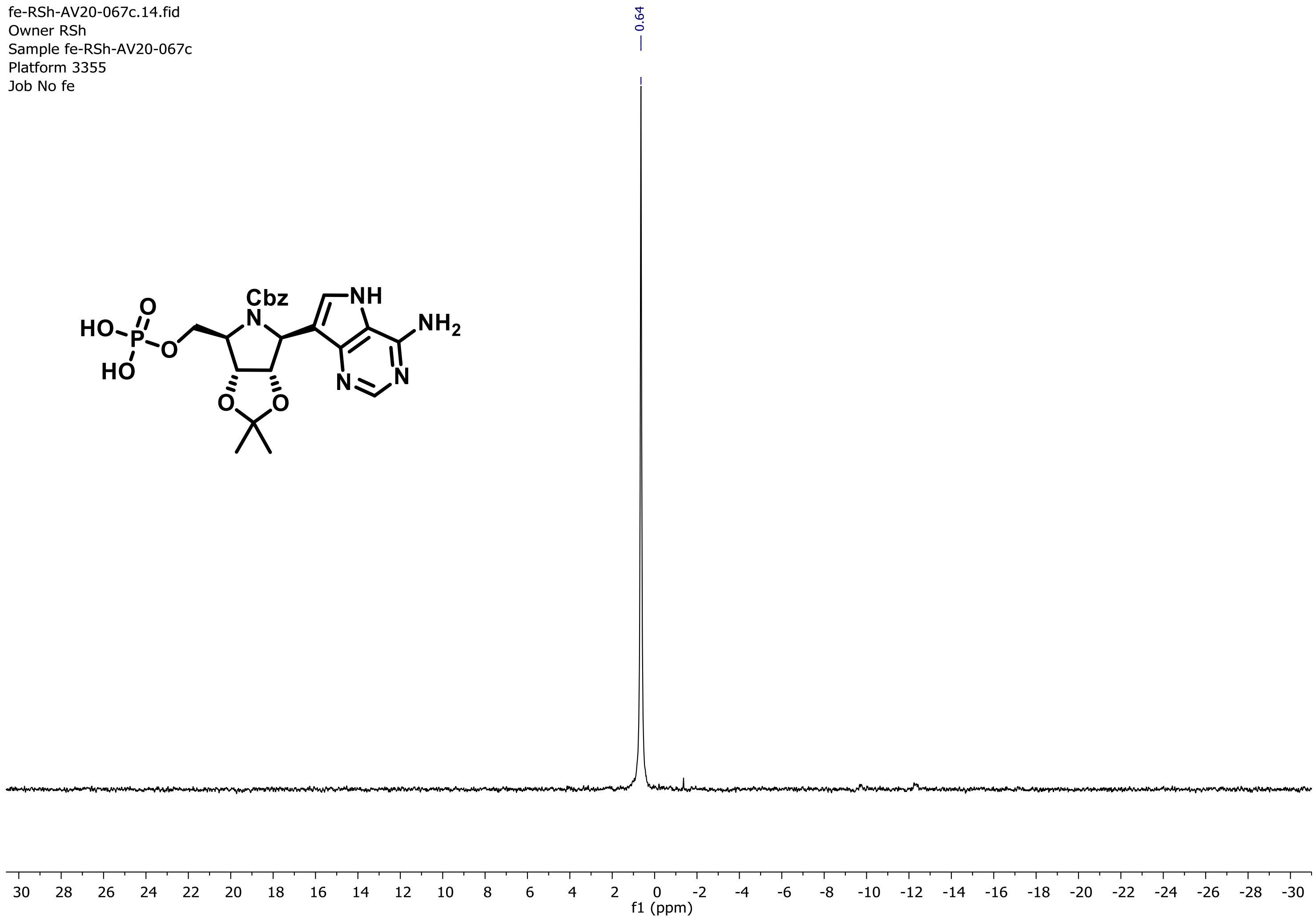

HRMS (ESI-MS)

#### Elemental Composition Report

##### Single Mass Analysis

Tolerance = 5.0 mDa / DBE: min = -1.5, max = 150.0
Element prediction: Off
Number of isotope peaks used for i-FIT = 3

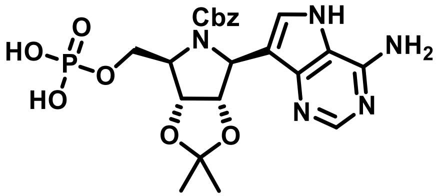

Monoisotopic Mass, Even Electron Ions
194 formula(e) evaluated with 3 results within limits (up to 100 closest results for each mass)
Elements Used:
C: 0-120 H: 0-120 N: 5-5 O: 0-40 Na: 0-1 P: 1-1

RSh-AV20-067c- 18 (0.171) AM2 (Ar,22000.0,248.96,0.00); ABS; Cm (15:20)

04-Aug-2020
TOF MS ES-
2.16e+007

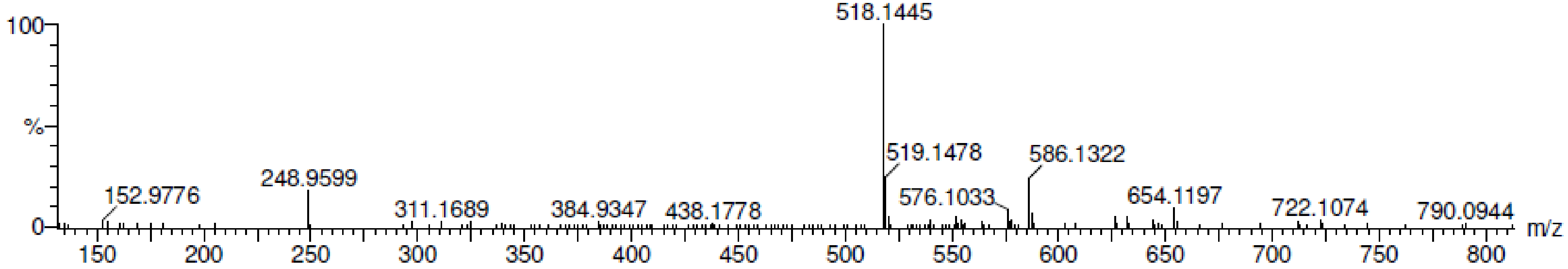

| | | | | | | | | |
|---|---|---|---|---|---|---|---|---|
| Minimum: | | | | -1.5 | | | | |
| Maximum: | | 5.0 | 5.0 | 150.0 | | | | |
| Mass | Calc. Mass | mDa | PPM | DBE | i-FIT | Norm | Conf(%) | Formula |
| 518.1445 | 518.1441 | 0.4 | 0.8 | 13.5 | 59.3 | 6.570 | 0.14 | C22 H25 N5 O8 P |
| | 518.1417 | 2.8 | 5.4 | 10.5 | 52.8 | 0.001 | 99.86 | C20 H26 N5 O8 Na P |
| | 518.1475 | -3.0 | -5.8 | 1.5 | 65.1 | 12.300 | 0.00 | C13 H30 N5 O13 Na P |

Compound **9**

$^{1}H$ NMR (500 MHz, $D_2O$)

fe-RSh-AV20-072b.10.fid
Owner RSh
Sample fe-RSh-AV20-072b
Platform 3355
Job No fe

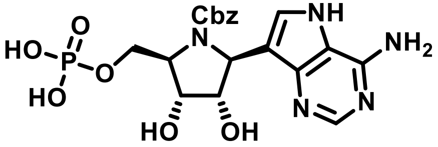

**(triethylammonium salt)**

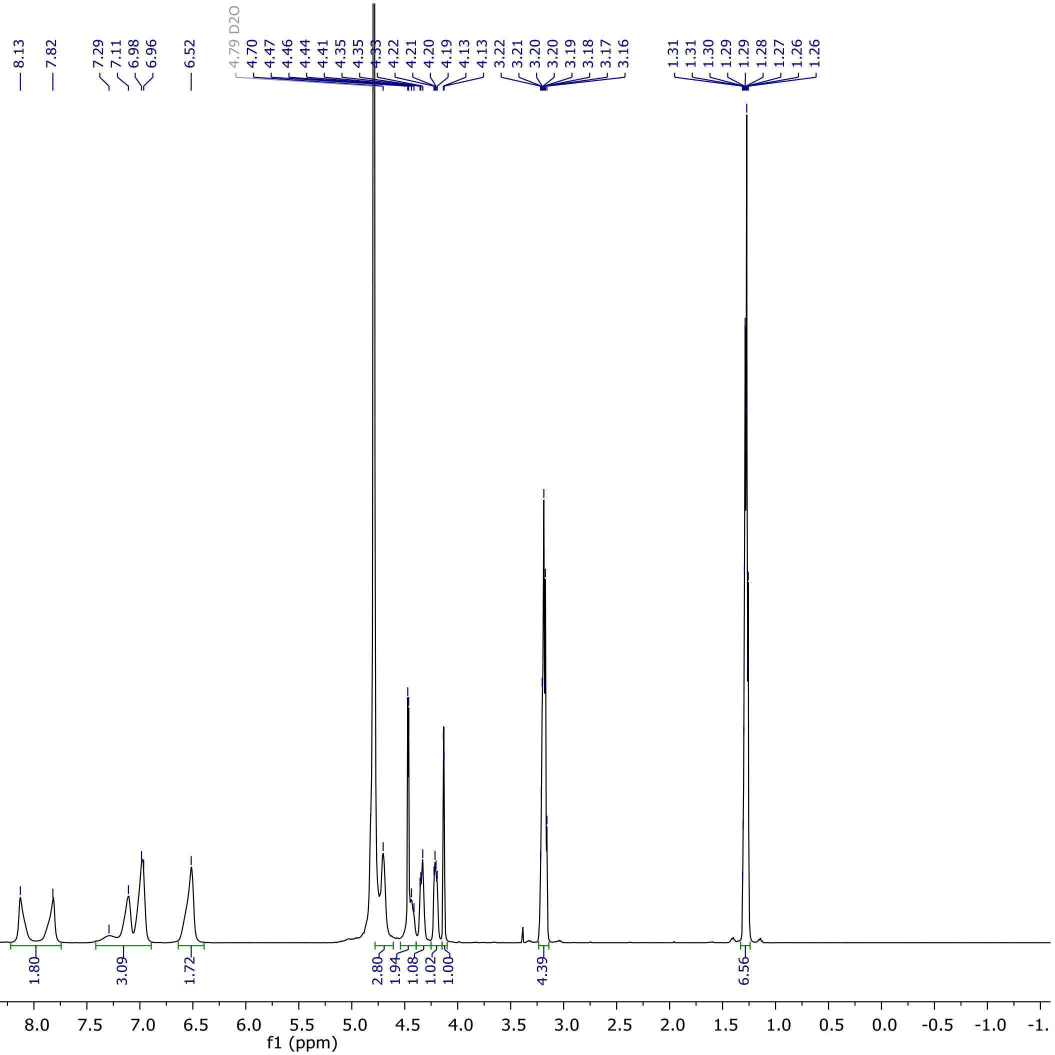

$^{13}C$ NMR (126 MHz, $D_2O$)

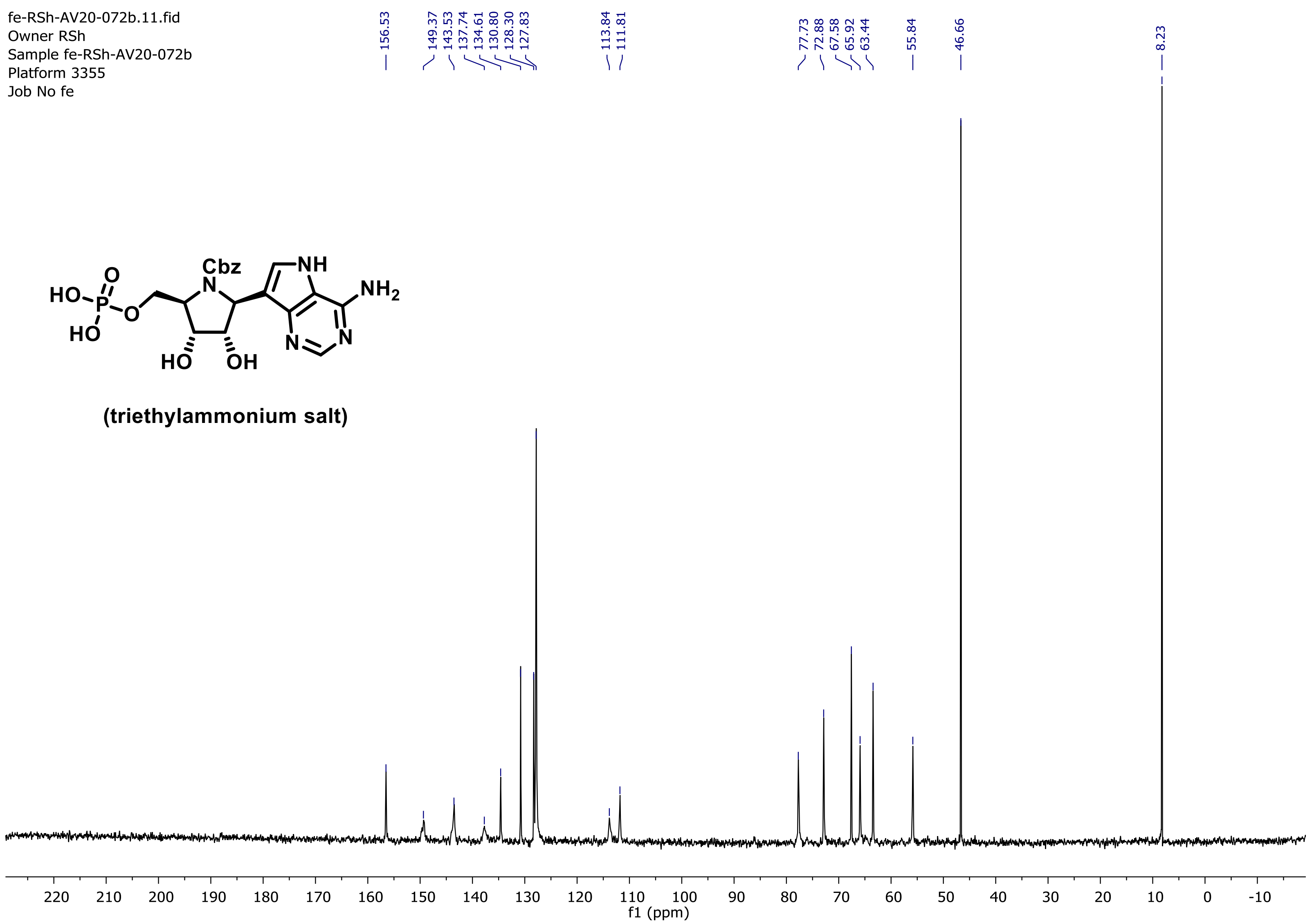

$^{31}P$ NMR (202 MHz, $D_2O$)

fe-RSh-AV20-072b.15.fid
Owner RSh
Sample fe-RSh-AV20-072b
Platform 3355
Job No fe

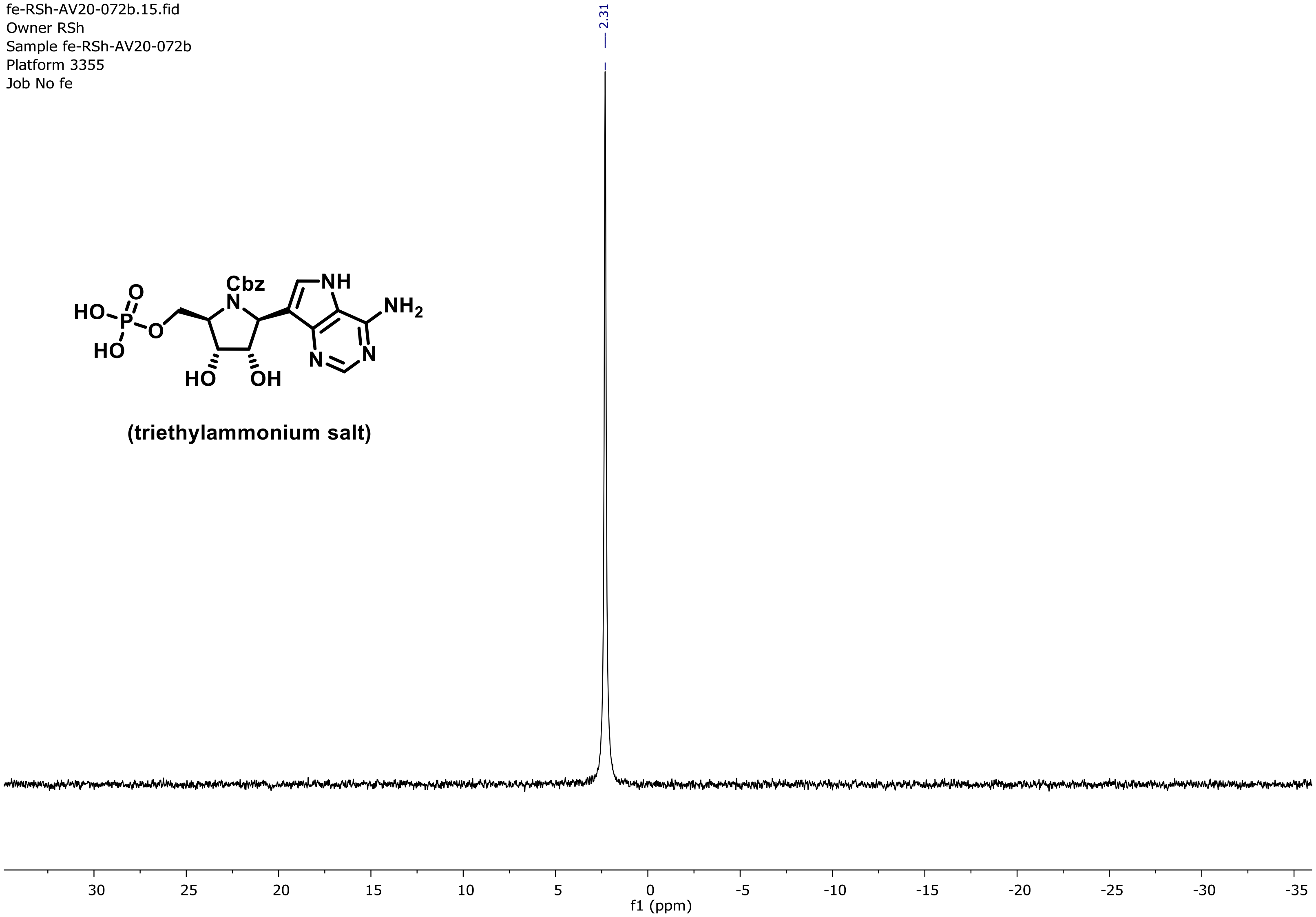

HRMS (ESI-MS)

### Elemental Composition Report

#### Single Mass Analysis

Tolerance = 5.0 mDa / DBE: min = -1.5, max = 150.0
Element prediction: Off
Number of isotope peaks used for i-FIT = 3

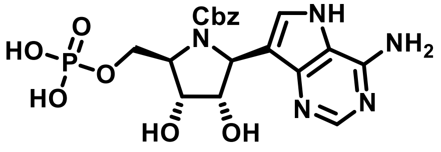

**(triethylammonium salt)**

Monoisotopic Mass, Even Electron Ions
164 formula(e) evaluated with 3 results within limits (up to 100 closest results for each mass)
Elements Used:
C: 0-120 H: 0-120 N: 5-5 O: 0-40 Na: 0-1 P: 1-1

RSh-AV20-072b- 9 (0.117) Cm (9:11)

10-Aug-2020
1: TOF MS ES-
1.44e+007

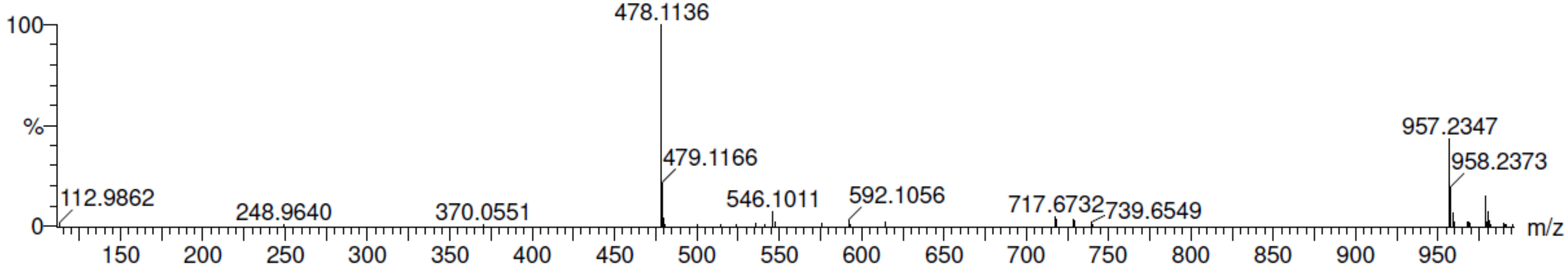

| Mass | Calc. Mass | mDa | PPM | DBE | i-FIT | Norm | Conf(%) | Formula |
|---|---|---|---|---|---|---|---|---|
| Minimum: | | | | -1.5 | | | | |
| Maximum: | | 5.0 | 5.0 | 150.0 | | | | |
| 478.1136 | 478.1128 | 0.8 | 1.7 | 12.5 | 101.9 | 5.016 | 0.66 | C19 H21 N5 O8 P |
| | 478.1162 | -2.6 | -5.4 | 0.5 | 109.0 | 12.127 | 0.00 | C10 H26 N5 O13 Na P |
| | 478.1104 | 3.2 | 6.7 | 9.5 | 96.9 | 0.007 | 99.34 | C17 H22 N5 O8 Na P |

Compound **10**

$^1$H NMR (500 MHz, $D_2O$)

fe-RSh-AV20-077e.10.fid
Owner rsh
Sample RSh-AV20-077e
Platform 3355
Job No fe

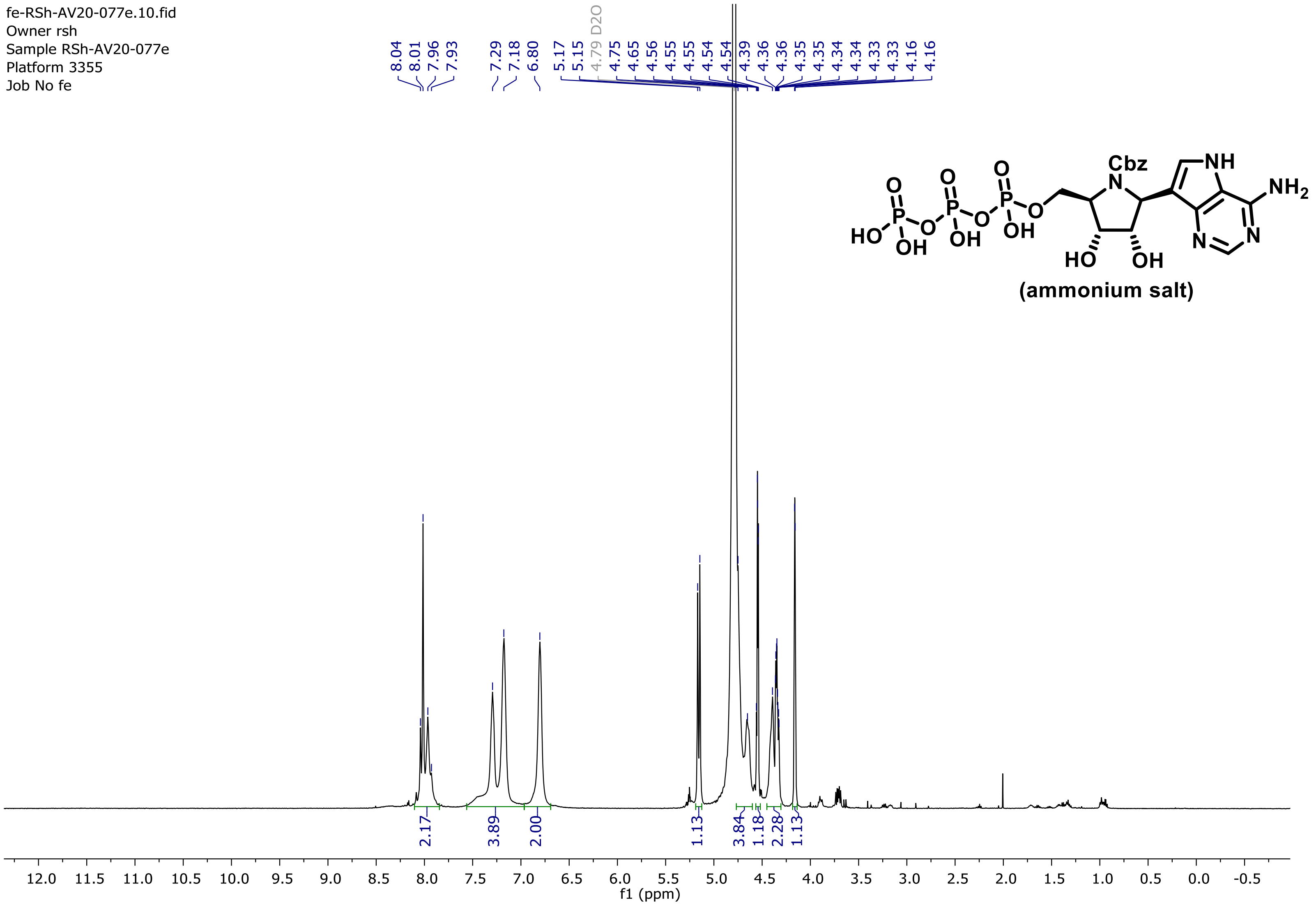

$^{13}C$ NMR (126 MHz, $D_2O$)

fe-RSh-AV20-077e.11.fid
Owner rsh
Sample RSh-AV20-077e
Platform 3355
Job No fe

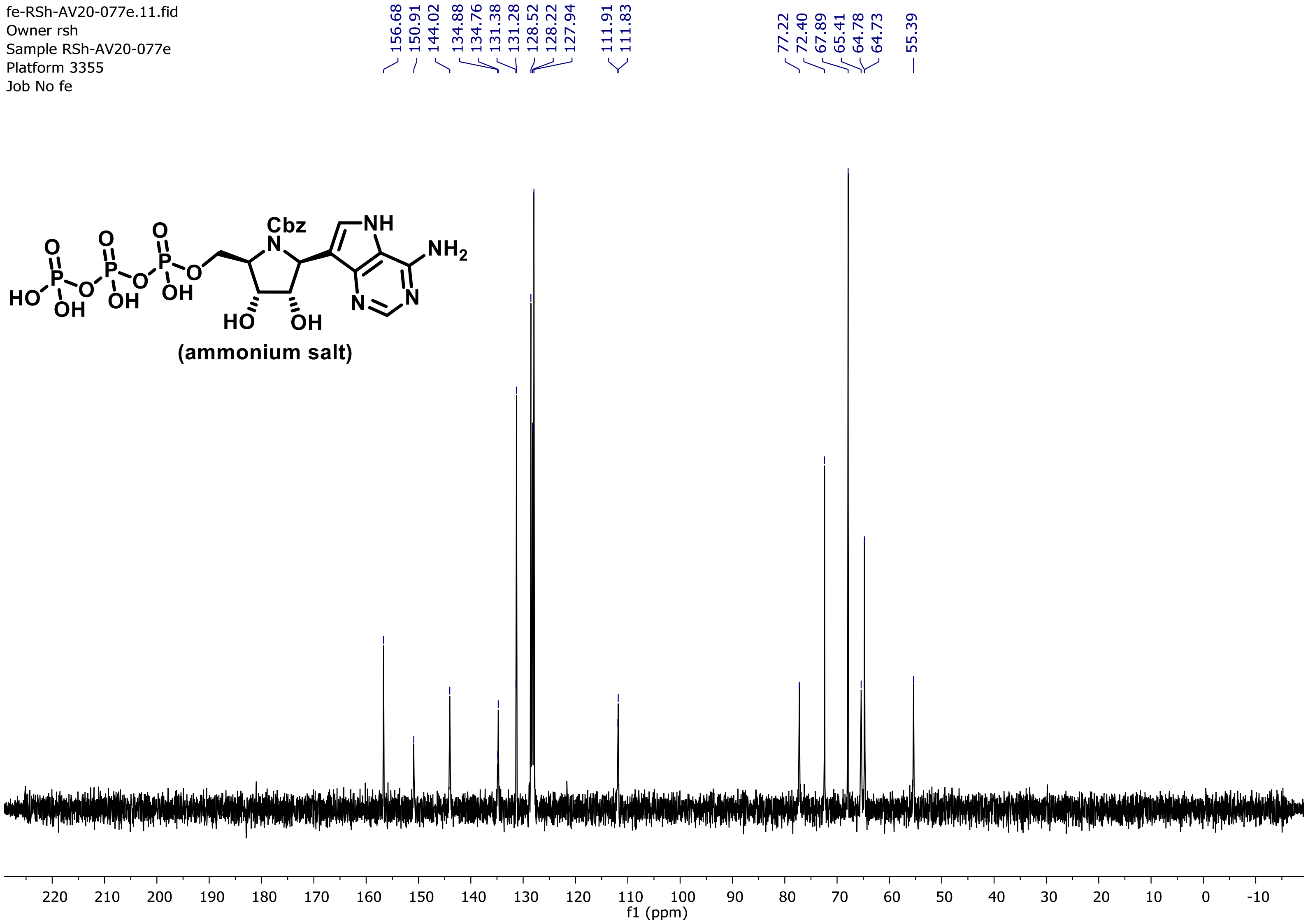

$^{31}P$ NMR (202 MHz, $D_2O$)

fe-RSh-AV20-077e.11.fid
Owner RSh
Sample fe-RSh-AV20-077e
Platform 3355
Job No fe

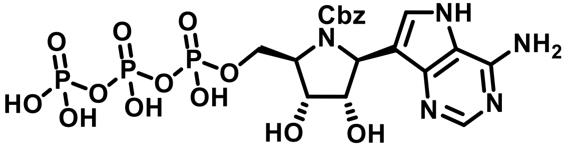

(ammonium salt)

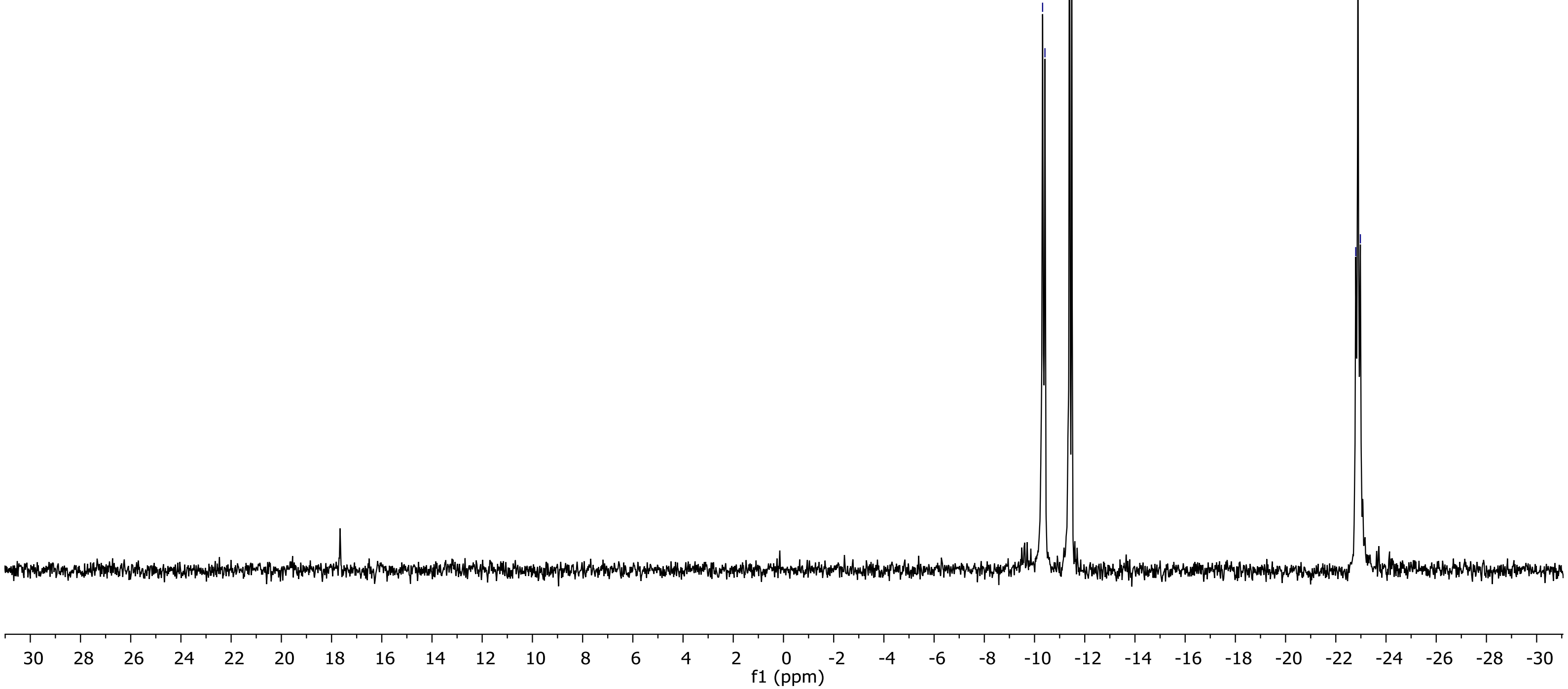

HRMS (ESI-MS)

### Elemental Composition Report

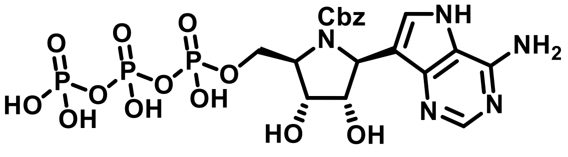

#### Single Mass Analysis

Tolerance = 5.0 mDa / DBE: min = -1.5, max = 150.0
Element prediction: Off
Number of isotope peaks used for i-FIT = 3

Monoisotopic Mass, Even Electron Ions
251 formula(e) evaluated with 6 results within limits (up to 100 closest results for each mass)
Elements Used:
C: 0-120 H: 0-150 N: 5-5 O: 0-40 Na: 0-1 P: 3-3

RSh-AV20-077c 14 (0.160) Cm (8:22-1:6)

19-Aug-2020
1: TOF MS ES+
1.31e+008

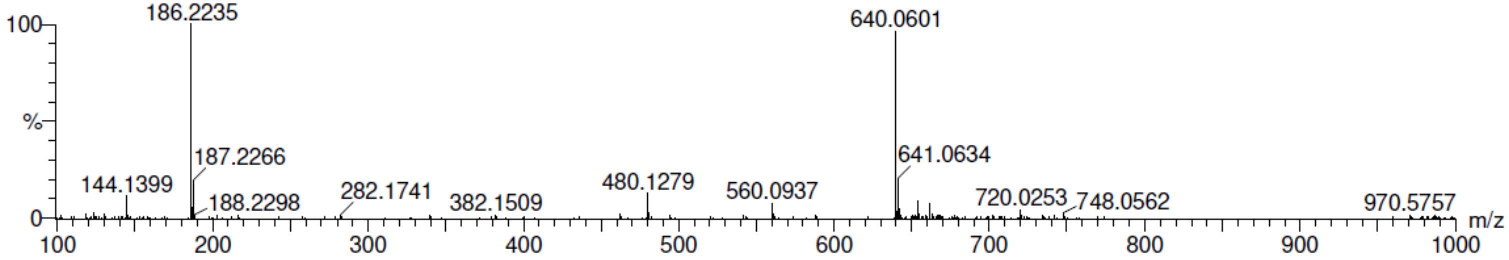

| | | | | | | | | |
|---|---|---|---|---|---|---|---|---|
| Minimum: | | | | -1.5 | | | | |
| Maximum: | | 5.0 | 5.0 | 150.0 | | | | |
| Mass | Calc. Mass | mDa | PPM | DBE | i-FIT | Norm | Conf(%) | Formula |
| 640.0601 | 640.0611 | -1.0 | -1.6 | 11.5 | 144.9 | 7.249 | 0.07 | C19 H25 N5 O14 P3 |
| | 640.0587 | 1.4 | 2.2 | 8.5 | 137.7 | 0.001 | 99.93 | C17 H26 N5 O14 Na P3 |
| | 640.0622 | -2.1 | -3.3 | 30.5 | 154.8 | 17.106 | 0.00 | C35 H18 N5 O Na P3 |
| | 640.0646 | -4.5 | -7.0 | 33.5 | 156.1 | 18.372 | 0.00 | C37 H17 N5 O P3 |
| | 640.0646 | -4.5 | -7.0 | -0.5 | 150.8 | 13.076 | 0.00 | C10 H30 N5 O19 Na P3 |
| | 640.0552 | 4.9 | 7.7 | 20.5 | 152.6 | 14.878 | 0.00 | C26 H21 N5 O9 P3 |

Compound **11**

$^1$H NMR (500 MHz, $D_2O$)

fe-RSh-AV20-080b.20.fid
Owner RSh
Sample fe-RSh-AV20-080b
Platform 3355
Job No fe

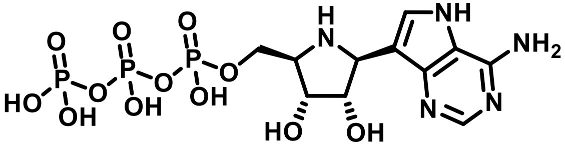

(2 $Bu_3N$ salt)

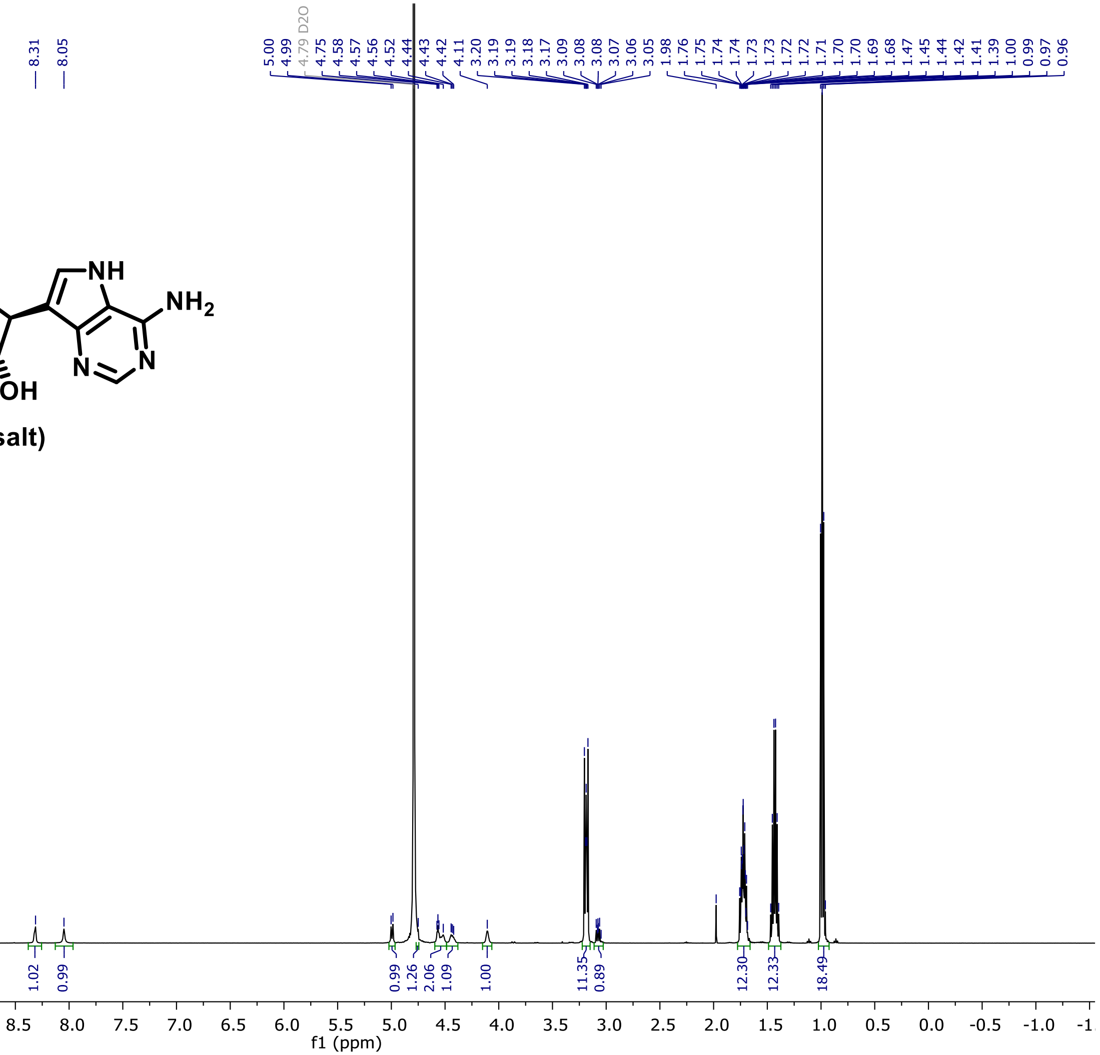

$^{13}C$ NMR (126 MHz, $D_2O$)

fe-RSh-AV20-080b.21.fid
Owner RSh
Sample fe-RSh-AV20-080b
Platform 3355
Job No fe

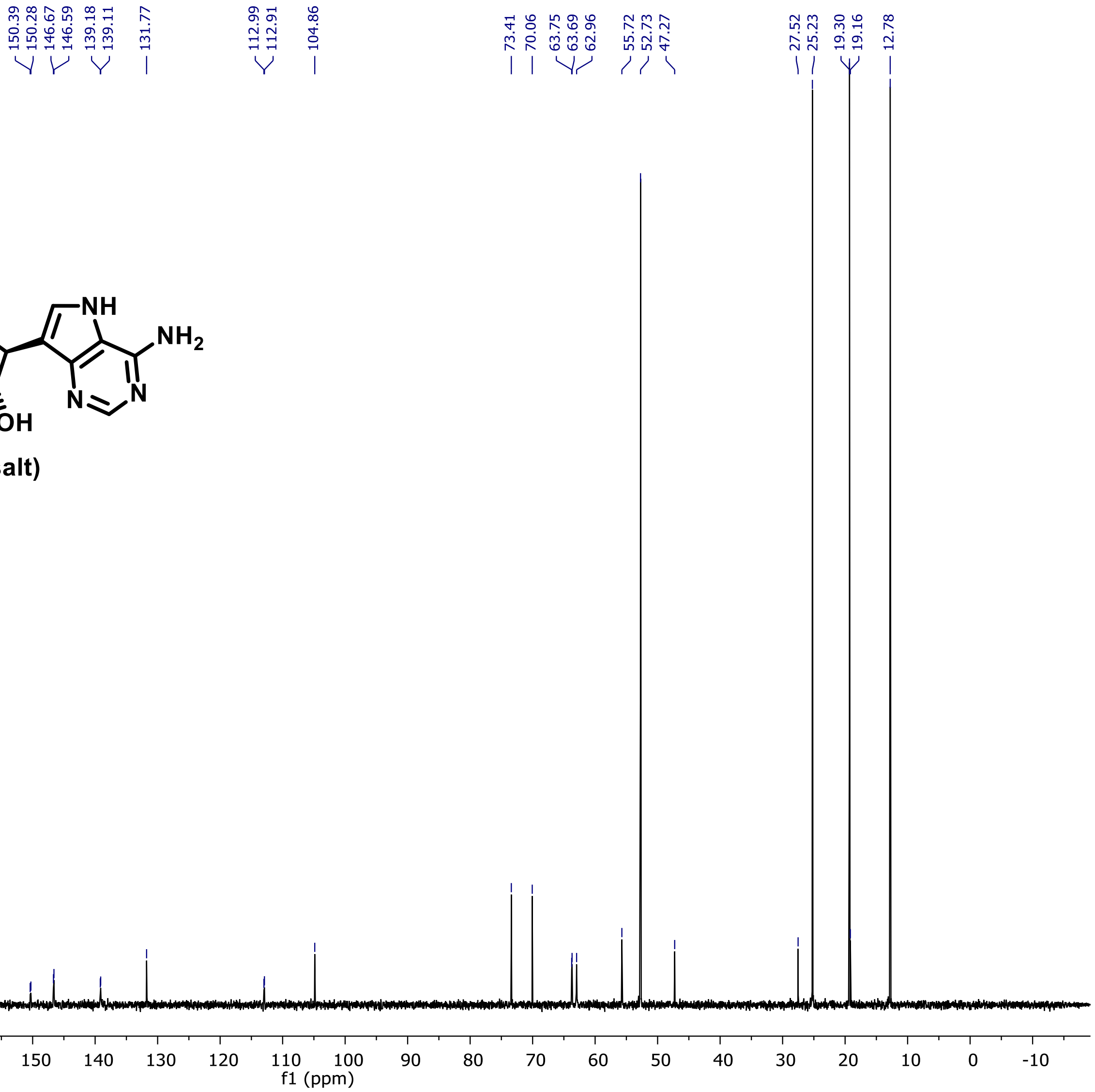

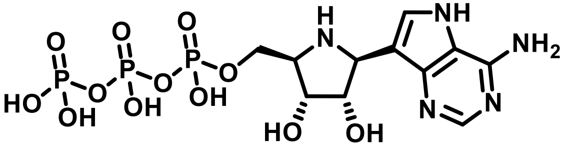

(2 $Bu_3N$ salt)

$^{31}P$ NMR (202 MHz, $D_2O$)

fe-RSh-AV20-080b.24.fid
Owner RSh
Sample fe-RSh-AV20-080b
Platform 3355
Job No fe

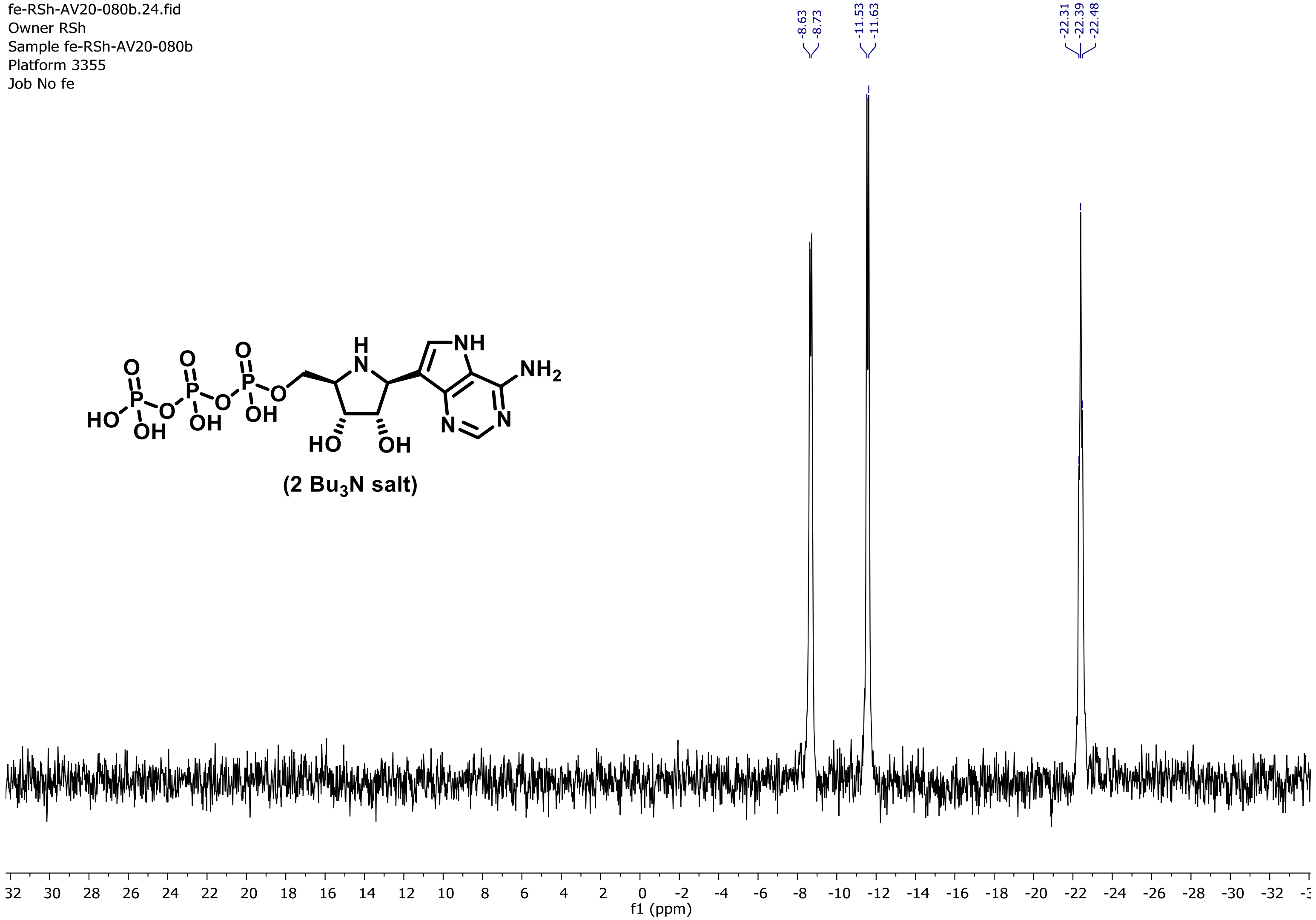

(2 $Bu_3N$ salt)

HRMS (ESI-MS)

### Elemental Composition Report

#### Single Mass Analysis

Tolerance = 5.0 mDa / DBE: min = -1.5, max = 150.0
Element prediction: Off
Number of isotope peaks used for i-FIT = 3

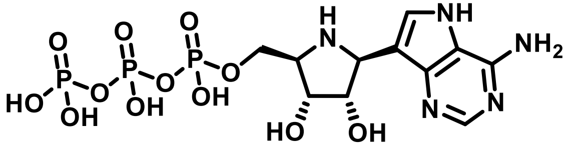

(2 $Bu_3N$ salt)

Monoisotopic Mass, Even Electron Ions
145 formula(e) evaluated with 2 results within limits (up to 100 closest results for each mass)
Elements Used:
C: 0-120 H: 0-150 N: 5-5 O: 0-40 Na: 0-1 P: 3-3

RSh-AV20-080b- 12 (0.143) Cm (9:19-1:7)

25-Aug-2020
1: TOF MS ES-
1.30e+008

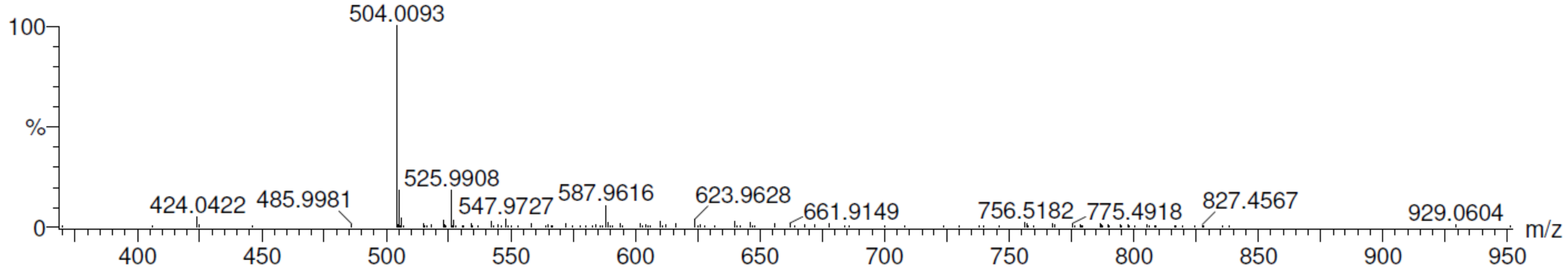

| | | | | |
|---|---|---|---|---|
| Minimum: | | | | -1.5 |
| Maximum: | | 5.0 | 5.0 | 150.0 |

| Mass | Calc. Mass | mDa | PPM | DBE | i-FIT | Norm | Conf(%) | Formula |
|---|---|---|---|---|---|---|---|---|
| 504.0093 | 504.0087 | 0.6 | 1.2 | 7.5 | 187.9 | 0.069 | 93.37 | C11 H17 N5 O12 P3 |
| | 504.0063 | 3.0 | 6.0 | 4.5 | 190.6 | 2.713 | 6.63 | C9 H18 N5 O12 Na P3 |
